# Discovery of orally bioavailable Zika virus NS2B-NS3 protease inhibitors with efficacy in a murine infection model

**DOI:** 10.1101/2025.06.19.660642

**Authors:** Nathaniel T. Kenton, Haim Barr, Artem Cherepakha, Lotte Coelmont, Caroline Collard, David Cousins, Geraint H. M. Davies, Oleksii Degtyarenko, Andrew Dirksen, Daniel Elbrecht, Daren Fearon, James Gayvert, Edward Griffen, Dirk Jochmans, Suzanne J. F. Kaptein, Pavlo Kliatskiy, Artem Kochetkov, Mykyta Kordubailo, Noa Lahav, Nir London, Elke Maas, Peter Marples, Nataliya Nady, Johan Neyts, Luong Nguyen, Xiaomin Ni, Yuliia Ogorodnik, Matthew C. Robinson, Carolina Turk Simpson, Tamas Szommer, Ihor Tarabara, Anton Tkachenko, Annette von Delft, Frank von Delft, Oleksandr Yakymenko, Alpha A. Lee

## Abstract

Flaviviruses are a class of pathogenic viruses with pandemic potential that are typically transmitted via infected arthropods. In addition, Zika virus is sexually transmissible and causes congenital malformations if infection occurs during pregnancy. Although over 1.5 million people were infected during the 2015-2016 Zika outbreak, to date, there are no clinical-stage vaccines or antivirals. Herein, we report the discovery of potent inhibitors of the Zika virus NS2B-NS3 protease that also show activity against the West Nile virus NS2B-NS3 protease. Starting from a crystallographic fragment screen, we employed a pharmacophore approach coupled with high-throughput library chemistry to elaborate fragments in the active site. Potent, metabolically stable, non-covalent, non-peptidomimetic inhibitors were identified with antiviral activity *in vitro*. The lead compound was progressed to a three-day Zika virus challenge study in AG129 mice, where it displayed significant viral RNA load reduction in both mouse plasma and spleen.

## Introduction

Zika virus (ZIKV) is an important pathogen of concern for global health. ZIKV infection is usually asymptomatic in healthy adults, but in pregnant women the virus can cross the placental barrier. Fetal infection can cause microcephaly and serious neurological damage, leading to lifelong disability^1^. During the 2015-2016 Zika epidemic in the western tropics, over 1.5 million people were infected, and an estimated 3500 children were born with disabilities^2,3^. Although also sexually transmissible^4^, ZIKV is mainly spread by mosquitoes of the genus *Aedes*^*5*^. Climate change increases the range of these mosquitoes^6^, compounding the pandemic risk from increased geographical distribution, increased infection, and increased risk of relevant mutants^6^. Other orthoflaviviruses transmitted by *Aedes*, such as dengue virus (DENV)^7^ and West Nile virus (WNV)^8^, already have a conspicuous impact even in more temperate climates.

There are no ZIKV-directed antivirals approved or in clinical trials. More broadly, across the orthoflavivirus genus, only DENV-specific antivirals have been advanced to clinical trials by Johnson & Johnson and Novartis, with both assets targeting the NS3–NS4B interaction^9,10^. Therefore, the identification of broad-spectrum small molecule inhibitors with complementary mechanisms of action is important for pandemic preparedness.

The ZIKV genome comprises a single-stranded, positive-sense RNA which encodes a polyprotein with ∼3400 amino acids. During viral replication, this polyprotein is processed into three structural proteins and seven nonstructural proteins (NS1, NS2A, NS2B, NS3, NS4A, NS4B and NS5). The NS2B-NS3 protein is a serine protease, responsible for making 5 polyprotein cleavages. While NS3 contains the active site and catalytic triad, it has limited activity without the NS2B protein, a membrane-bound scaffolding protein that reinforces the active conformation of NS3 via noncovalent interactions^11^. Across orthoflaviviruses, the NS2B-NS3 protease is highly conserved, corroborating its importance in the viral life cycle. Inhibition of the NS2B-NS3 protease is expected to block viral replication, a potentially useful therapy^12^.

Although protease inhibition is a precedented therapeutic mechanism for antivirals^13^, there is thus far a dearth of advanced chemical matter directed at ZIKV NS2B-NS3. Many efforts have focused on covalent inhibitors targeting the active site Ser135 of the ZIKV NS2B-NS3 protease, including several reports of boronate esters^14,15^, aldehydes^16^, or electrophilic esters^17^. However, covalent inhibitors can display promiscuous reactivity with human off-targets, thus posing a safety risk^18^. Among non-covalent small molecules, research efforts have primarily focused on polycationic and/or peptidomimetic compounds (representative structures shown in Figure 1)^19–21^. Many described inhibitors show excellent biochemical potency, but tend to have little to no cellular antiviral activity, limiting their therapeutic utility.

**Figure 1.**
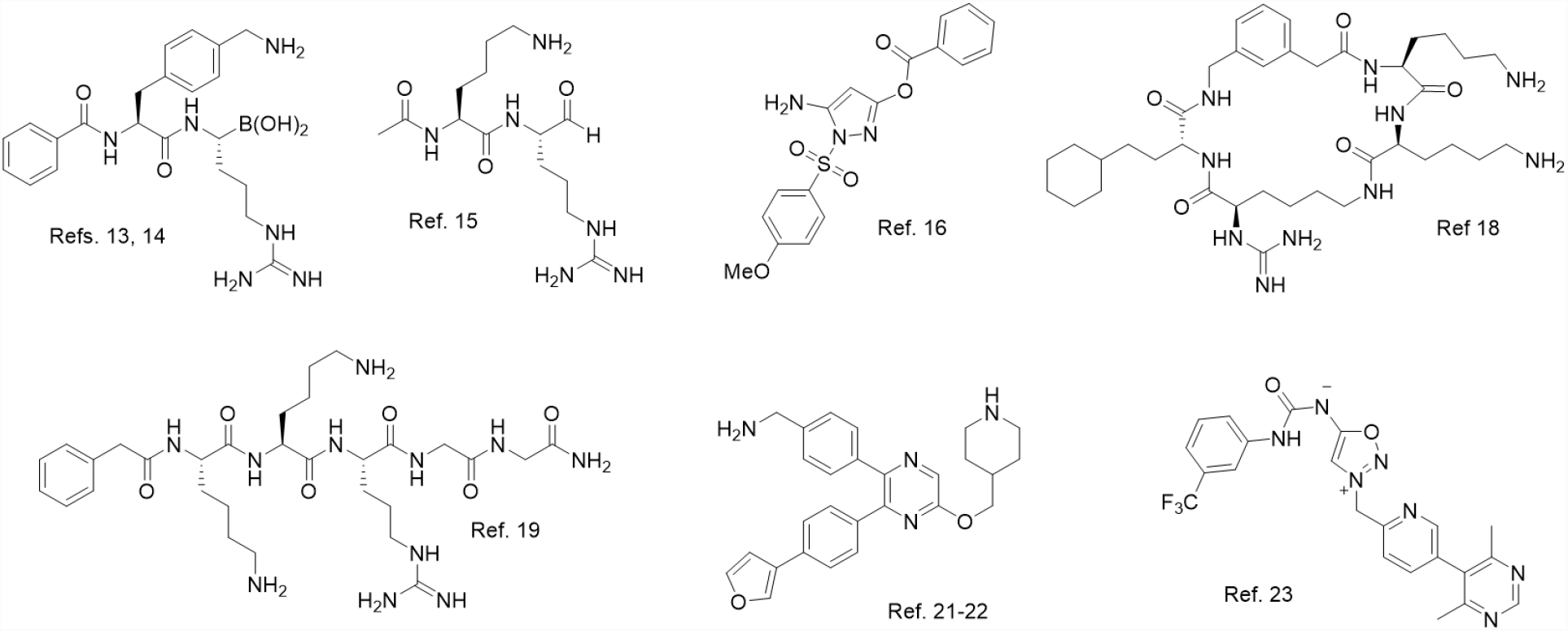
Representative ZIKV NS2B-NS3 inhibitors from the chemical literature. While many potent compounds have been reported, including some with broad-spectrum NS2B-NS3 inhibitory activity, most reported potent inhibitors are covalent, peptidomimetic, polycationic, or allosteric binders.

To our knowledge, the most advanced ZIKV NS2B-NS3 inhibitors reported to date bind an allosteric site of the protease; in fact, two independent groups have reportedly achieved murine efficacy with allosteric binding compounds^22–24^. The results of Ontoria et al. are particularly impressive: they report an unusual and orally bioavailable N-acyl-sydnone imine compound that, at 100 mg/kg twice-daily dosing (BID), suppressed viral RNA in plasma by 6 log_10_ and rescued the entire treatment group in a lethal 14-day murine challenge study^24^. This result not only serves as a powerful benchmark for progress of future ZIKV NS2B-NS3 inhibitors, but may spur interest more generally among medicinal chemists in sydnone derivatives and related compounds. However, targeting allosteric sites is a known risk in antiviral drug discovery, as resistance mutations are expected to develop more quickly^25^. Indeed, Ontoria et al. report the point mutation I156T on NS3 significantly impacts antiviral activity through resistance selection experiments.

Herein we report the fragment-based discovery of non-peptidomimetic, non-covalent inhibitors that bind the active site of the ZIKV NS2B-NS3 protease, filling an important gap in the field. Starting from a crystallographic fragment screen, we used a pharmacophore approach coupled with iterative high-throughput library chemistry to elaborate fragments whilst maintaining favorable physicochemical properties^26^. We were able to identify a lead compound and progress it to a murine model of ZIKV infection, where it displayed robust viral suppression in a three-day challenge study.

### Fragment based hit discovery with pharmacophore predictions

We began with a crystallographic fragment screen containing 1076 unique compounds directed at the co-expressed ZIKV NS2B-NS3 protease^27^. We focused our attention on the active site of the protein, since it is hypothesized that targeting the substrate envelope in the active site of viral enzymes confers robustness to resistance^25^. Although 46 fragments are bound near or within the active site, 34 of these were densely clustered in a deep pocket in the S1 site, sandwiched between Ala132 and Tyr161, and directly adjoining the His51/Asp75/Ser135 catalytic triad (Figure 2A). The majority (28/34) of these were aromatic compounds, supporting a notion of aromatic interactions with Tyr161. None of the fragments that were observed to bind the active site showed activity in our biochemical FRET assay up to 100 µM.

**Figure 2.**
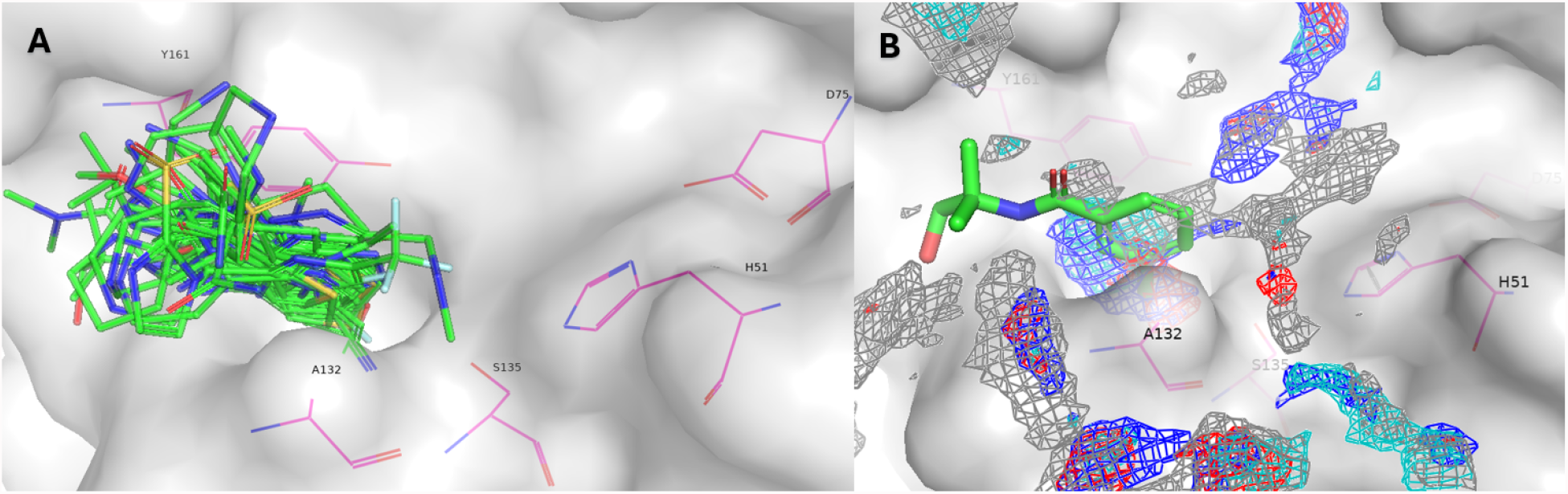
(A) Overlay of the fragments that bound the S1 site of the ZIKV NS2B-NS3 protease. The high aromatic ligandability of this site inspired compound design. (B) A machine-learning model was used to predict pharmacophore hotspots for many fragment-bound structures—the meta vector of benzamide fragment **Z68299550** (green, see also PDB ID: 7H1J) was considered an attractive growth opportunity based on this analysis (gray mesh = aromatic, blue mesh = HB donor, red mesh = HB acceptor, cyan mesh = pos ionizable). The NS2B-NS3 catalytic triad (His51, Asp75, Ser135) is shown in yellow.

An initial pharmacophore analysis of the active site utilized machine learning to predict locations of hydrogen bond donor, hydrogen bond acceptor, positive ionizable, and aromatic pharmacophores in the active site. These predictions were overlaid on the fragment hits, allowing us to select fragments with well-positioned vectors, such as **Z68299550** (Figure 2B, PDB ID: 7H1J), which were expanded to capture untapped interactions.

Starting from **Z68299550**, we employed machine-learning-based generative techniques for library generation at the desired vector, then scored the library based on pose alignment with the predicted pharmacophore fields^28^. These virtual libraries were triaged based on predicted binding pose with **Z68299550** as an anchor fragment. Figure 3B shows a predicted pose and potential interactions with three additional pharmacophores for one of our designs,**ASAP-0015373**, superimposed on fragment **Z68299550**. Figure 3C shows that our predicted binding mode qualitatively aligns with the solved structure. The binding mode of **ASAP-0015373**, when overlaid with the predicted pharmacophore fields around it, indicated rational avenues for optimization. In particular, aromatic and hydrogen bond donor pharmacophores were predicted, but not satisfied, by **ASAP-0015373** (Figure 3D). These predictions were leveraged to design another set of virtual libraries more closely related to **ASAP-0015373** in a second design cycle. **ASAP-0016806** emerged as a more potent inhibitor, which was crystallized with ZIKV NS2B-NS3 (Figure 3E).Comparison of the **ASAP-0015373** (PDB ID:9RM4) and **ASAP-0016806** (7I9J) crystal structures provided a structural hypothesis for improved potency (Figure 3D): a) replacement of the C-linked carbonyl of ‘373 with an NH-linked carbonyl of ‘806 satisfied the hydrogen-bond-donor pharmacophore that had been predicted for the former (dark blue arrow); b) reversal of the amide connecting the chlorophenyl group with the His51-targeted aromatic relaxed the linker and reduced the distance between His51 and the corresponding aromatic (5.0 Å for **ASAP-0015373** pyrrole vs. 3.7 Å for aniline of **ASAP-0016806**; light blue arrow); at the same time, a new hydrogen bond formed between the transposed secondary amide and Gly151, and the distance between the amine substituent and the Asp83 carboxylic acid group appeared to have closed (3.8 to 3.0 Å).

**Figure 3.**
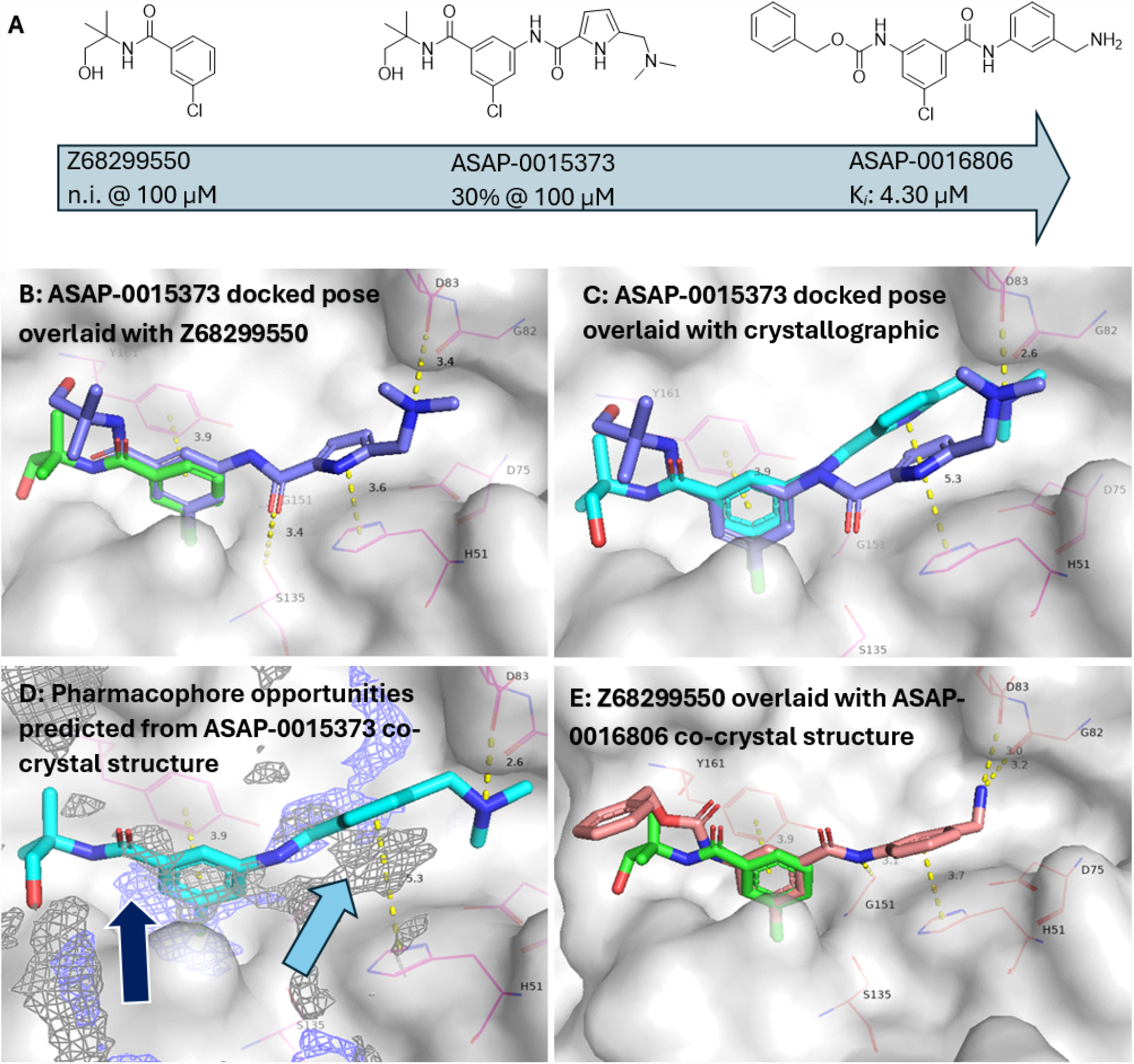
(A) Milestones in the advancement of fragment **Z68299550** (no activity in biochemical assay up to 100 µM) to inhibitor **ASAP-0016806**. (B) Overlay of fragment **Z68299550** (green) and **ASAP-0015373** docked pose (purple). (C) Comparison of the docked (purple) and crystallographic (cyan) poses of **ASAP-0015373**. (D) Predicted pharmacophore opportunities in the binding site (arrows) motivated introduction of a proximal hydrogen bond donor (blue mesh, dark blue arrow), and suggests an opportunity for an aromatic interaction above His51 (gray mesh, light blue arrow). (E) The crystal structure of **ASAP-0016806** (salmon, 7I9J) is shown overlaid with fragment **Z68299550** (green).

### Hit expansion with parallel library chemistry

We next turned to expand the hit **ASAP-0016806** to increase potency. The compound’s modular structure enabled a library design based on amide coupling to vary both sides of the molecule (Figure 4). Each library identified a breakthrough compound: isoindoline **ASAP-0020915** was found in Library 1 and indazole **ASAP-0023261** was found in Library 2. Both compounds markedly improved on both the ligand efficiency (defined here as the quotient of p*K*_*i*_ and total heavy count^29^) and absolute biochemical potency of the starting point **ASAP-0016806**. The privileged isoindoline and indazole groups featured in both molecules were then combined, affording a yet more potent inhibitor, **ASAP-0027808**.

**Figure 4.**
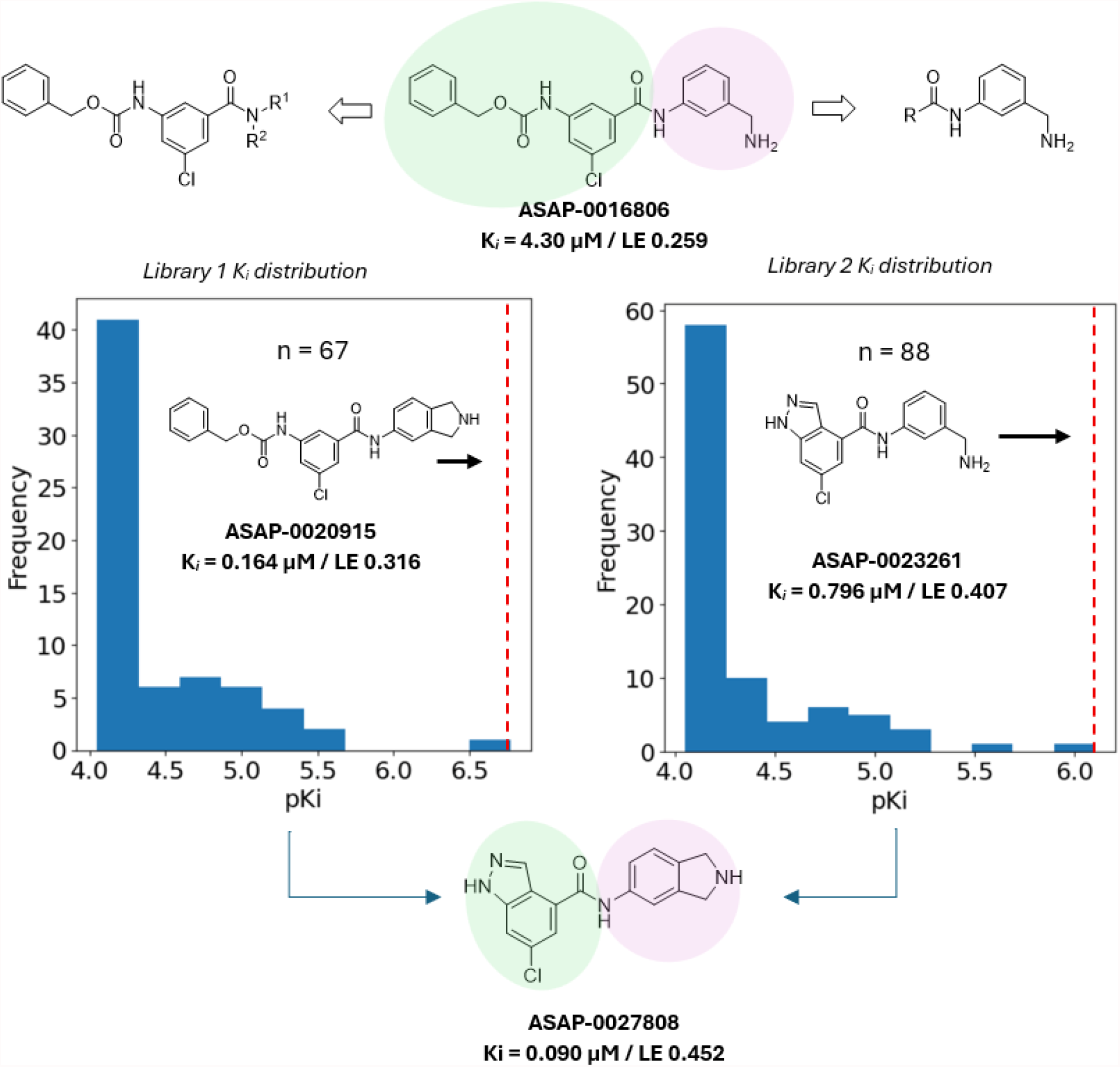
Independent libraries each maintaining one subunit of the **ASAP-0016806** structure were enumerated for hit expansion; for library 1 (left panel), the green-shaded region was kept constant; for library 2 (right panel), the pink-shaded region was kept constant. **ASAP-0020915** (library 1, red dotted line) and **ASAP-0023261** (library 2, red dotted line) both improved upon the ZIKV potency and ligand efficiency of the starting point. The privileged isoindoline and chloroindazole groups were then combined to generate **ASAP-0027808**, which showed a marked improvement in biochemical potency and ligand efficiency over the starting point, **ASAP-0016806**. (Ligand efficiency = p*K*_*i*_ / heavy atom count).

We profiled **ASAP-0027808** more extensively, noting ADME properties suitable for further development, excellent selectivity over 30 human proteases in a Eurofins selectivity panel (the highest inhibitory activity observed was 21% at 10 µM against urokinase, see Supporting Information, Section 6), and some activity against the WNV NS2B-NS3 protease (Figure 5A), suggesting that the chemotype might be developable toward useful pan-orthoflavivirus inhibitors. Upon crystallization in the active site of ZIKV NS2B-NS3, we noted the compound participating with most of the same residues as **ASAP-0016806** (Figure 5B): an aromatic interaction with Tyr161, a hydrogen-bonding interaction with Gly151, an aromatic interaction with His51, and an electrostatic interaction with Asp83 (with a possible additional hydrogen bonding interaction to Ser81). We observed a robust and dose-dependent antiviral effect against ZIKV in VeroE6 cells treated with a P-gp inhibitor, and no apparent cytotoxicity up to 50 µM (Figure 5C).

**Figure 5.**
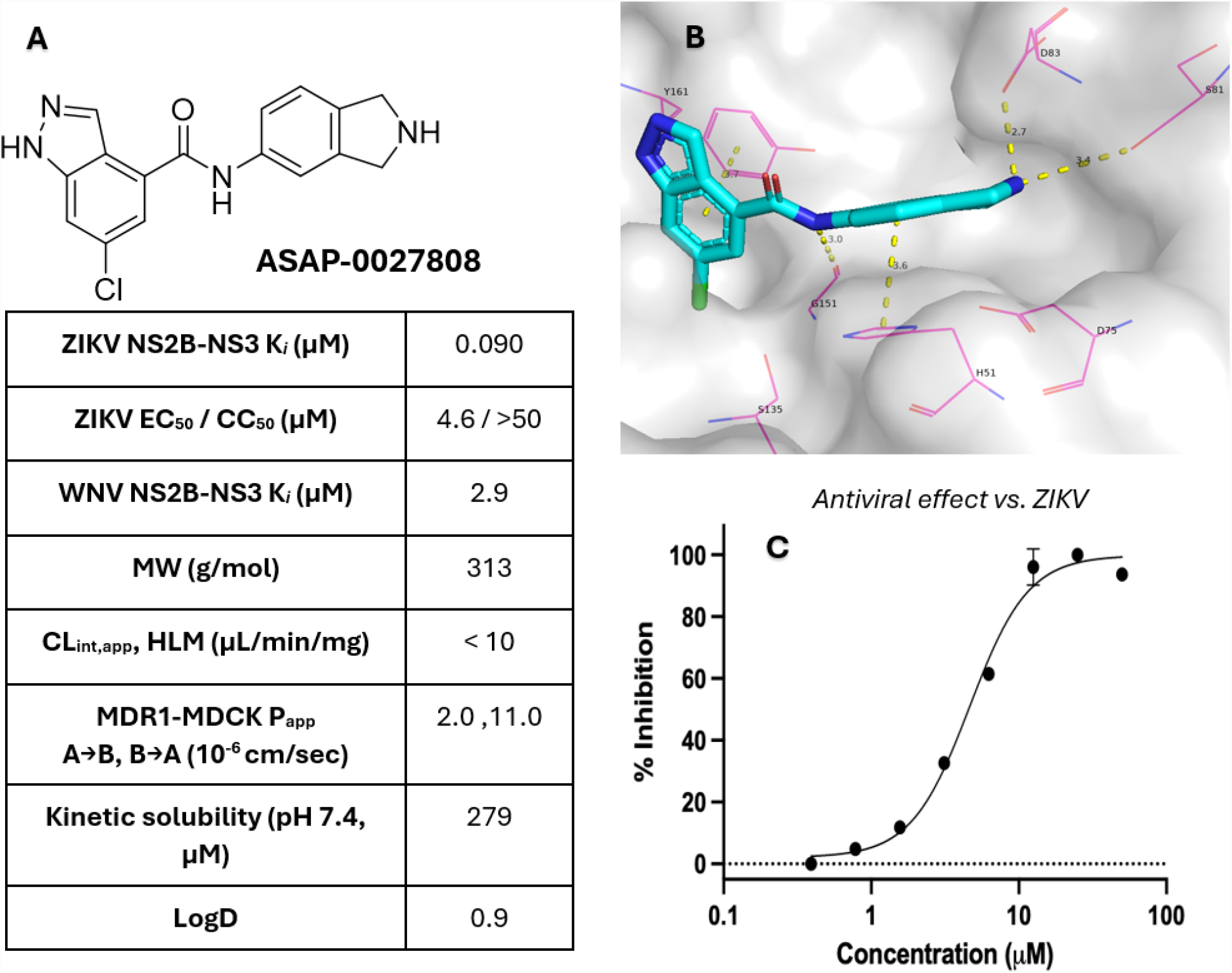
(A) Further characterization of **ASAP-0027808**. P_app_ was determined in MDR1-MDCK cells. Antiviral activity in VeroE6 cells as determined in a 7-day infection experiment using 0.5 uM CP100356 as a P-gp inhibitor (see Supporting Information for more details). (B) The compound was successfully crystallized with ZIKV NS2B-NS3. **ASAP-0016806** and **ASAP-0027808** (PDB ID: 9RM6) both appear to interact with the same residues (catalytic triad His51/Asp75/Ser135 colored yellow, other key residues Tyr161, NS2B Ser81, and NS2B Asp83 colored green). (C) Dose-dependent effect in a VeroE6 CPE-based assay (n=2). Cells were co-treated with 0.5 µM CP100356 (as P-gp inhibitor).

### Core hopping led to the discovery of lead compounds

Having established a connection between protease inhibition and cellular antiviral activity, we sought to further optimize potency by revisiting the predicted pharmacophores. Prediction of pharmacophores near the bound inhibitor revealed several opportunities for new polar interactions that might be captured with an sp^3^ hybridized linker rather than the amide linker (Figure 6A, yellow arrows). Thus we hypothesized that multi-component coupling reactions between Compound **1**, Compounds **2**/**3**, and various nucleophiles might furnish an sp^3^ hybridized group which could extend laterally to exploit the predicted pharmacophores.Compounds featuring nonpolar substituents such as **ASAP-0029525** and **ASAP-0029049** were less potent in the FRET assay, in line with our pharmacophore predictions. However, polar groups were tolerated, and in particular **ASAP-0029000**, derived from isopropyl isocyanide via an Ugi reaction^30^, showed the success of this pharmacophore-driven approach: compared to the starting point **ASAP-0027808**, it was equipotent against ZIKV NS2B-NS3 and 10-fold more potent against the WNV protease. Upon chiral separation, the more potent enantiomer, **ASAP-0029974**, showed similar biochemical and antiviral potencies as **ASAP-0027808** (Figure 6B).

**Figure 6.**
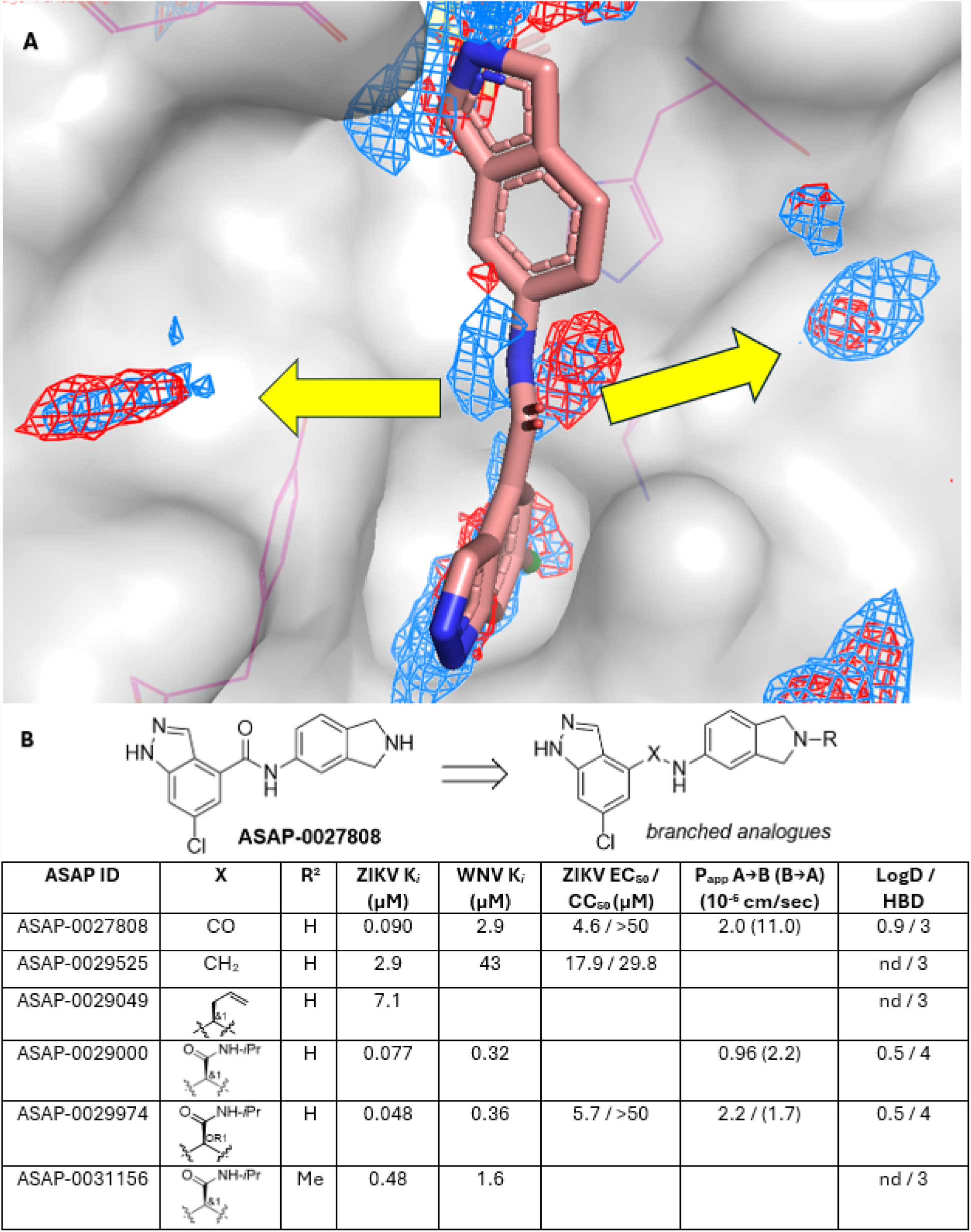
Exploring new interactions via multicomponent reactions, guided by pharmacophore predictions. (A) **ASAP-0027808** crystal structure with ZIKV NS2B-NS3 overlaid with key ML-predicted pharmacophores (red mesh = hydrogen bond acceptor; blue mesh = hydrogen bond donor). sp^2^ hybridization of the linker prevents lateral ligand growth to capture these interactions (yellow arrows). (B) sp3-hybridized analogues of ASAP-0027808 were used to investigate the pharmacophore predictions.

**ASAP-0029000** introduced a fourth hydrogen bond donor, threatening to push any follow-up compounds in the chemotype further from developable physicochemical space, especially with respect to membrane permeability^31^. Analyzing pharmacophores predicted from the **ASAP-0029000** structure (Figure 7A, PDB ID: 7I9R), we noticed that regions defining hydrogen-bond-donor and hydrophobic pharmacophore fields overlapped near the indazole NH. This suggested an opportunity to tune the physicochemical properties of the series toward more permeable and lipophilic compounds without compromising antiviral activity. A library based on the three-component Ugi reaction (Figure 7B) was designed to expand upon **ASAP-0029000** in which one of the primary goals was to validate this pharmacophore hypothesis. The most potent compounds from this library were also in more desirable physicochemical space, which together contributed to greater antiviral potency. Of the compounds in the library, **ASAP-0031651** was the most potent antiviral (*K*_*i*_ = 7.8 nM; VeroE6 EC_50_ = 0.45 µM).

**Figure 7.**
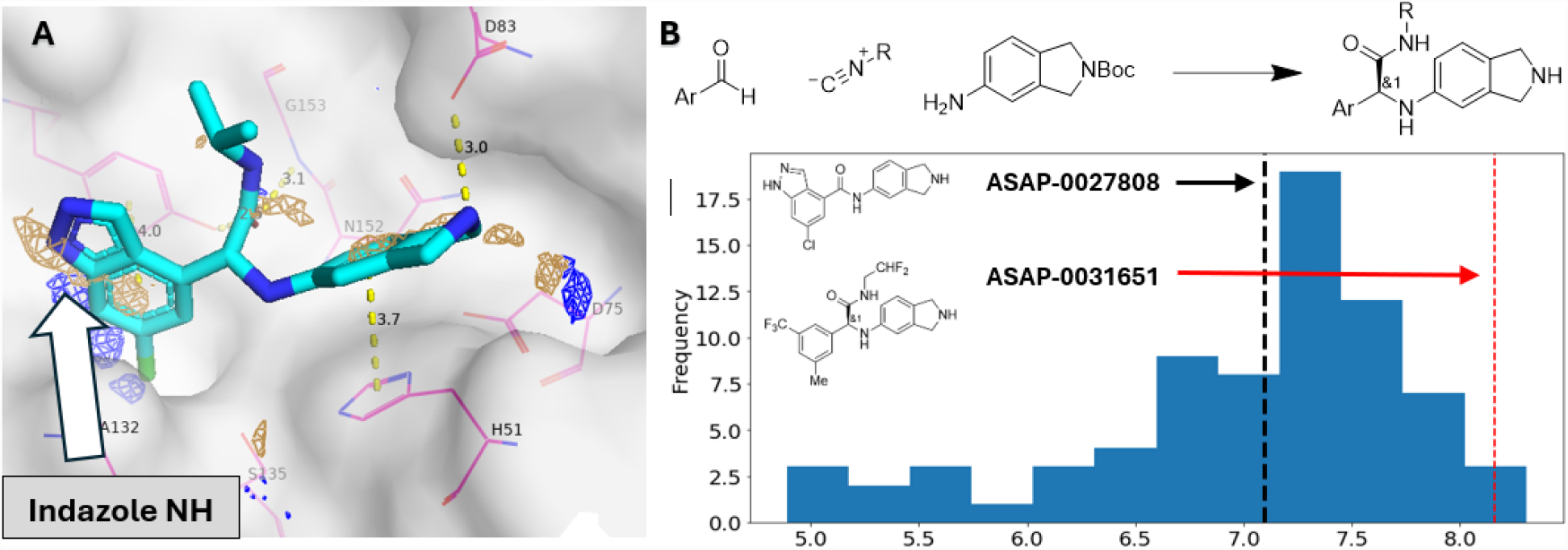
Optimization of the Ugi chemotype via library chemistry. (A) Overlapping hydrogen bond donor (blue) and hydrophobic (tan) pharmacophores were predicted at the indazole NH (white arrow), suggesting an opportunity to improve permeability by replacing a hydrogen bond donor with a lipophilic group (pdb ID: 7I9R). (B) A library that varied both isocyanide and aldehyde components was designed to explore the SAR. The isoindoline group was kept constant. Results for the Ugi library; progression from lead benzamide **ASAP-0027808** (black line) to top hit **ASAP-0031651** (black dotted line), is indicated.

To learn more about the new Ugi-derived chemotype, chiral separations were performed on many of the most potent compounds from the library. Of note, **ASAP-0036543**, the most potent of the two enantiomers of **ASAP-0031651**, achieved high biochemical potency against both ZIKV and WNV NS2B-NS3 proteases, alongside high kinetic solubility, low microsomal clearance, and acceptable permeability in an MDR1-MDCK cell line (Figure 8A). Its respectable antiviral potency against WNV in VeroE6 cells was also noted, which had been hitherto difficult to achieve with the chemotype. Most excitingly, we observed potent antiviral activity against ZIKV in three different cell lines, with close alignment of the observed EC_50,u_ (Fig. 8A-B). VeroE6 cells were used as the standard cell line in the ZIKV antiviral and cell viability assays, whilst the other 2 cell lines represent more relevant human cell types for ZIKV replication (JEG-3 - placental cell line, SH-SY5Y - neuroblastoma cell line). **ASAP-0036543** crystallized with ZIKV NS2B-NS3 (Figure 8C) and displayed a similar binding mode to **ASAP-0029000**, with the aromatic sitting next to Tyr161, the amide carbonyl oxygen atom coordinated in a possible bidentate hydrogen-bonding pattern with Gly153 and Tyr161, and the isoindoline making key interactions with His51 (aromatic) and Asp83 (electrostatic). We noted that ASAP-0036543 is a much weaker inhibitor of DENV-2 NS2B-NS3 than of the ZIKV and WNV proteases. The dramatic shift may be driven in part by a key residue difference at NS2B position 83: whereas ZIKV and WNV NS2Bs have acidic residues (Asp83 and Glu83, respectively) that likely form electrostatic interactions with the isoindoline group, DENV-2 NS2B has an uncharged threonine instead.

**Figure 8.**
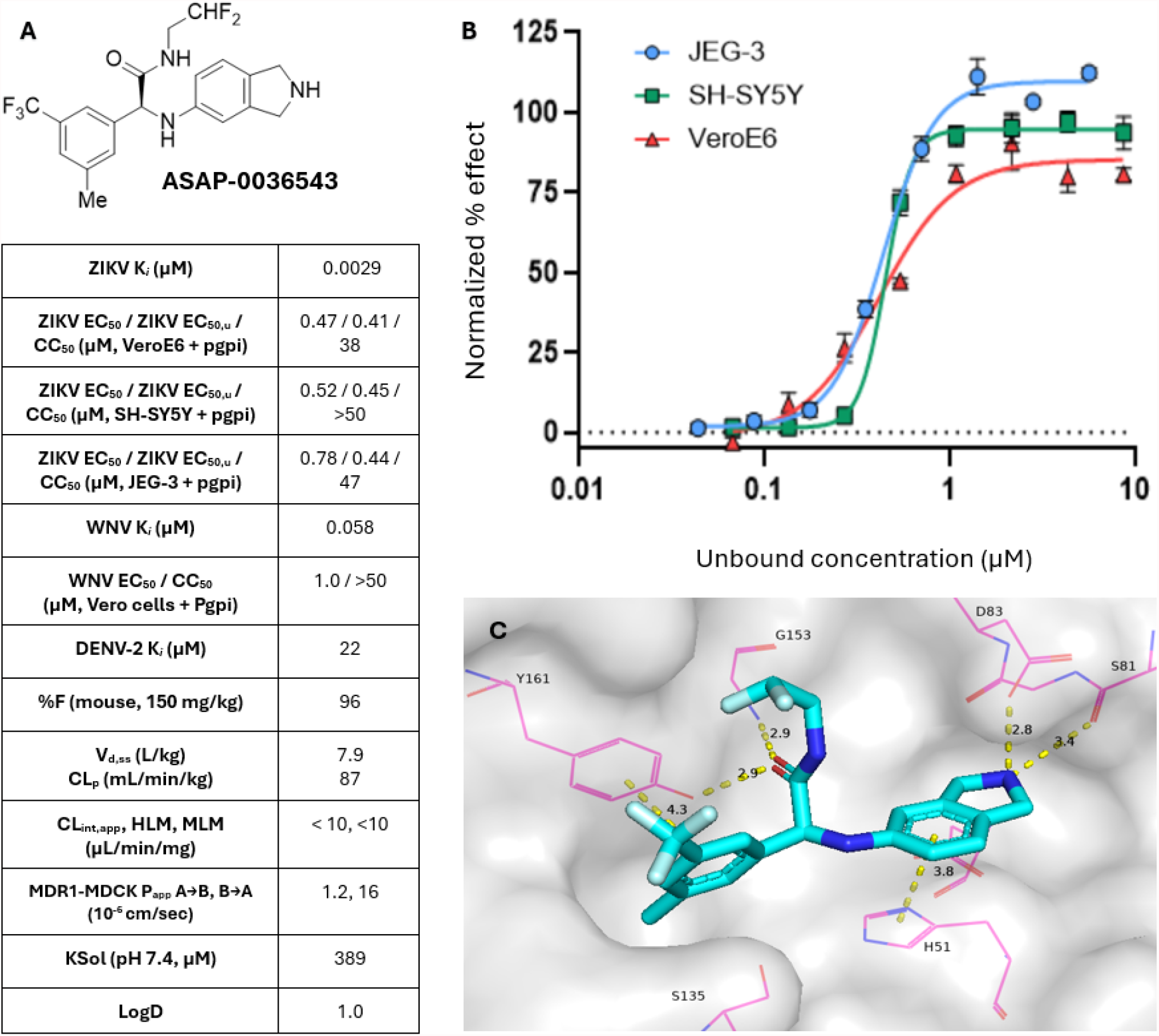
(A) Full profile of lead compound **ASAP-0036543**, the more active enantiomer of **ASAP-0031651**. (B) Antiviral data for ASAP-0036543 across three cell lines (cell viability measured by MTS, normalized to infected and uninfected controls). For VeroE6 + SH-SY5Y cells, the assay contains 2% FCS; for JEG-3, the assay contains 10% FCS (see Supporting Information section 5 for more information regarding unbound calculations). (C) **ASAP-0036543** was assigned the (*S*) configuration by crystallography (pdb ID 9RM7) and the compound assumed a similar binding mode to that observed with **ASAP-0029000**.

### Efficacy in a ZIKV challenge study in AG129 mice

**ASAP-0036543** was advanced to murine pharmacokinetic studies to assess its potential as an *in vivo* tool compound^32^. The compound was well-tolerated when dosed PO at 150 mg/kg, and free plasma exposure above unbound EC_50_ was achieved for almost 8 h (Figure 9A)^33,34^. In light of these results, **ASAP-0036543** was advanced to a ZIKV challenge study in AG129 mice^32,35^.

**Figure 9.**
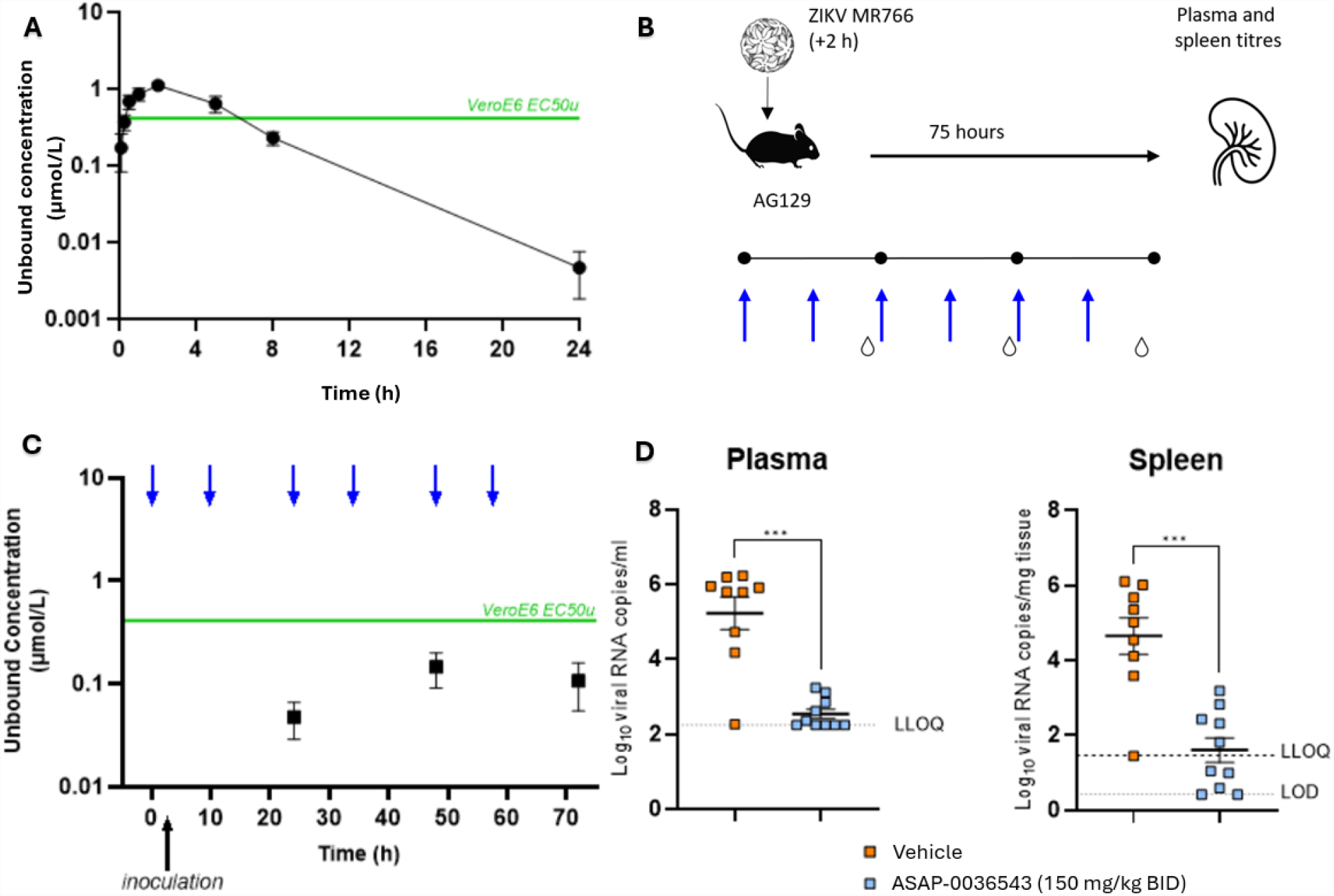
(A) Murine *in vivo* pharmacokinetics at 150 mg/kg (PO solution dosing, three animals). Free exposure over unbound EC_50_ in VeroE6 cells was observed for approximately 6 hours. (B) Study design: groups of 10 mixed sex AG129 mice were treated with vehicle-only or **ASAP-0036543** at study start (PO, 150 mg/kg, BID), followed by inoculation with ZIKV MR766 2 hours later. Additional doses of **ASAP-0036543** or vehicle only were given BID over the course of 3 days (blue arrows). ASAP-0036543 was quantitated in plasma at C_min_ following doses 2, 4, and 6. (C) PK data from the murine challenge study; male mice only. (D) Viral RNA load reduction was observed in plasma and spleen samples harvested at the end of the study (T+75 h, p < 0.001).

With a potential tool molecule in hand, we sought to achieve antiviral proof-of-concept in a previously-described ZIKV challenge study in AG129 mice^32^. Based on the PK study, we dosed orally at 150 mg/kg BID for 3 days. The compound was not tolerated at higher dose levels. During the course of the efficacy study, concentration of the compound was determined thrice at C_min_ following doses 2, 4, and 6 (at T+24, 48, and 72h of study; black boxes, Fig. 9B). We observed significant viral RNA load reduction after only three days in both plasma and spleen (p < 0.001), with viral RNA levels in many samples below the lower level of quantitation (LLOQ, Fig. 9C). With such robust efficacy, it is interesting that full coverage over EC_50_ was not achieved during the study, although it is likely that C_max_ was above unbound EC_50_ (see Fig. 9A). This is an encouraging result for future lead optimization efforts targeting the NS2B-NS3 protease.

## Conclusions

Novel, broad-spectrum orthoflavivirus therapeutics are urgently needed to combat the rising incidence of flavivirus infections world-wide. Of particular concern is the paucity of reported flavivirus inhibitors with Zika virus efficacy. Here, we describe a structure-guided fragment-based lead discovery campaign against the ZIKV NS2B-NS3 protease, identifying potent compounds active against both ZIKV and WNV, as shown in biochemical and cell-based assays. We used pharmacophores predicted by machine-learning to inform both a key scaffold hop and an opportunity to rationally tune physicochemical properties. High-throughput Ugi-based chemistry was used to identify key hit **ASAP-0031651**, which afforded our lead compound, **ASAP-0036543**, upon chiral separation.

It is noteworthy that, as the chemotype matured from the first potent hit to the lead compound, there was no commensurate increase in synthetic complexity. Both **ASAP-0016806** and **ASAP-0036543** were synthesized in two simple synthetic steps from commercially available building blocks. We hope the simple structure of the chemotype and the study design disclosed herein facilitates future efforts in lead optimization for this important viral threat. Using this compound, we observed significant viral load reduction in ZIKV-infected AG129 mice relative to vehicle-treated control animals after 3 days. We hope that our work may be the first step towards a pan-flavivirus NS2B-NS3 inhibitor, with DENV spectrum being a key endpoint to optimise.

## Methods

### Cell lines and culture conditions

SH-SY5Y and Vero E6 cells were maintained in Dulbecco’s Modified Eagle Medium (DMEM, Gibco cat. no. 41965-039) supplemented with 10% heat-inactivated fetal calf serum. (Vero E6 cultures additionally contained 1% sodium bicarbonate.) JEG-3 cells were cultured in Minimum Essential Medium (MEM, Gibco cat. no. 25080-060) supplemented with 10% heat-inactivated FCS, 1% glutamine, 1% non-essential amino acids (NEAA, Gibco cat. no. 11140-035), 1% HEPES, and 1% sodium pyruvate. Cells were cultured at 37°C with 5% CO_2_ and passaged at a 1:10 ratio once weekly. For passaging, confluent cultures were washed with Dulbecco’s phosphate-buffered saline (DBPS), treated with 0.25% trypsin-EDTA, incubated for 5 min (15 min for Vero E6 cells) at 37°C (room temperature for JEG-3 cells), and resuspended in 10 mL growth medium. 1 mL of the cell suspension was transferred into new T150 flasks or bottles containing 25 mL fresh growth medium.

### In vitro antiviral assays: ZIKV

Zika virus antiviral activity was evaluated in SH-SY5Y, Vero E6, and JEG-3 cells. Assays for SH-SY5Y and Vero E6 cells employed MEM, 2% v/v heat-inactivated FCS, 1% sodium bicarbonate, and 1% glutamine. For JEG-3 cells, assays were performed in the culture medium described above. Cell densities were determined using a Coulter counter. Cells were diluted to 10,000 cells/100 µL (SH-SY5Y), 10,000 cells/50 µL (Vero E6), or 3,000 cells/100 µL (JEG-3) in the assay medium. CP100356 was added to the cell suspension at 1 µM (SH-SY5Y and JEG-3; dilution 1:2000 from a 2 mM stock) or 2 µM (Vero E6; dilution 1:1000 from a 2 mM stock). For SH-SY5Y and JEG-3 assays, 100 µL of the cell suspension was seeded into each well of a 96-well Falcon plate and incubated overnight at 37°C and 5% CO_2_ prior to compound addition.For Vero E6 assays, 100 µL assay medium was first added to each well before compound and virus addition, followed by addition of 50 µL cell suspension.

Compounds were serially diluted 1:3 across columns 2–9 of the plate. Briefly, 50 µL medium was added to column 2, followed by addition of 3 µL compound stock solution (10 mM) to achieve a starting concentration of 100 µM, or 3 µL of a 1:10 predilution to achieve a starting concentration of 10 µM. Serial dilutions were performed by transferring 50 µL between consecutive columns with tip changes in columns 4 and 7. ZIKV MR766 virus stocks were diluted in assay medium to the appropriate working concentration (SH-SY5Y and JEG-3: stock titer 2.6 × 10^7^ PFU/mL, diluted 1:400 for a final dilution of 1:1600; Vero E6: stock titer 1.98 × 10^5^ PFU/mL, diluted 1:100 for a final dilution of 1:400 corresponding to MOI 0.01). Virus suspension (50 µL) was added to columns 2–10 (or columns 1–10 for Vero E6 assays), followed by addition of assay medium or cell suspension as appropriate. Final assay volumes resulted in a CP100356 concentration of 0.5 µM. Plates were incubated at 37°C and 5% CO_2_, and cell viability was determined using MTS^36^ on day 7 post-infection (SH-SY5Y and Vero E6 cells) or day 4 post-infection (JEG-3 cells).

#### In vitro antiviral assays: WNV

The assay medium consisted of MEM (Gibco cat. no. 21090-022) supplemented with 2% heat-inactivated FCS, 1% sodium bicarbonate, 1% glutamine, and 1% non-essential amino acids. Confluent Vero E6 cultures were washed with DPBS, detached with 0.25% trypsin-EDTA as described above, and resuspended in the assay medium. Cell density was determined using a Coulter counter, after which cells were diluted to 10,000 cells per 100 µL in the assay medium. CP100356 was added to the cell suspension at a concentration of 1 µM (prepared from a 2 mM stock at a 1:2000 dilution). 100 µL of the cell suspension was seeded into each well of a 96-well Falcon plate and incubated overnight at 37°C and 5% CO_2_.

Compounds were serially diluted 1:3 across columns 2–9 of the assay plate. Briefly, 50 µL medium was added to column 2, followed by addition of 3 µL of a 10 mM compound stock solution to achieve a starting concentration of 100 µM, or 3 µL of a 1:10 predilution to achieve a starting concentration of 10 µM. Serial dilutions were performed by transferring 50 µL sequentially between columns with tip changes in columns 4 and 7. WNV NY99 virus stock (8.89 × 10^7^ TCID_50_/mL) was diluted 1:125 in assay medium to obtain a final assay dilution of 1:500. 50 µL of virus suspension was added to columns 2–10, followed by addition of 50 µL assay medium to columns 2–11. Final assay conditions consisted of 100 µL cells containing 1 µM CP100356, 50 µL compound dilution, and 50 µL virus suspension, resulting in a final CP100356 concentration of 0.5 µM. Where necessary, border wells were filled with medium instead of cells. Plates were incubated at 37°C and 5% CO_2_, and cell viability was determined on day 7 post-infection using MTS^36^.

### Murine ZIKV challenge study: experiment setup

ASAP-0035643 was formulated in 10% Solutol prepared in 50 mM citrate buffer (pH 3) for oral administration at a dose of 150 mg/kg. The compound was administered twice daily (BID), and fresh formulations were prepared immediately prior to each dosing.

Male and female AG129 mice (129/Sv background deficient in both IFN-α/β and IFN-γ receptors), aged 7–15 weeks and weighing 20–25 g, were bred in-house from breeding pairs originally obtained from Marshall BioResources. The specific pathogen-free status of the animals was routinely monitored at the KU Leuven Rega Institute animal facility. Mice were housed in individually ventilated isolator cages (IsoCage N Biocontainment System, Techniplast) under controlled environmental conditions (21°C, 55% humidity, 12 h light/dark cycle) with ad libitum access to food and water. Environmental enrichment was provided using cotton nesting material and cardboard tunnels. All housing conditions and experimental procedures were approved by the KU Leuven Ethical Committee for Animal Experimentation (License DMIT-110/2025) in accordance with institutional guidelines and standards of the Federation of European Laboratory Animal Science Associations. All animal experiments were conducted under Biosafety Level 2 conditions.

Experimental groups consisted of five male and five female mice per study arm. At study initiation, mice assigned to the treatment group received the first dose of ASAP-0035643. Two hours later, all animals were inoculated intraperitoneally (200 μL) with 1 × 10^4^ PFU of Zika virus (ZIKV strain MR766). Animals were monitored daily for changes in body weight and clinical signs of disease. Mice exhibiting ≥20% body weight loss and/or severe disease symptoms, or those reaching the study endpoint, were euthanized by intraperitoneal administration of 50 μL Dolethal.

### Blood draws and PK quantitation

During the in vivo efficacy study, blood samples were collected from the tail veins of male mice at 24 h intervals into K_2_EDTA-coated tubes. Approximately 20 μL plasma was obtained for pharmacokinetic (PK) analysis, heat-inactivated for 30 min at 56°C, and stored at −80°C. Plasma samples were collected at 12 h (immediately prior to the third dose), 48 h (immediately prior to the fifth dose), and 72 h (3 h before study termination). A 30% isopropanol solution in water was used as the homogenization buffer.

Samples were stored at −80°C until shipment to Frontage Laboratories (Exton, PA, USA) for quantitation of ASAP-0036543 in mouse plasma. For LC–MS/MS analysis, 10 μL of control matrix was dispensed into separate wells of a 96-well plate for preparation of standards (STDs), quality controls (QCs), and blanks. For study samples, 10 μL of plasma sample and 10 μL diluent were added to separate wells. Protein precipitation was initiated by adding 200 μL internal standard solution containing 200 ng/mL warfarin and verapamil in acetonitrile to all wells. Plates were shaken for 20 min using a Qiagen TissueLyser II at 15 Hz and centrifuged at 4000 rpm for 10 min at 15°C. Following centrifugation, 100 μL supernatant was transferred to a fresh 96-well plate containing 100 μL water per well. The plate was vortexed for approximately 10 min prior to LC–MS/MS injection.

### RNA isolation and quantitative RT-PCR

RNA was isolated from 75 μl plasma using the NucleoSpin RNA virus kit (Filter Service, Germany). LightCycler96 (Roche) was used for RT-qPCR. The ZIKV NS1 region (nucleotides 2472–2565) was amplified using primers 5’-TGA CTC CCC TCG TAG ACT G-3’ and 3’-CTC TCC TTC CAC TGA TTT CCA C-5’ and a double-quenched probe 5’-6-FAM/AGA TCC CAC /ZEN/AAA TCC CCT CTT CCC/3’IABkFQ/ (Integrated DNA Technologies, IDT). Viral RNA copy numbers were determined using a standard curve generated from serial dilutions of a synthetic gene block containing 145 bp of ZIKV NS1 (nucleotides 2456–2603): 5’-GGT ACA AGT ACC ATC CTG ACT CCC CTC GTA GAC TGG CAG CAG CCG TTA AGC AAG CTT GGG AAG AGG GGA TTT GTG GGA TCT CCT CTG TTT CTA GAA TGG AAA ACA TAA TGT GGA AAT CAG TGG AAG GAG AGC TCA ATG CAA TCC TAG-3’ (Integrated DNA Technologies).

## Acknowledgment

Research reported here, excluding any in vivo studies, was supported in part by NIAID of the National Institutes of Health under award number U19AI171399. The content is solely the responsibility of the authors and does not necessarily represent the official views of the National Institutes of Health. Further funding was provided by the FWO Hercules Foundation (grant number ZW13-02, Caps-It infrastructure). We thank Elke Maas, Joost Schepers, Thibault Francken, and Winston Chiu for their excellent technical support on HCI screening, and Maïlis Darmuzey, Niels Cremers, Stijn Hendrickx, Carolien De Keyzer and Lindsey Bervoets for their excellent help with the in vivo work. The dataset has been deposited in ChEMBL (accession: ChEMBL_v36) and is publicly available^37,38^.

## Author contributions

All authors contributed to the conception and design of the studies. N.T.K. and A.A.L. wrote the manuscript. All authors read and approved the submitted version.

