## Supplementary material for "Discovery of orally bioavailable Zika virus NS2B-NS3 protease inhibitors with efficacy in a murine infection model": PDF of supporting information

#### Contents

1. Analytical methods for organic synthesis
2. Synthetic chemistry
3. NMR spectra for key compounds
4. Supplementary tables for cocrystal structures
5. FRET assay protocols for inhibition of ZIKV, WNV, DENV-2 NS2B-NS3 proteases
6. Antiviral assay protocols for ZIKV, WNV
7. In vitro ADMET protocols
8. Human protease selectivity for ASAP-0027808
9. Murine pharmacokinetics protocols
10. In vivo efficacy study
11. Safety profiling of ASAP-0036543
12. References

#### 1. ANALYTICAL METHODS

NMR spectroscopy was performed on Agilent ProPulse 600, Bruker AVANCE DRX 500, and Varian UNITYplus 400 instruments, and spectra were recorded at 600, 500, or 400 MHz, respectively. Chemical shifts (δ) are given in parts per million (ppm) and are listed upfield with tetramethylsilane as a reference. Peaks are described as singlets (s), doublets (d), triplets (t), quartets (q), quintets (quint) multiplets (m), and broad (br).

LCMS was performed using an Agilent Poroshell 120 SB-C18 4.6x30mm 2.7 µm column maintained at 60 °C. The default method "method A" was used unless otherwise specified: mobile phase A (water + 0.1% formic acid) and B (acetonitrile + 0.1% formic acid), flow rate: 3 ml/min, gradient: 0.01 min – 1% B, 1.5 min – 100% B, 1.73 min – 100% B. Electrospray ionization (ESI). UV detection was performed at 215 nm, 254nm and 280 nm.

In some cases, "method B" was used:

| **Ionization mode** | **DAD** | **Scan range** | | **Injection volume** | **Flow rate** |
|---|---|---|---|---|---|
| Electrospray ionization (ESI) | 215 nm, 254 nm, 280 nm | m\z 83-1000 | | 0.5 µl | 1.5 ml\min |
| **Instruments** | **Instument specifications** | **Column** | **Mobile phase** | **Gradient** | |
| 36 | Agilent Technologies 1290 Infinity LC/MSD system with DAD/ ELSD Agilent and Agilent LC/MSD G6125B mas-spectrometer | Poroshell 120 SB-C18 4.6x30mm 2.7 Micron | A-acetonitrile:water 95:5 (0.1% formic acid) B-water (0.1% formic acid) | 0.36 - 1/99 (A/B) 5.36 - 100 (A) 6.49 - 100 (A) | |

Details for analytical SFC separations are noted where appropriate.

#### 2. SYNTHETIC CHEMISTRY

**Method 1:**
**N-[3-chloro-5-[(2-hydroxy-1,1-dimethyl-ethyl)carbamoyl]phenyl]-5-[(dimethylamino)methyl]-1H-pyrrole-2-carboxamide (ASAP-0015373)**

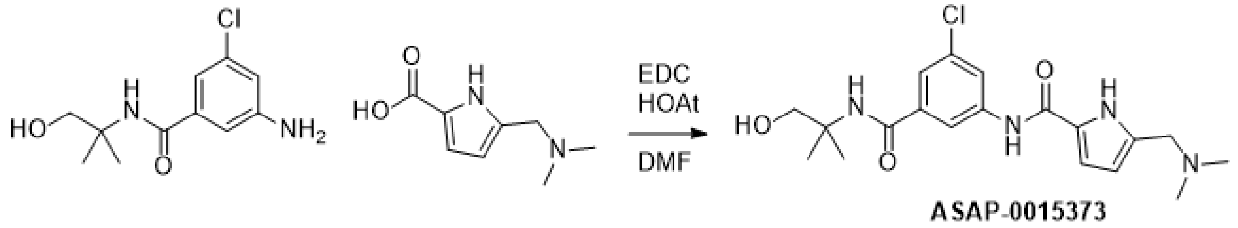

To a stirred solution of 3-amino-5-chloro-N-(2-hydroxy-1,1-dimethyl-ethyl)benzamide (Enamine ID: CSMB00006368432, 0.22 g, 0.091 mmol), 5-[(dimethylamino)methyl]-1H-pyrrole-2-carboxylic acid (Enamine ID: EN300-1588741, 0.16 g, 0.95 mmol), EDC (246 mg, 1.59 mmol), HOAt (174 mg, 1.28 mmol), and diisopropylethylamine (387 mg, 3 mmol) were mixed in DMF (2 mL). The resulting reaction mixture was stirred at rt for 4 d. The obtained crude product was purified by reverse phase HPLC to afford N-[3-chloro-5-[(2-hydroxy-1,1-dimethyl-ethyl)carbamoyl]phenyl]-5-[(dimethylamino)methyl]-1H-pyrrole-2-carboxamide (ASAP-0015373) (19.8 mg, 7.77 umol, 9% yield) as a white solid.

LCMS(ESI): $[M+H]^+$ m/z: calcd 393.2; found 393.2 ; Rt = 0.897 min.
$^1$H NMR (500 MHz, Acetonitrile-d3) δ (ppm): 1.38 (s, 6H), 2.77 (s, 6H), 3.62 (s, 2H), 4.29 (s, 2H), 6.43 (d, 1H), 6.80 (br s, 1H), 6.98 (d, 1H), 7.50 (s, 1H), 8.06 (s, 1H), 8.10 (s, 1H), 9.21 (s, 1H), 11.94 (br s, 1H). $^{13}$C NMR (125 MHz, DMSO-d6) δ (ppm): 23.94, 45.09, 55.49, 55.63, 67.58, 109.39, 113.08, 117.85, 121.29, 121.48, 125.15, 133.16, 135.05, 138.46, 141.26, 159.57, 165.63. MS (ESI) m/z calculated for $C_{19}H_{26}ClN_4O_3$ $[M+H]^+$ 393.16, found 393.2. Retention time (method A): 0.900 min.

**Method 2: Representative Procedure: The Synthesis of Benzyl (3-((3-(aminomethyl)phenyl)carbamoyl)-5-chlorophenyl)carbamate (ASAP-0016806)**

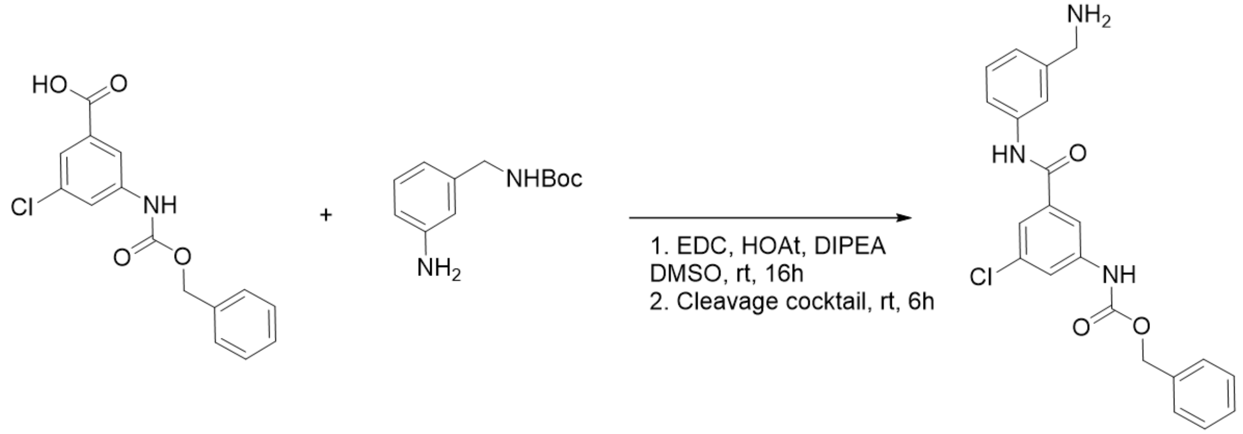

tert-Butyl N-[(3-aminophenyl)methyl]carbamate (Enamine ID: EN300-51726, 54.2 mg, 2.44 mmol), 3-{[(benzyloxy)carbonyl]amino}-5-chlorobenzoic acid (Enamine ID: EN300-4338956, 81.87 mg, 2.68 mmol), EDC (49.2 mg, 3.17 mmol.), HOAt (34.8 mg, 2.56 mmol), and diisopropylethylamine (78.77 mg, 6.10 mmol) were mixed in DMSO (appr. 0.7 ml per 100 mg of product). The mixture was sealed and stirred at ambient temperature for 16 hours. Then the cleavage cocktail (trifluoroacetic acid, triisopropylsilane, water (93:5:2; v/v) appr. 0.7 ml per 100 mg of product) was added to the mixture. The mixture was stirred at ambient temperature for 6 hours. The solvent was evaporated under reduced pressure, and the residue was dissolved in DMSO (appr. 1 ml up to 100 mg of product). The solution was filtered, analyzed by LCMS, and transferred for HPLC purification (XBridge Prep C18 5mkm OBD (Waters) using a gradient from 40% to 90% of phase B in phase A (A - 0.1% TFA in water, B - 0.1% TFA in acetonitrile)) to afford 70.9 mg (70.9% yield) of benzyl (3-((3-(aminomethyl)phenyl)carbamoyl)-5-chlorophenyl)carbamate (ASAP-0016806). LCMS(ESI): $[M+H]^+$ m/z: calcd 410.0; found 410.0; Rt = 1.010 min. $^1$H NMR (500 MHz, DMSO-d6) δ (ppm) 4.02 (m, 2H), 5.17 (s, 2H), 7.20 (d, 1H), 7.36 (m, 3H), 7.42 (m, 3H), 7.61 (d, 1H), 7.63 (s, 1H), 7.76 (s, 1H), 7.95 (s, 2H), 8.14 (br s, 3H), 10.21 (s, 1H), 10.43 (s, 1H). $^1$H NMR (125 MHz, DMSO-d6) δ (ppm): 27.38, 42.91, 66.61, 116.80, 121.04, 121.31, 121.37, 124.76, 128.63, 128.94, 129.49, 133.69, 134.90, 136.72, 137.11, 139.54, 141.28, 153.75, 164.66. $^{13}$C NMR (125 MHz, DMSO-d6) δ (ppm): 50.43, 50.73, 66.62, 115.16, 116.79, 120.97, 121.33, 123.53, 128.57, 128.93, 130.64, 133.69, 136.07, 137.68, 139.46, 141.26, 153.75, 164.36. MS (ESI) m/z calculated for $C_{22}H_{21}ClN_3O_3$ $[M+H]^+$ 410.12, found 410.0. Retention time (method A): 1.010 min.

**Method 3: The Synthesis of N-[(6-chloro-1H-indazol-4-yl)methyl]isoindolin-5-amine (ASAP-0029525)**

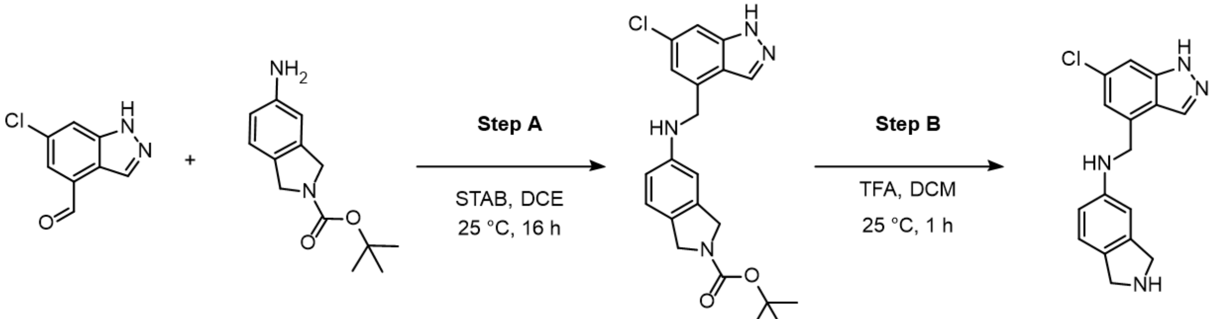

ASAP-0029525

**Step A: The synthesis of tert-butyl 5-[(6-chloro-1H-indazol-4-yl)methylamino]isoindoline-2-carboxylate**

To a stirred solution of 6-chloro-1H-indazole-4-carbaldehyde (Enamine ID: EN300-26280982, 50 mg, 276.87 µmol) and tert-butyl 5-aminoisoindoline-2-carboxylate (Enamine ID: EN300-188167, 64.87 mg, 276.87 µmol) in DCE (2 mL) were added sodium triacetoxyborohydride (117.36 mg, 553.74 µmol) at 25 °C. The resulting reaction mixture was stirred at 25 °C for 16 hours. Upon completion, the reaction mixture was quenched with water (10 mL) and DCM (20 mL). Organic phase was washed with water (10 mL), brine (10 mL), dried over $Na_2SO_4$ and concentrated under reduced pressure to obtain crude product tert-butyl 5-[(6-chloro-1H-indazol-4-yl)methylamino]isoindoline-2-carboxylate (0.12 g, 252.70 µmol, 91.27% yield, 84% purity), that was used without further purification. LCMS (ESI): m/z: calcd 398.8; found 299.0 $[M\text{-}Boc+H]^+$; Rt = 3.697 min (L764742F LCMS-31 6min_4-6x30_1-5_V.M 16:30 31.05.2024)

**Step B: The synthesis of N-[(6-chloro-1H-indazol-4-yl)methyl]isoindolin-5-amine (ASAP-0029525)**

To a stirred solution of tert-butyl 5-[(6-chloro-1H-indazol-4-yl)methylamino]isoindoline-2-carboxylate (0.12 g, 252.70 µmol) in DCM (1 mL) were added TFA (1.49 g, 13.07 mmol, 1 mL) respectively at 25 °C. The resulting reaction mixture was stirred at 25 °C for 1 hour. The solution was evaporated and obtained crude product was purified by reverse phase HPLC (SYSTEM 25-25-65% 0-1-6min $H_2O$/ACN/$NH_4OH$ flow: 60ml/min (loading pump 4ml/min ACN target mass 299 column: XBridge OBD 100x30mm 5um ) to afford product N-[(6-chloro-1H-indazol-4-yl)methyl]isoindolin-5-amine (ASAP-0029525) (30.8 mg, 103.09 µmol, 40.79% yield) as a off-white solid.
LCMS(ESI): m/z: calcd 299.0; found 299.2 $[M+H]^+$; Rt = 0.739 min
$^1$H NMR (500 MHz, DMSO-d6) δ (ppm) 3.86 (s, 4H), 4.56 (d, 2H), 6.27 (t, 1H), 6.41 (d, 1H), 6.46 (s, 1H), 6.88 (d, 1H), 7.05 (s, 1H), 7.44 (s, 1H), 8.29 (s, 1H), 13.20 (br s, 1H). $^{13}$C NMR (125 MHz, DMSO-d6) δ (ppm): 45.02, 51.96, 51.98, 51.99, 52.81, 106.27, 106.29, 108.52, 111.51, 118.79, 120.79, 122.82, 129.71, 131.36, 132.94, 136.16, 140.95, 143.19, 147.95. MS (ESI) m/z calculated for $C_{16}H_{16}ClN_4$ $[M+H]^+$ 299.12, found 299.2 Retention time (method A): 0.739 min.

**Method 4: The Synthesis of N-[(1S)-1-(6-chloro-1H-indazol-4-yl)but-3-enyl]isoindolin-5-amine (ASAP-0029049)**

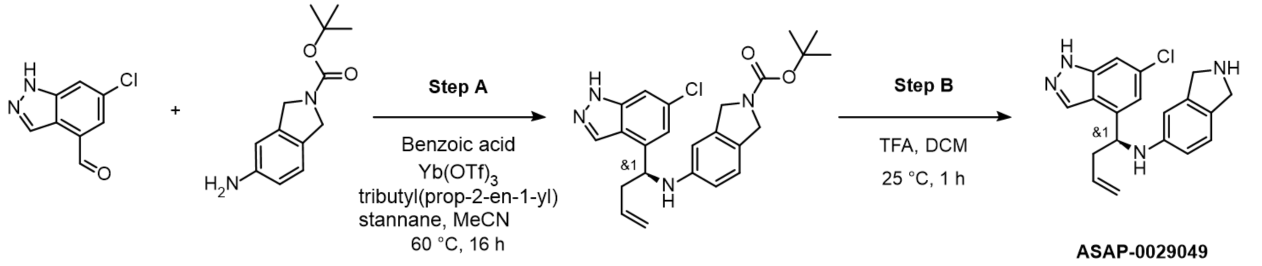

**Step A: The synthesis of tert-butyl 5-[[(1S)-1-(6-chloro-1H-indazol-4-yl)but-3-enyl]amino]isoindoline-2-carboxylate**

To a stirred solution of 6-chloro-1H-indazole-4-carbaldehyde (Enamine ID: EN300-26280982, 50 mg, 276.87 µmol), benzoic acid (33.81 mg, 276.87 µmol, 25.61 µL) and tert-butyl 5-aminoisoindoline-2-carboxylate (Enamine ID: Enamine ID: EN300-188167, 64.87 mg, 276.87 µmol) in MeCN (3 mL) were added $Yb(OTf)_3$ (17.67 mg, 27.69 µmol) and tributyl(prop-2-en-1-yl)stannane (110.01 mg, 332.24 µmol) respectively at 25°C. The resulting reaction mixture was stirred at 60 °C for 16 hours. Upon completion, SiliaMetS TAAcOH metal scavenger was added to RM, stirred for 1h, filtered, and the filtrate was concentrated under reduced pressure to obtain crude product tert-butyl 5-[[(1S)-1-(6-chloro-1H-indazol-4-yl)but-3-enyl]amino]isoindoline-2-carboxylate (276.35 mg, 195.17 µmol, 70.49% yield, 31% purity) as a yellow oil. LCMS(ESI): m/z: calcd 439.1; found 339.2 $[M\text{-}Boc+H]^+$; Rt = 3.988 min

**Step B: The synthesis of N-[(1S)-1-(6-chloro-1H-indazol-4-yl)but-3-enyl]isoindolin-5-amine (ASAP-0029049)**

To a stirred solution of tert-butyl 5-[[(1S)-1-(6-chloro-1H-indazol-4-yl)but-3-enyl]amino]isoindoline-2-carboxylate (276.35 mg, 195.17 µmol) in DCM (1 mL) was added TFA (1.49 g, 13.07 mmol, 1 mL) respectively at 25°C. The resulting reaction mixture was stirred at 25 °C for 1 hour. The obtained crude product was purified by reverse phase HPLC (Device, Mobile Phase, Column: SYSTEM

40-40-60% 0-1-6min $H_2O/ACN/NH_4OH$ flow: 60ml/min (loading pump 4ml/min ACN target mass 339 column: XBridge OBD 100x30mm 5um) to afford product N-[(1S)-1-(6-chloro-1H-indazol-4-yl)but-3-enyl]isoindolin-5-amine (**ASAP-0029049**) (38.7 mg, 114.22 µmol, 58.52% yield) as a brown gum.

LCMS(ESI): m/z: calcd 338.8; found 339.2 $[M+H]^+$; Rt = 1.037 min

$^1H$ NMR (500 MHz, DMSO-d6) δ (ppm) 2.61 (m, 2H), 4.4 - 4.27 (m, 1H), 3.9 - 3.77 (m, 2H), 4.80 (m, 1H), 4.97 (d, 1H), 5.03 (d, 1H), 5.82 (m, 1H), 6.23 (m, 1H), 6.42 (m, 2H), 6.83 (m, 1H), 7.11 (s, 1H), 7.39 (s, 1H), 8.44 (s, 1H), 13.15 (br s, 1H). $^{13}C$ NMR (125 MHz, DMSO-d6) δ (ppm): 41.52, 51.63, 52.43, 55.74, 106.71, 108.59, 111.97, 117.69, 118.66, 120.35, 122.76,

131.27, 133.02, 135.69, 141.14, 142.20. LCMS (ESI) m/z calculated for $C_{19}H_{20}ClN_4$ $[M+H]^+$ 339.17, found 339.2. Retention time (method A): 1.037 min.

**Method 5: Representative Procedure: The Synthesis of (2S)-2-(6-chloro-1H-indazol-4-yl)-2-(isoindolin-5-ylamino)-N-isopropyl-acetamide (ASAP-0029000) and single enantiomer ASAP-0029474**

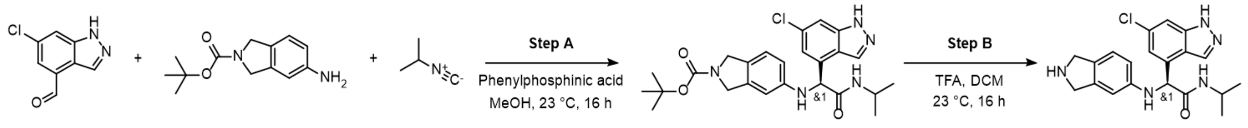

ASAP-0029000

**Step A: The synthesis of tert-butyl 5-[[(1S)-1-(6-chloro-1H-indazol-4-yl)-2-(isopropylamino)-2-oxo-ethyl]amino]isoindoline-2-carboxylate**

6-chloro-1H-indazole-4-carbaldehyde (Enamine ID: EN300-26280982, 100 mg, 553.74 µmol) and tert-butyl 5-aminoisoindoline-2-carboxylate (Enamine ID: EN300-188167, 129.74 mg, 553.74 µmol) were heated in MeOH (1 mL) at 65 °C for 1 hr. After that mixture was removed from heating bath. Phenylphosphinic acid, 98% (7.87 mg, 55.37 µmol) was added. 2-isocyanopropane (76.53 mg, 1.11 mmol) was added and the mixture was stirred at 23 °C for 16 hours. Mixture was poured into EtOAC (30 mL) and washed 3 times with $H_2O$ (20 mL). Dry organic phase concentrated under reduced pressure to provide tert-butyl 5-[[(1S)-1-(6-chloro-1H-indazol-4-yl)-2-(isopropylamino)-2-oxo-ethyl]amino]isoindoline-2-carboxylate (190 mg, 109.92 µmol, 19.85% yield, 28% purity) that was used in the next step without further purification.

LCMS(ESI): m/z: calcd 484.2; found 484.0 $[M+H]^+$; Rt = 1.342 min

**Step B: The synthesis of (2S)-2-(6-chloro-1H-indazol-4-yl)-2-(isoindolin-5-ylamino)-N-isopropyl-acetamide (ASAP-0029000)**

Tert-butyl 5-[[(1S)-1-(6-chloro-1H-indazol-4-yl)-2-(isopropylamino)-2-oxo-ethyl]amino]isoindoline-2-carboxylate (190 mg, 392.57 µmol) was dissolved in DCM (10 mL). TFA (1.49 g, 13.07 mmol, 1 mL) was added and the mixture was stirred at 23 °C for 16 hours. Mixture was concentrated under reduced pressure. Residue was purified with reverse phase HPLC (Device, Mobile Phase, Column: SYSTEM 15-15-50% 0-1-5min $H_2O$/MEOH/0.1%TFA, flow: 30ml/min (loading pump 4ml/minMEOH) target mass 384 column: Chromatorex 18 SMB100-5T 100x19mm 5) to provide (2S)-2-(6-chloro-1H-indazol-4-yl)-2-(isoindolin-5-ylamino)-N-isopropyl-acetamide (**ASAP-0029000**) (14.3 mg, 33.90 µmol, 8.64% yield, 91% purity).

LCMS(ESI): m/z: calcd 383.8; found 384.2 $[M+H]^+$; Rt = 0.730 min

1H NMR (500 MHz, DMSO-d6) δ (ppm) 0.93 (d, 3H), 1.09 (d, 3H), 3.78 (m, 1H), 4.26 (s, 2H), 4.30 (s, 2H), 5.36 (s, 1H), 6.66 (m, 3H), 7.03 (d, 1H), 7.24 (s, 1H), 7.49 (s, 1H), 8.30 (d, 1H), 8.41 (s, 1H), 9.13 (br s, 2H), 13.17 (br s, 1H). $^{13}$C NMR (125 MHz, DMSO-d6) δ (ppm): 22.51, 22.71, 41.22, 59.72, 107.01, 109.41, 113.19, 119.76, 120.60, 123.27, 131.21, 133.44, 134.81, 141.06, 147.38, 168.76. LCMS (ESI) m/z calculated for $C_{20}H_{23}ClN_5O$ $[M+H]^+$ 384.20, found 384.2. Retention time (method A): 0.730 min.

**Step C: Preparative chiral separation of ASAP-0029000 to afford enantiomer ASAP-0029974**

rac-2-(6-chloro-1H-indazol-4-yl)-2-(isoindolin-5-ylamino)-N-isopropyl-acetamide (50.0 mg, 100.63 µmol, TFA) was subjected to chiral separation employing a CHIRALPAK IC (250x20 mm, 5 mkm) column and mobile phase: Hexane(0.3% DEA):MeOH:IPA, 70:15:15 (Flow Rate: 12 mL/min, column temperature: 40 °C). ASAP-0029974 (rel-*2R*)-2-(6-chloro-1H-indazol-4-yl)-2-(isoindolin-5-ylamino)-N-isopropyl-acetamide (25.24 mg, 65.75 µmol, 65.34% yield) eluted first (RT1 = 21.09 min) and was isolated as a brown solid. Analytical RT (Chiralpak IC (250x4.6 mm, 5 mkm), Hexane(0.1% EDA):IPA(0.1% EDA):MeOH(0.1% EDA), 70:15:15; Flow Rate: 0.6 ml/min) = 15.90 min.

**Other compounds synthesized according to method 2:**

**ASAP-0020915** (benzyl (3-chloro-5-(isoindolin-5-ylcarbamoyl)phenyl)carbamate) was synthesized according to method 2, employing tert-butyl 5-amino-2,3-dihydro-1H-isoindole-2-carboxylate (Enamine ID: EN300-188167, 55.1 mg, 0.235 mmol) and 3-{[(benzyloxy)carbonyl]amino}-5-chlorobenzoic acid (Enamine ID: EN300-4338956, 79.0 mg, 0.258 mmol). Overall yield: 50.7 mg, 0.12 mmol, 51%.

$^{1}$H NMR (500 MHz, $CDCl_3$) δ (ppm) 4.47 (s, 2H), 4.51 (s, 2H), 5.17 (s, 2H), 7.35 (m, 3H), 7.41 (m, 3H), 7.62 (m, 2H), 7.76 (s, 1H), 7.88 (s, 1H), 7.93 (s, 1H), 9.38 (br s, 2H), 10.21 (s, 1H), 10.45 (s, 1H). $^{13}$C NMR (125 MHz, $CDCl_3$) δ (ppm): 50.43, 50.74, 66.62, 115.16, 116.79, 120.97, 121.33, 123.53, 128.57, 128.93, 130.64, 133.69, 136.07, 136.72, 137.68, 139.46, 141.26, 153.75, 162.33. LCMS (ESI) m/z calculated for $C_{23}H_{21}ClN_3O_3$ $[M+H]^+$ 422.21, found 422.0. Retention time (method A): 0.954 min.

**ASAP-0023261** (*N*-(3-(aminomethyl)phenyl)-6-chloro-1*H*-indazole-4-carboxamide) was synthesized according to method 2, employing 6-chloro-1H-indazole-4-carboxylic acid (EN300-6412153, 100 mg, 0.51 mmol) and tert-butyl N-[(3-aminophenyl)methyl]carbamate (Enamine ID: EN300-51726, 113 mg, 0.51 mmol). Overall yield 48 mg, 0.16 mmol, 31%.

$^{1}$H NMR (500 MHz, DMSO-d6) δ (ppm) 4.05 (m, 2H), 7.22 (d, 1H), 7.43 (t, 1H), 7.65 (d, 1H), 7.79 (s, 1H), 7.88 (s, 1H), 8.06 (s, 1H), 8.17 (br s, 3H), 8.37 (s, 1H), 10.59 (s, 1H). $^{13}$C NMR (125 MHz, DMSO-d6) δ (ppm): 42.92, 121.05, 121.16, 121.38, 124.78, 129.05, 129.50, 130.70, 134.93, 139.58, 164.53. LCMS (ESI) m/z calculated for $C_{15}H_{14}ClN_4O$ $[M+H]^+$ 300.11, found 301.0. Retention time (method A): 0.811 min.

**ASAP-0027808 (**6-chloro-N-isoindolin-5-yl-1H-indazole-4-carboxamide) was synthesized according to method 2, employing 6-chloro-1H-indazole-4-carboxylic acid (Enamine ID: EN300-6412153, 150 mg, 0.77 mmol) and tert-butyl 5-amino-2,3-dihydro-1H-isoindole-2-carboxylate (Enamine ID: EN300-188167, 180 mg, 0.77 mmol). Overall yield 44.9 mg, 0.14 mmol, 19%.

$^{1}$H NMR (500 MHz, DMSO-d6) δ (ppm) 4.03 (s, 2H), 4.07 (s, 2H), 7.22 (d, 1H), 7.54 (d, 1H), 7.75 (m, 2H), 7.84 (s, 1H), 8.37 (s, 1H), 10.41 (s, 1H). $^{13}$C NMR (125 MHz, DMSO-d6) δ (ppm): 57.19, 57.65, 118.12, 118.73, 124.16, 124.99, 125.74, 127.34, 134.23, 135.47, 142.58, 142.85, 146.00, 147.86, 169.02. LCMS (ESI) m/z calculated for $C_{16}H_{14}ClN_4O$ $[M+H]^+$ 313.18, found 313.0. Retention time (method A): 0.722 min.

*Compounds described in Figure 4 were also synthesized according to method 2.*

**Other compounds synthesized according to method 5:**

**ASAP-0031156** was synthesized according to method 5, employing 6-chloro-1H-indazole-4-carbaldehyde (Enamine ID: EN300-26280982, 100 mg, 553.74 µmol), 2-methyl-2,3-dihydro-1H-isoindol-5-amine (Enamine ID: EN300-53042, 82.07 mg, 553.74 µmol) and 1,1-difluoro-2-isocyanoethane (Enamine ID: EN300-1933591, 45.92 mg, 664.49 µmol). Overall yield: 40.4 mg, 79.07 µmol, 14% yield (TFA salt).

$^{1}H$ NMR (500 MHz, DMSO-d6) δ (ppm) 0.93 (d, 3H), 1.09 (d, 3H), 2.93 (m, 3H), 4.23 (m, 4H), 4.57 (d, 1H), 5.36 (s, 1H), 6.66 (m, 2H), 7.03 (d, 1H), 7.25 (m, 1H), 7.49 (s, 1H), 8.30 (d, 1H), 8.41 (s, 1H), 10.31 (br s, 1H). $^{13}C$ NMR (125 MHz, DMSO-d6) δ (ppm): 22.47, 22.69, 59.48, 59.65, 60.03, 109.46, 119.82, 120.54, 122.31, 123.64, 123.66, 131.21, 133.47, 134.58, 135.76, 141.07, 148.05, 168.55. LCMS (ESI) m/z calculated for $C_{16}H_{14}ClN_4O$ $[M+H]^+$ 398.2, found 398.2. Retention time (method B): 2.27 min.

**ASAP-0031651**
(N-(2,2-difluoroethyl)-2-(isoindolin-5-ylamino)-2-[3-methyl-5-(trifluoromethyl)phenyl]acetamide) was synthesized according to method 5, employing 3-methyl-5-(trifluoromethyl)benzaldehyde (Enamine ID: EN300-7361767, 70 mg, 0.37 mmol) and 1,1-difluoro-2-isocyanoethane (Enamine ID: EN300-1933591, 68 mg, 0.74 mmol). Overall yield 31.8 mg, 0.14 mmol, 47%.

*Analytical data for ASAP-0031651 are included in the next section (scale-up chemistry).*

*Compounds described in Figure 7 were also synthesized according to method 7.*

**Scale up synthesis of ASAP-0031651 and its (*S*) enantiomer ASAP-0036543**:

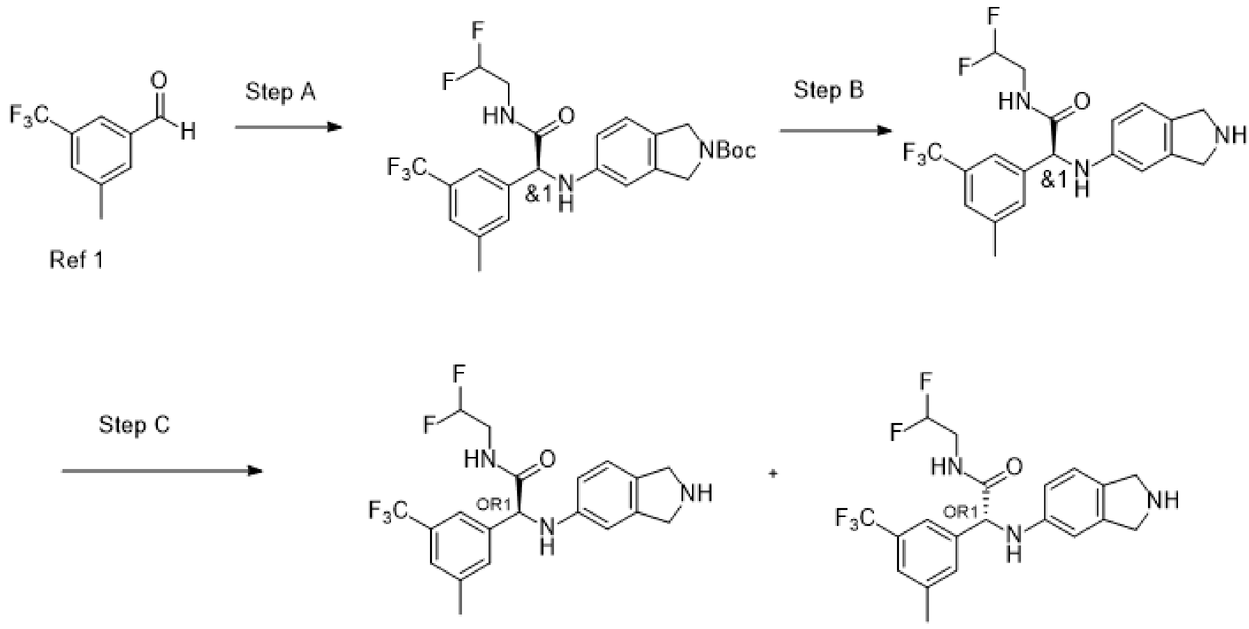

**Step A:**

3-methyl-5-(trifluoromethyl)benzaldehyde[1] (29 g, 154.14 mmol) and tert-butyl 5-aminoisoindoline-2-carboxylate (Enamine ID: EN300-188167, 36.11 g, 154.14 mmol) in MeOH (300 mL) was heated to 60 °C for 1h. After that, Phenylphosphinic acid, 98% (21.90 g, 154.14 mmol) and 1,1-difluoro-2-isocyano-ethane (14.04 g, 154.14 mmol) were added. The resulting mixture was stirred for 16h at 50 °C. The solution was concentrated, then the residue was partitioned between water (300 mL) and EtOAc (2x500mL). The organic phase was washed with brine (2x500mL), dried over Na2SO4, evaporated, and sent to prep-FCC to afford tert-butyl 5-[[2-(2,2-difluoroethylamino)-1-[3-methyl-5-(trifluoromethyl)phenyl]-2-oxo-ethyl]amino]isoindoline-2-carboxylate (34 g, 66.21 mmol, 42.96% yield).

LCMS(ESI): $[M+H]^+$ m/z: 514.2; Rt = 1.560 min.

**Step B: The synthesis of N-(2,2-difluoroethyl)-2-(isoindolin-5-ylamino)-2-[3-methyl-5-(trifluoromethyl)phenyl]acetamide (ASAP-0031651)**

To a solution of tert-butyl 5-[[2-(2,2-difluoroethylamino)-1-[3-methyl-5-(trifluoromethyl)phenyl]-2-oxo-ethyl]amino]isoindoline-2-carboxylate (15 g, 29.21 mmol) in DCM (200 mL), was added HCl 4.0 M in 1,4-dioxane, 29.21 mmol) at 0 °C. The resulting reaction was stirred for 16 h. After that period, the solution was evaporated and diluted with aq. $NaHCO_3$ (300 mL) and extracted with EtOAc (2x200mL). Organic phase was dried over Na2SO4, evaporated to afford

N-(2,2-difluoroethyl)-2-(isoindolin-5-ylamino)-2-[3-methyl-5-(trifluoromethyl)phenyl]acetamide (9.16 g, 22.16 mmol, 75.86% yield).

$^{1}$H NMR (500 MHz, DMSO-$d_6$) δ (ppm) 8.78 - 8.67 (m, 1H), 7.65 (s, 1H), 7.59 (s, 1H), 7.45 (s, 1H), 7.03 - 6.88 (m, 1H), 6.64 - 6.44 (m, 2H), 6.39 - 6.15 (m, 1H), 6.11 - 5.82 (m, 2H), 5.18 - 5.08 (m, 1H), 4.4 (d, J = 14.7 Hz, 1H), 3.89 (d, J = 9 Hz, 3H), 3.58 - 3.41 (m, 2H), 2.36 (s, 3H). $^{13}$C NMR (125 MHz, DMSO-d6) δ (ppm): 21.32, 50.57, 51.13, 60.59, 107.18, 112.89, 113.70, 114.80, 116.71, 121.55, 121.57, 123.44, 123.66, 125.30, 125.34, 125.36, 125.83, 128.00, 129.21, 129.46, 129.79, 132.46, 138.16, 139.61, 140.86, 147.55, 171.64. LCMS (ESI) m/z calculated for $C_{20}H_{21}F_5N_3O$ $[M+H]^+$ 414.16, found 414.2. Retention time (method B): 2.859 min min.

**Step C: Preparative chiral separation of ASAP-0031651 to afford (*S*) enantiomer ASAP-0036543**

Racemic N-(2,2-difluoroethyl)-2-(isoindolin-5-ylamino)-2-[3-methyl-5-(trifluoromethyl)phenyl]acetamide (9.2 g, 22.26 mmol) was subjected to chiral separation portionwise using a CHIRALCEL OD-H (250x21 mm, 5 mm) column with mobile phase: CO2/MeOH(2.0%NH3), 80:20, flow rate: 50.0 ml/min.
(2*S*)-N-(2,2-difluoroethyl)-2-(isoindolin-5-ylamino)-2-[3-methyl-5-(trifluoromethyl)phenyl]acetamide (**ASAP-0036543**) eluted first and was isolated as a brown solid (3.73 g, 8.57 mmol, 12.84% yield, retention time = 4.77 min). LCMS(ESI): $[M+H]^+$ m/z: 514.2; Rt = 1.569 min. Analytical RT (Chiralcel OD-H (250x4.6 mm, 5 mkm), 80:20 $CO_2$:MeOH(1.0%NH3); Flow Rate: 2.0 ml/min) = 4.202 min.

#### 3. NMR spectra

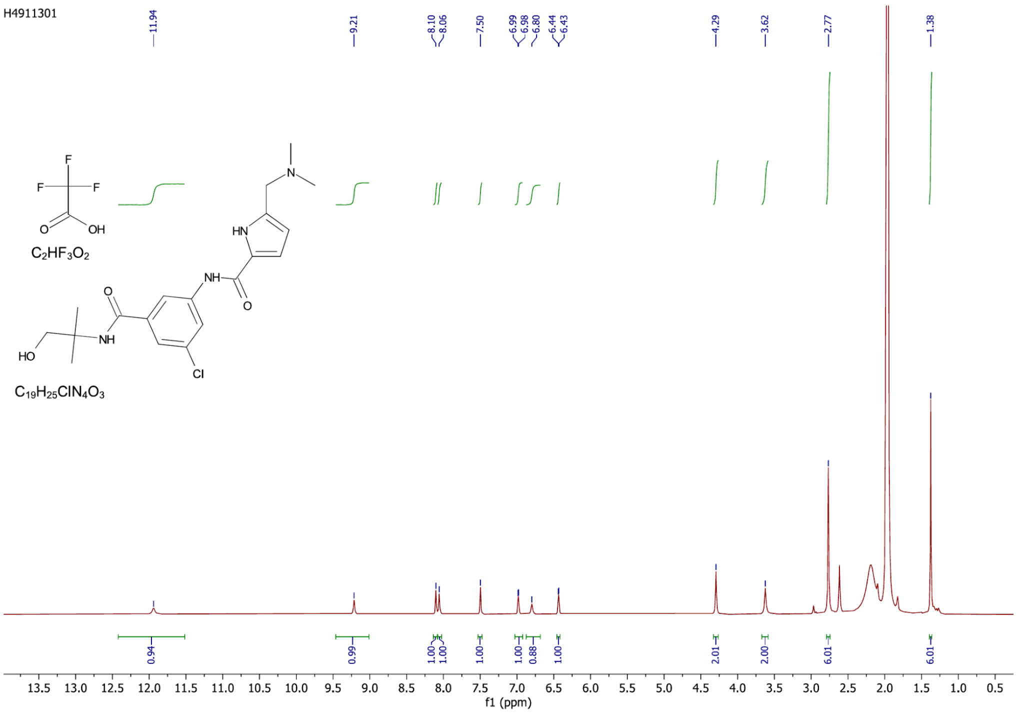

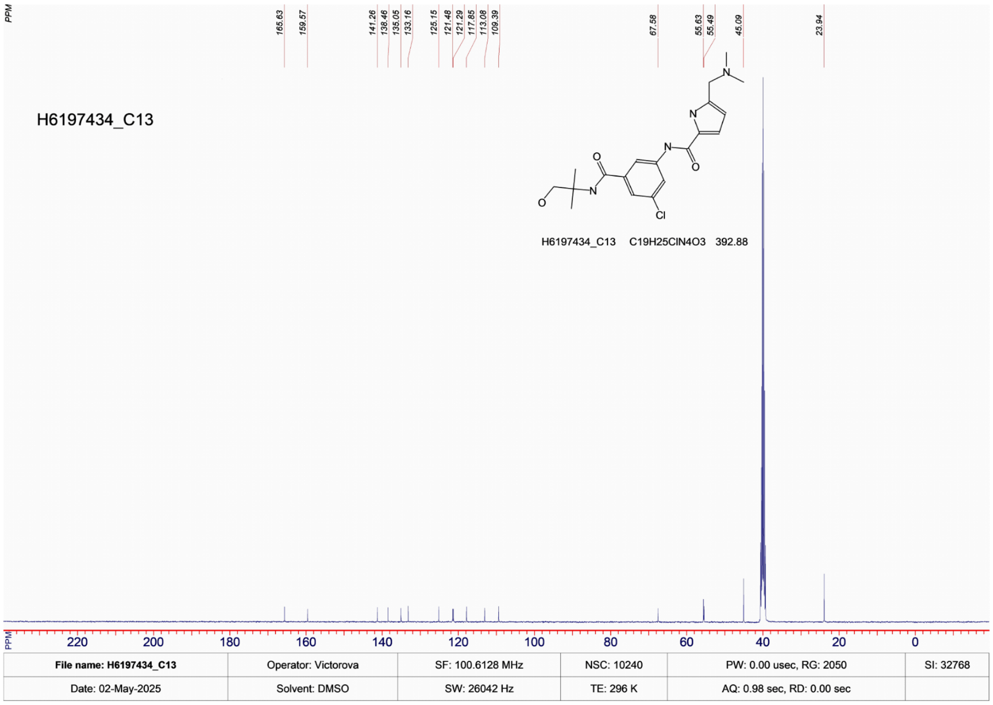

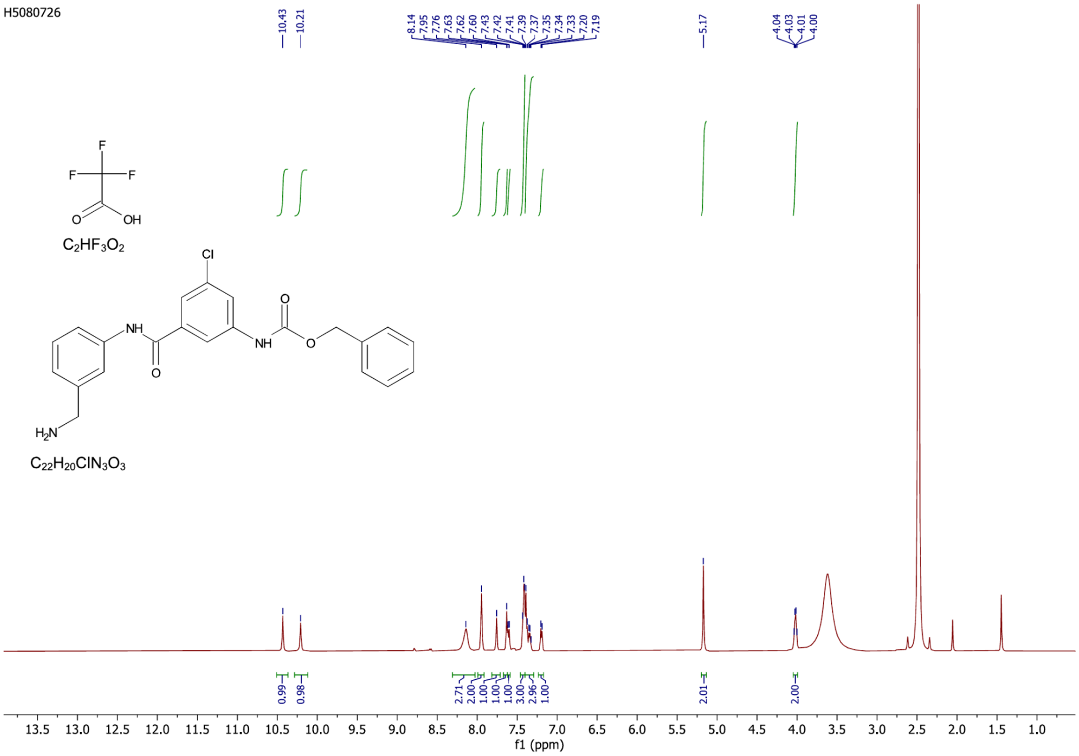

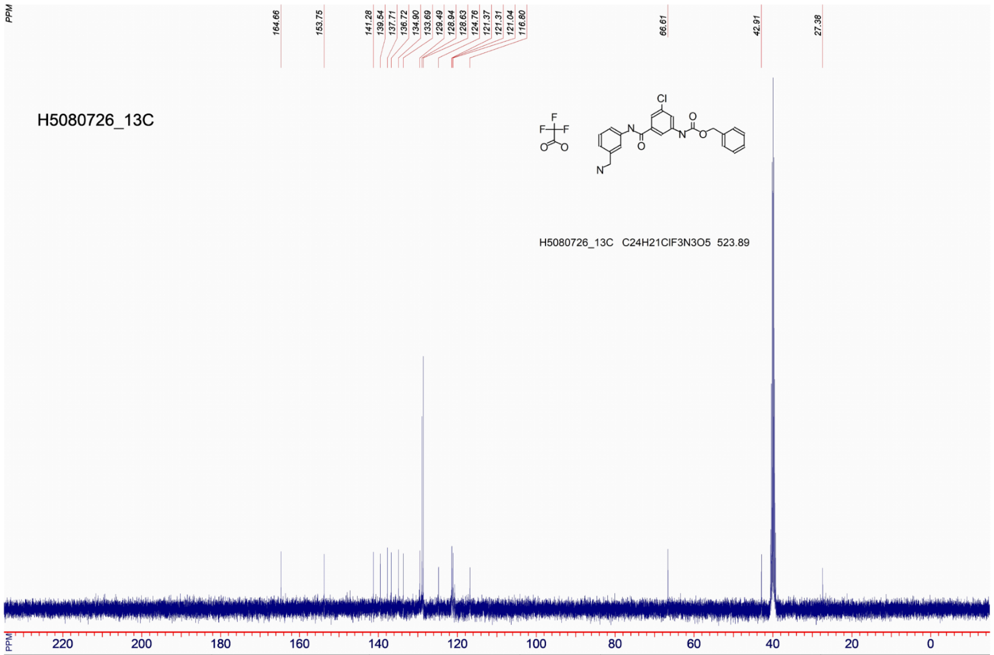

| | | | | | |
|---|---|---|---|---|---|
| File name: H5080726_13C | Operator: Victorova | SF: 100.5720 MHz | NSC: 0 | PW: 6.50 usec, RG: 30 | SI: 131072 |
| Date: 16-Apr-2025 | Solvent: dmso | SW: 25126 Hz | TE: 298 K | AQ: 1.30 sec, RD: 0.00 sec | Gradient Shimming |

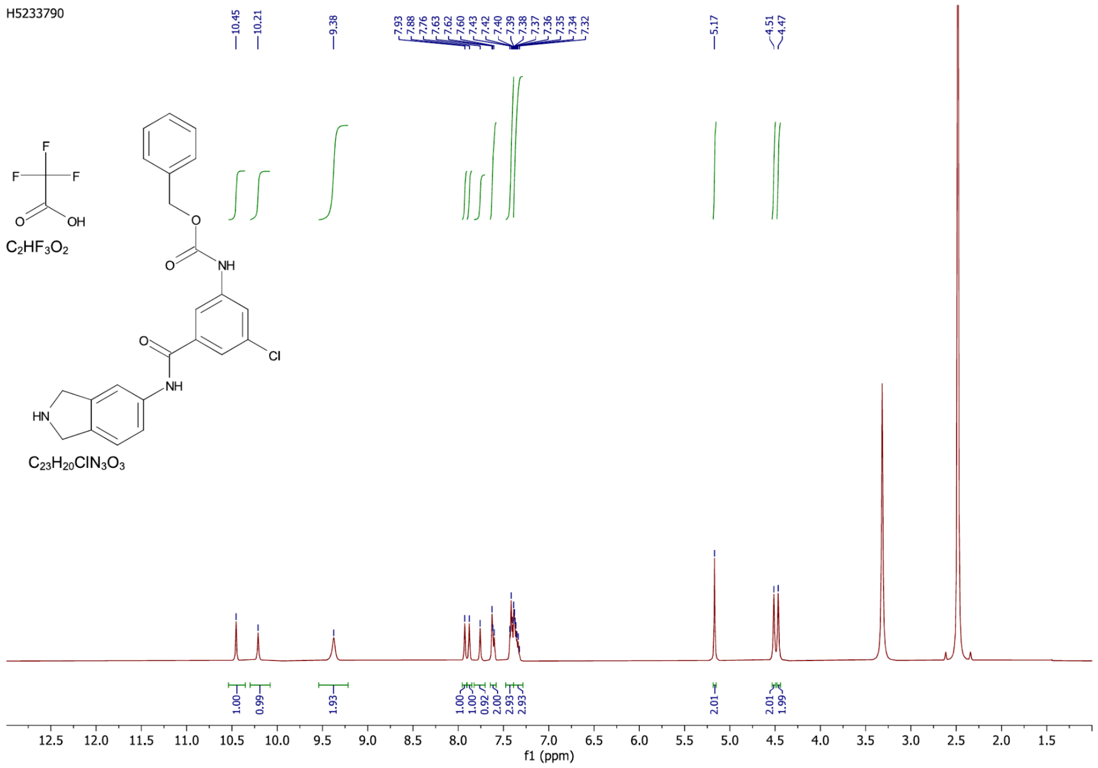

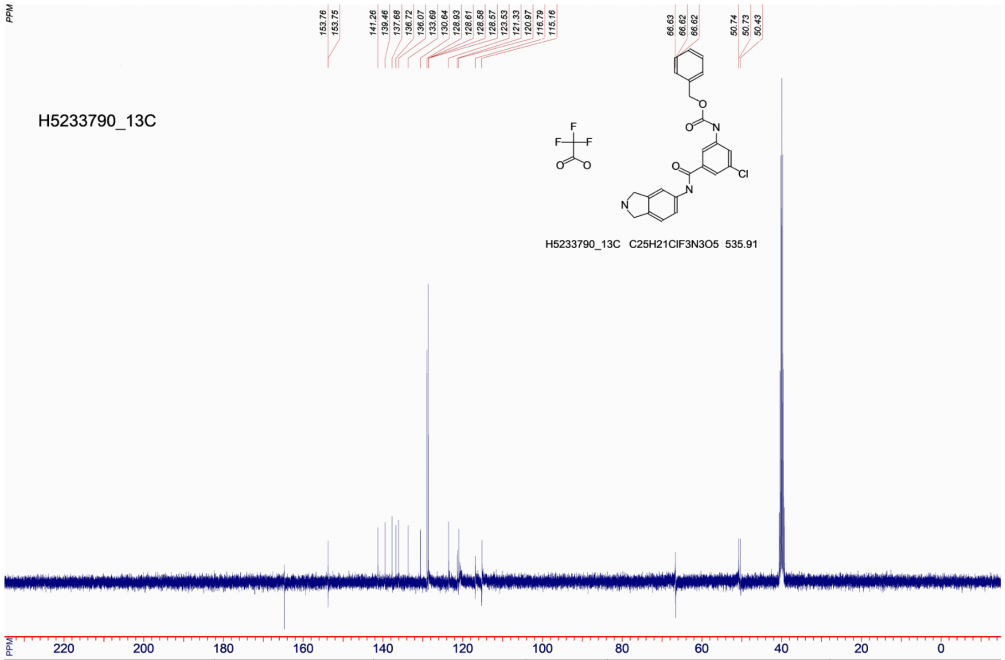

| | | | | | |
|---|---|---|---|---|---|
| **File name: H5233790_13C** | Operator: Victorova | SF: 100.5720 MHz | NSC: 0 | PW: 6.50 usec, RG: 30 | SI: 131072 |
| Date: 16-Apr-2025 | Solvent: dmso | SW: 25126 Hz | TE: 298 K | AQ: 1.30 sec, RD: 0.00 sec | Gradient Shimming |

H5283018

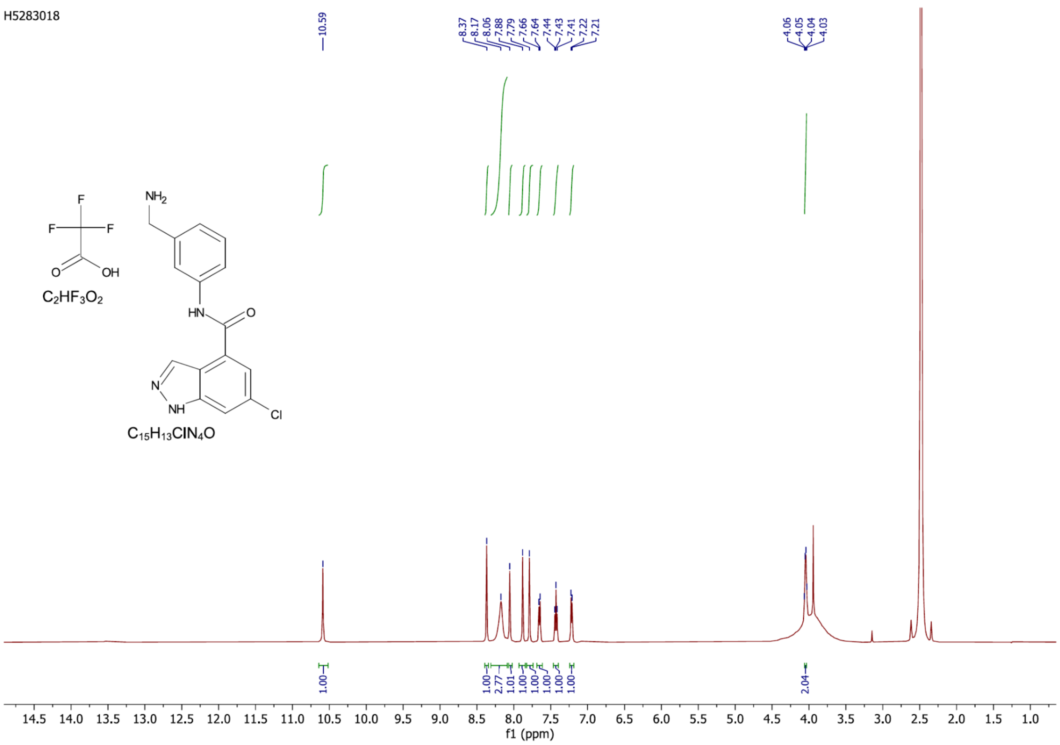

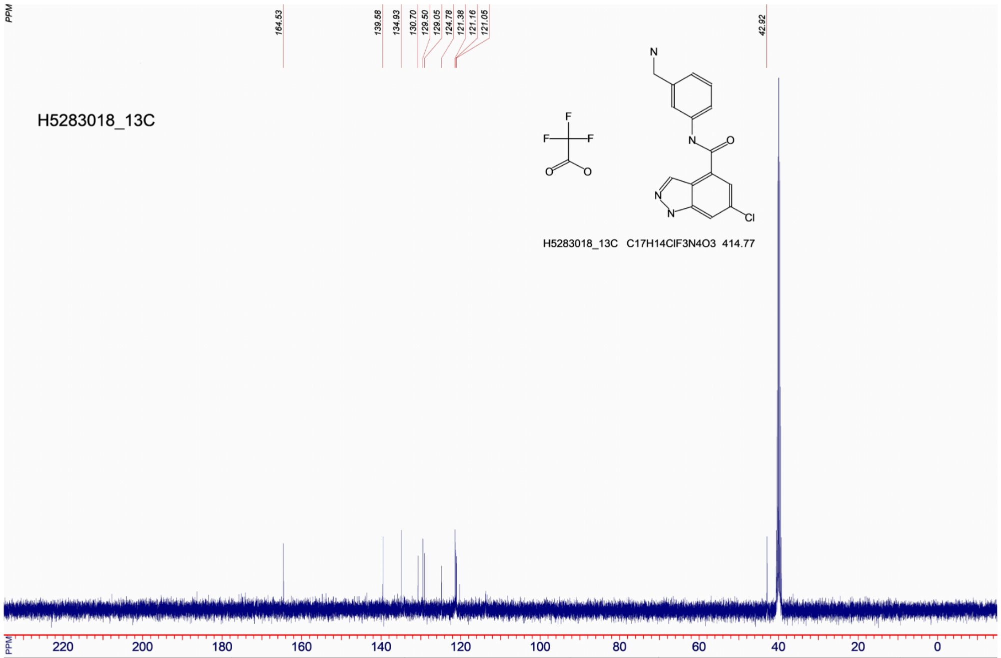

| File name: H5283018_13C | Operator: Victorova | SF: 100.5720 MHz | NSC: 0 | PW: 6.50 usec, RG: 30 | SI: 131072 |
|---|---|---|---|---|---|
| Date: 16-Apr-2025 | Solvent: dmso | SW: 25126 Hz | TE: 298 K | AQ: 1.30 sec, RD: 0.00 sec | |

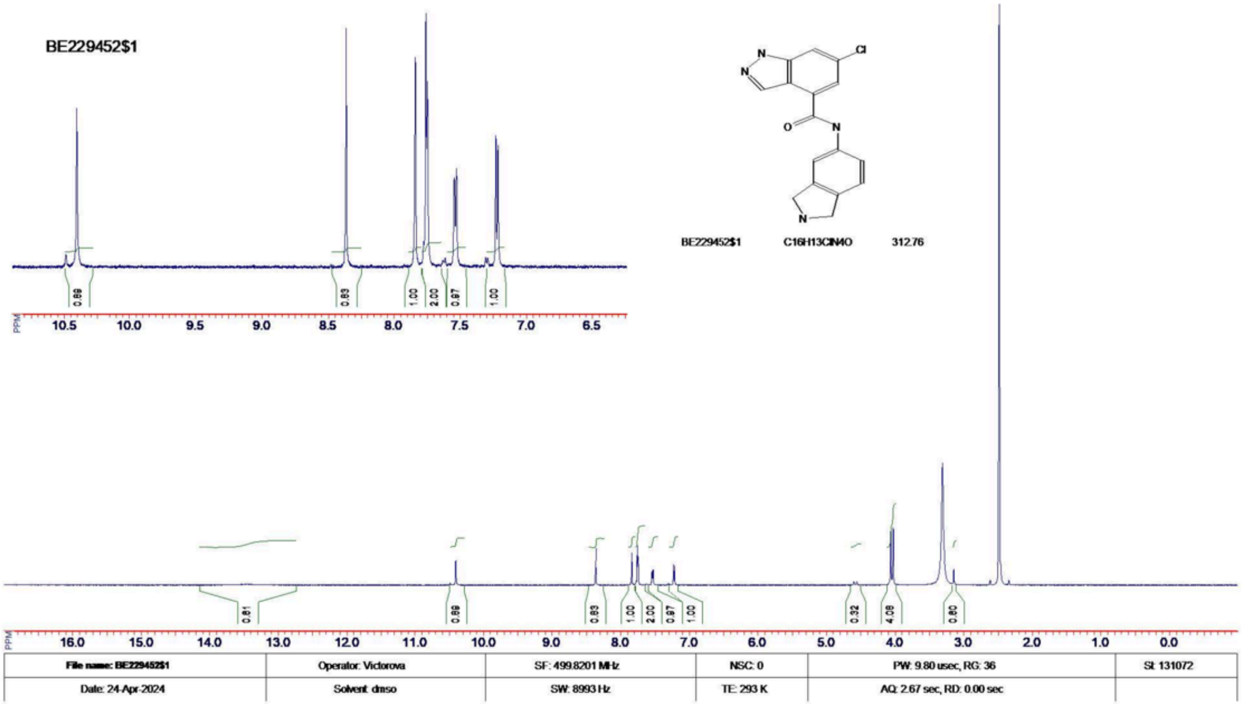
BE229452$1
Cl
N
O
BE229452$1 C16H13ClN4O 312.76
File name: BE229452$1
Operator: Victorova
SF: 499.8201 MHz
NSC: 0
PW: 9.80 usec, RG: 36
SI: 131072
Date: 24-Apr-2024
Solvent: dmso
SW: 8993 Hz
TE: 293 K
AQ: 2.67 sec, RD: 0.00 sec

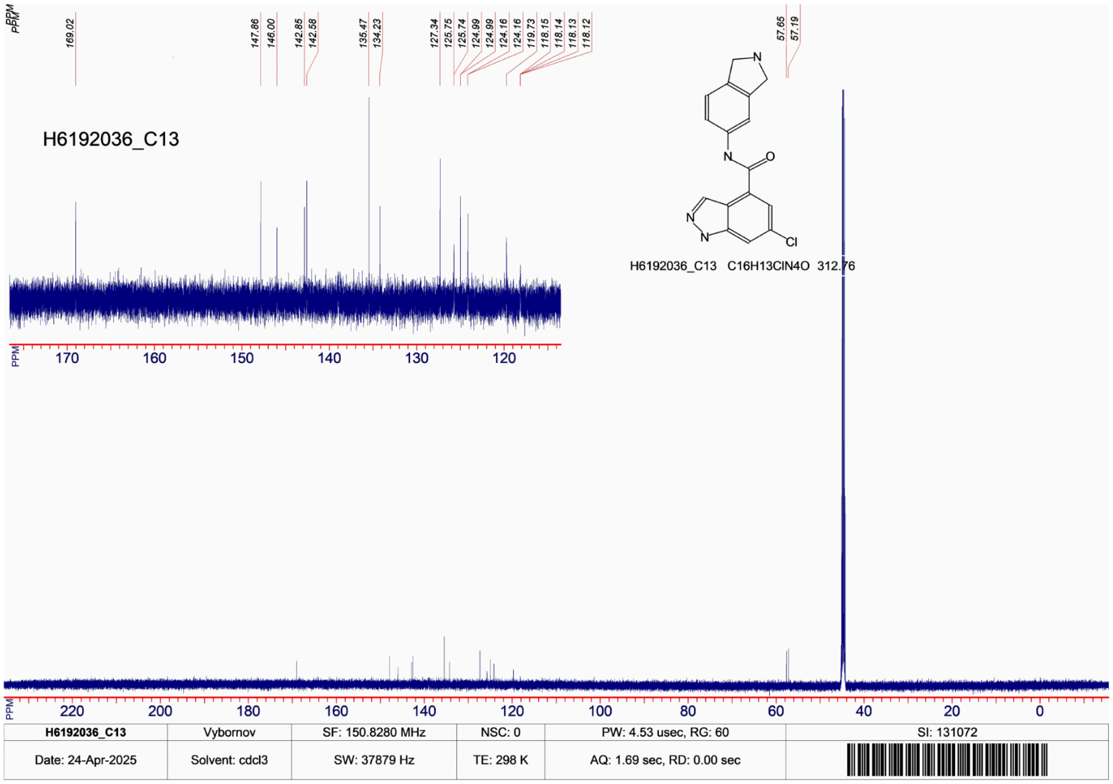
H6192036_C13
169.02
147.86
146.00
142.85
142.58
135.47
134.23
127.34
125.75
125.74
124.99
124.99
124.16
124.16
119.73
118.15
118.14
118.13
118.12
57.65
57.19
N
O
Cl
H6192036_C13 C16H13ClN4O 312.76
H6192036_C13
Vybornov
SF: 150.8280 MHz
NSC: 0
PW: 4.53 usec, RG: 60
SI: 131072
Date: 24-Apr-2025
Solvent: cdcl3
SW: 37879 Hz
TE: 298 K
AQ: 1.69 sec, RD: 0.00 sec

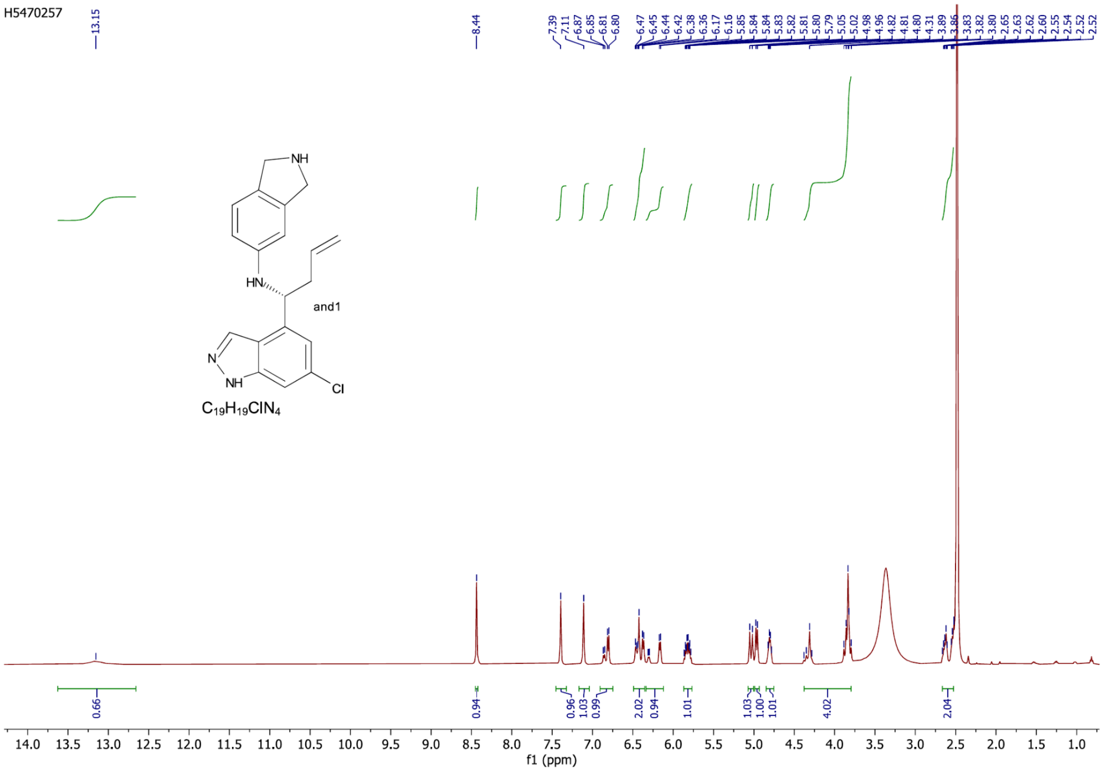

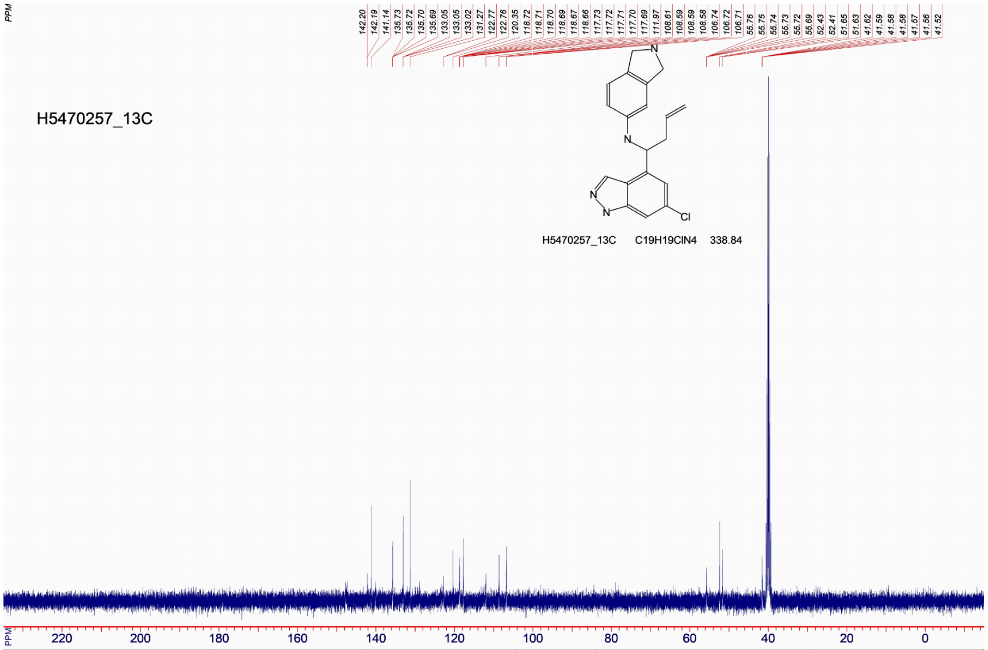

| | | | | | |
|---|---|---|---|---|---|
| **File name: H5470257_13C** | Operator: Victorova | SF: 100.5720 MHz | NSC: 0 | PW: 6.50 usec, RG: 30 | SI: 131072 |
| Date: 16-Apr-2025 | Solvent: dmso | SW: 25126 Hz | TE: 298 K | AQ: 1.30 sec, RD: 0.00 sec | Gradient Shimming |

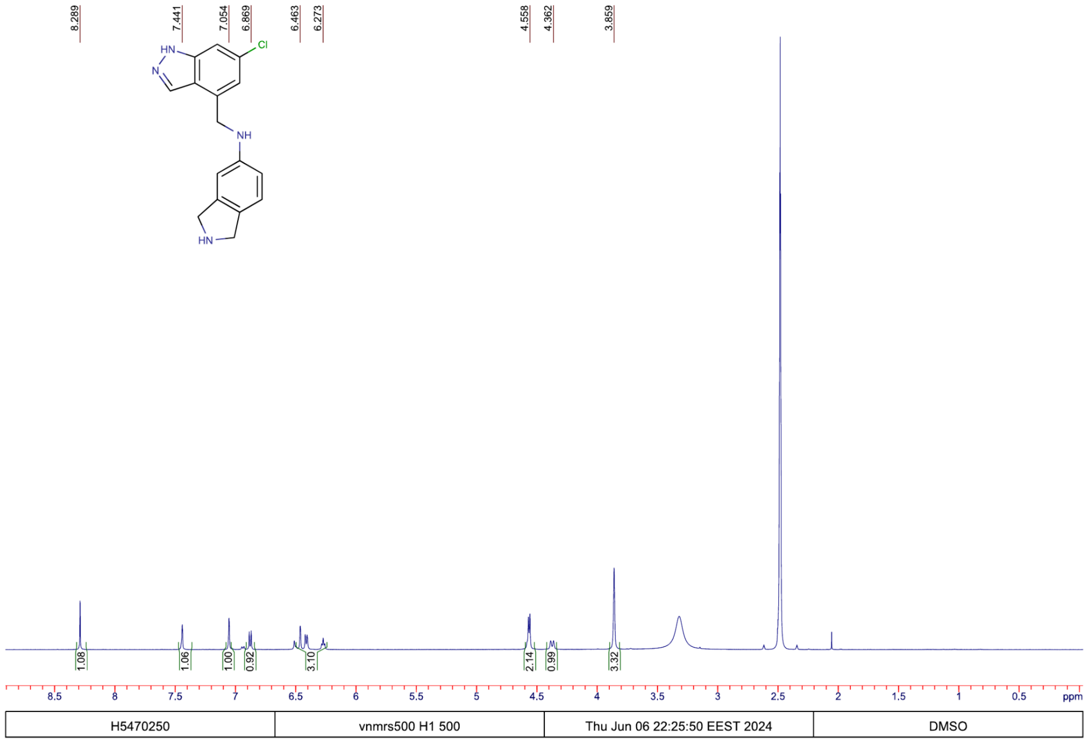
8.289
7.441
7.054
6.869
6.463
6.273
4.558
4.362
3.859
HN
N
Cl
NH
HN
1.08
1.06
1.00
0.92
3.10
2.14
0.99
3.32
8.5 8 7.5 7 6.5 6 5.5 5 4.5 4 3.5 3 2.5 2 1.5 1 0.5 ppm
H5470250
vnmrs500 H1 500
Thu Jun 06 22:25:50 EEST 2024
DMSO

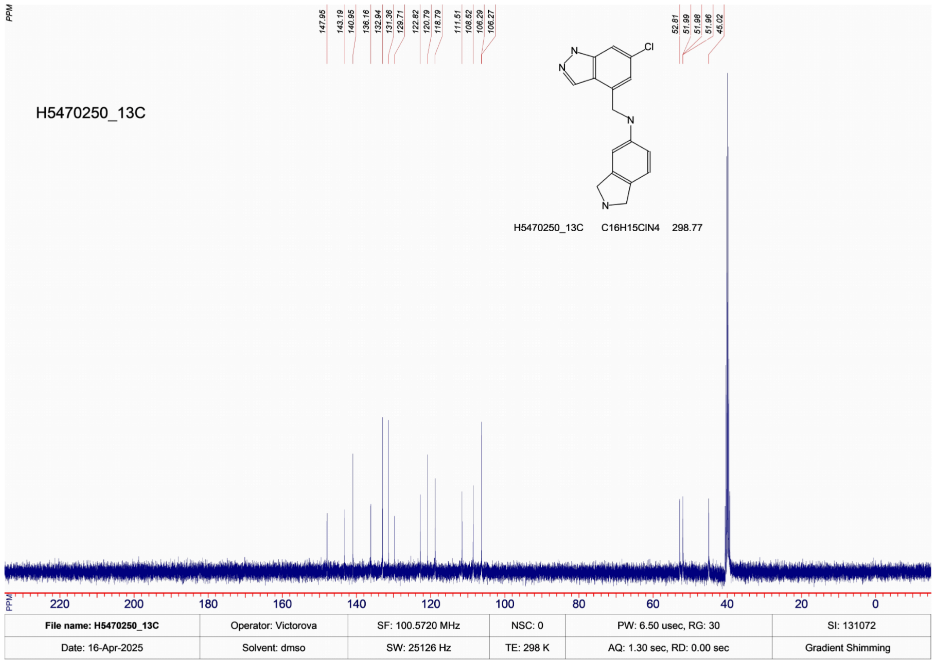
PPM
147.95
143.19
140.95
136.16
132.94
131.36
129.71
122.82
120.79
118.79
111.51
108.52
106.29
106.27
52.81
51.99
51.98
51.96
45.02
H5470250_13C
N
N
Cl
N
N
H5470250_13C C16H15ClN4 298.77
220 200 180 160 140 120 100 80 60 40 20 0
File name: H5470250_13C
Operator: Victorova
SF: 100.5720 MHz
NSC: 0
PW: 6.50 usec, RG: 30
SI: 131072
Date: 16-Apr-2025
Solvent: dmso
SW: 25126 Hz
TE: 298 K
AQ: 1.30 sec, RD: 0.00 sec
Gradient Shimming

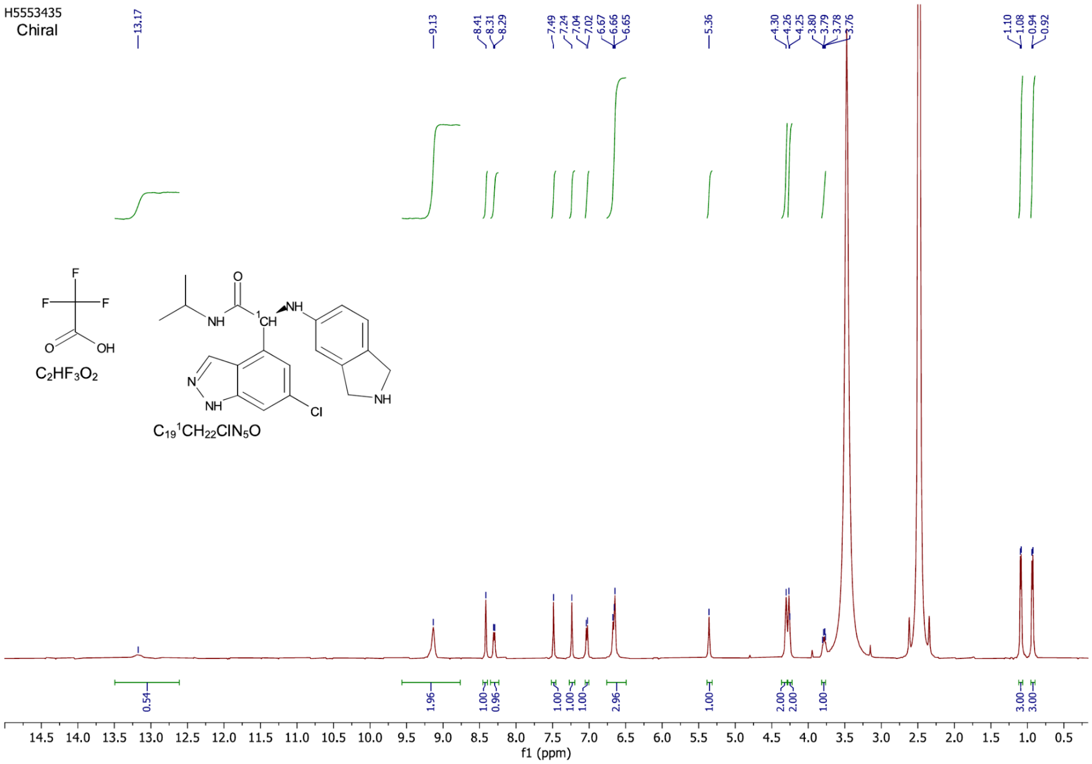

H5553435
Chiral
13.17
9.13
8.41 8.31 8.29
7.49 7.24 7.04 7.02 6.67 6.66 6.65
5.36
4.30 4.26 4.25 3.80 3.79 3.78 3.76
1.10 1.08 0.94 0.92
0.54
1.96
1.00 0.96
1.00 1.00 1.00
2.96
1.00
2.00 2.00
1.00
3.00 3.00
C2HF3O2
C19 1CH22ClN5O
f1 (ppm)

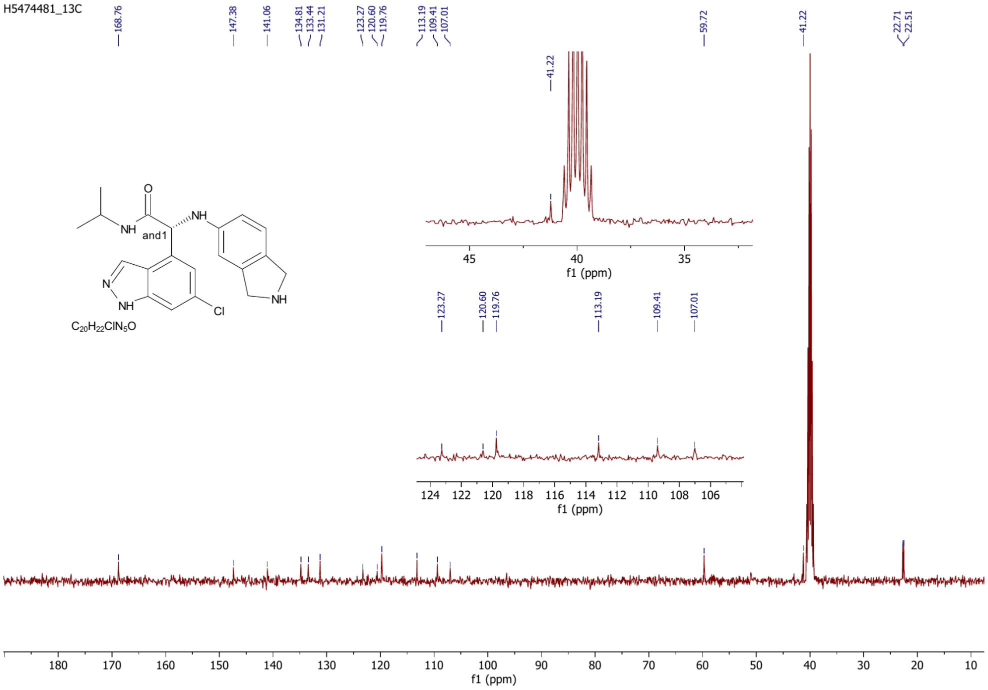

H5474481_13C
168.76
147.38
141.06
134.81 133.44 131.21
123.27 120.60 119.76
113.19 109.41 107.01
59.72
41.22
22.71 22.51
and1
C20H22ClN5O
f1 (ppm)

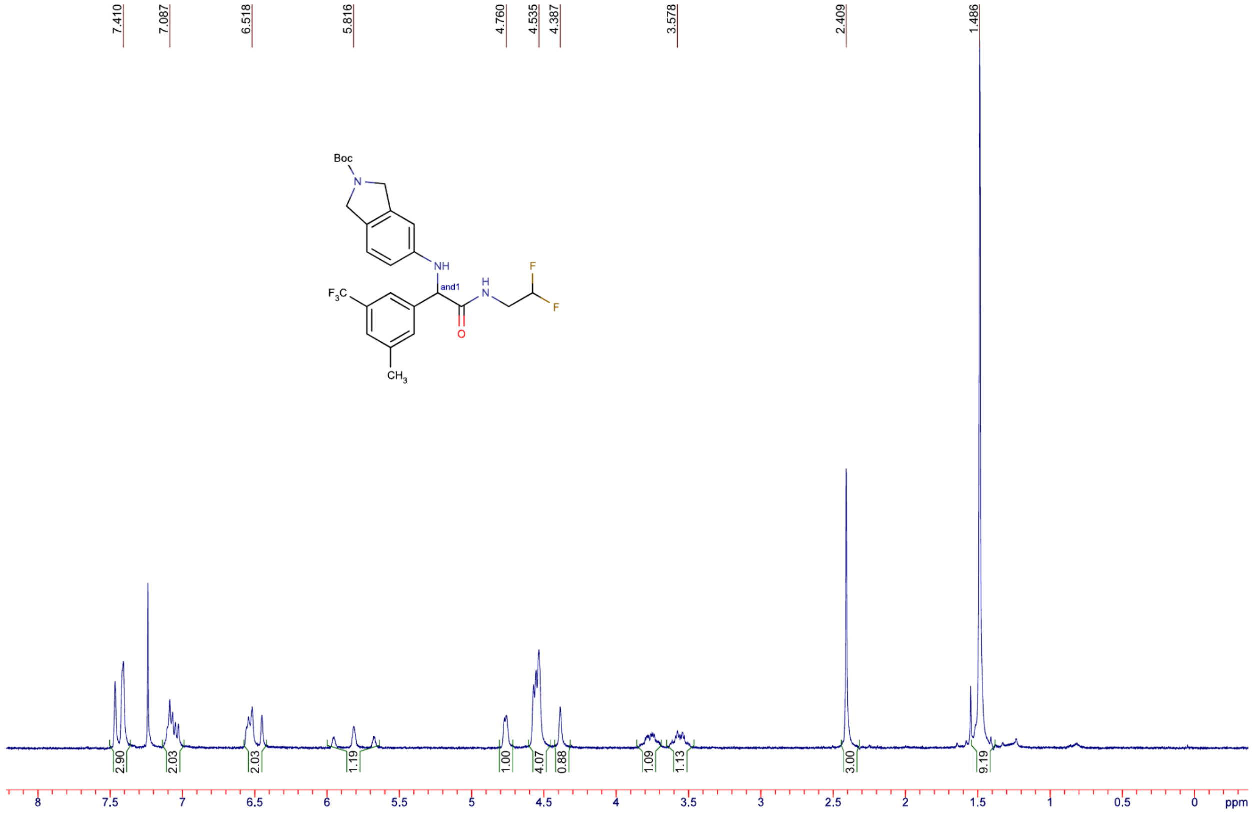

| BG563448-17 | inova400 H1 400 | Thu Feb 20 20:18:59 EET 2025 | CDCL3 |
|---|---|---|---|

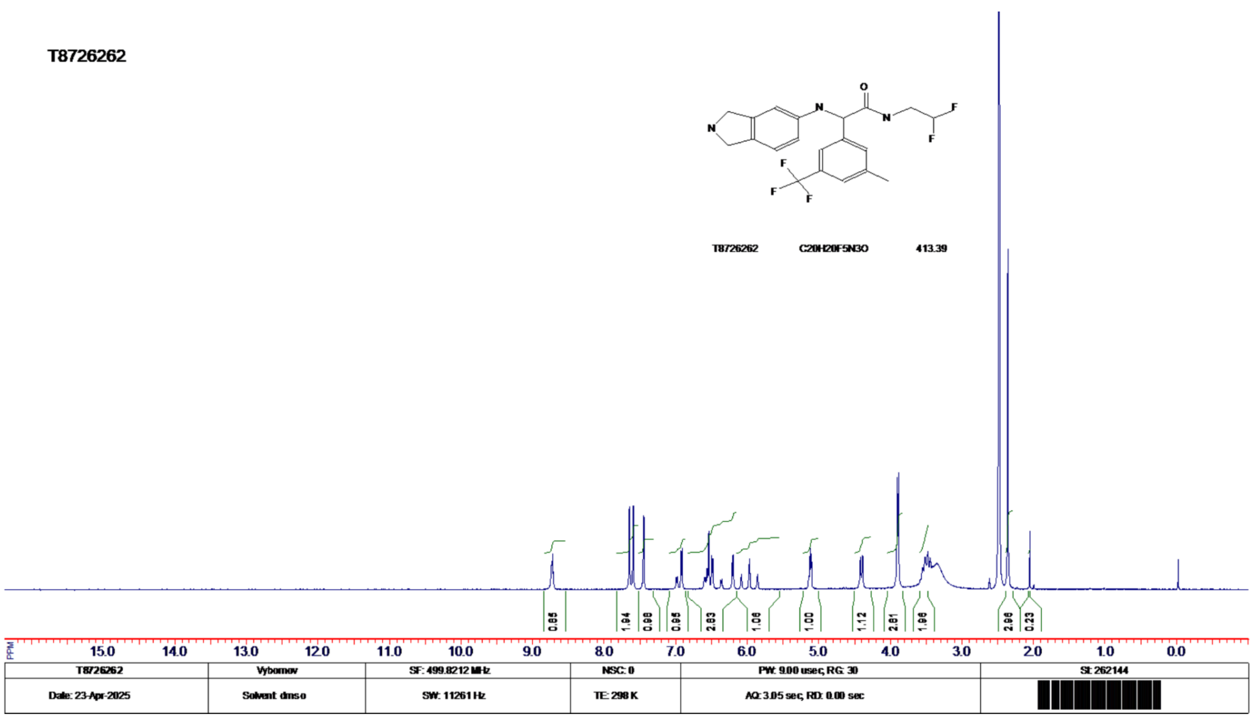
T8726262
O
N
N
N
F
F
F
F
F
T8726262 C20H20F5N3O 413.39
0.85
1.94
0.98
0.95
2.83
1.06
1.00
1.12
2.81
1.98
2.98
0.23
PPM 15.0 14.0 13.0 12.0 11.0 10.0 9.0 8.0 7.0 6.0 5.0 4.0 3.0 2.0 1.0 0.0
T8726262
Vybornov
SF: 499.8212 MHz
NSC: 0
PW: 9.00 usec, RG: 30
SI: 262144
Date: 23-Apr-2025
Solvent: dmso
SW: 11261 Hz
TE: 298 K
AQ: 3.05 sec, RD: 0.00 sec

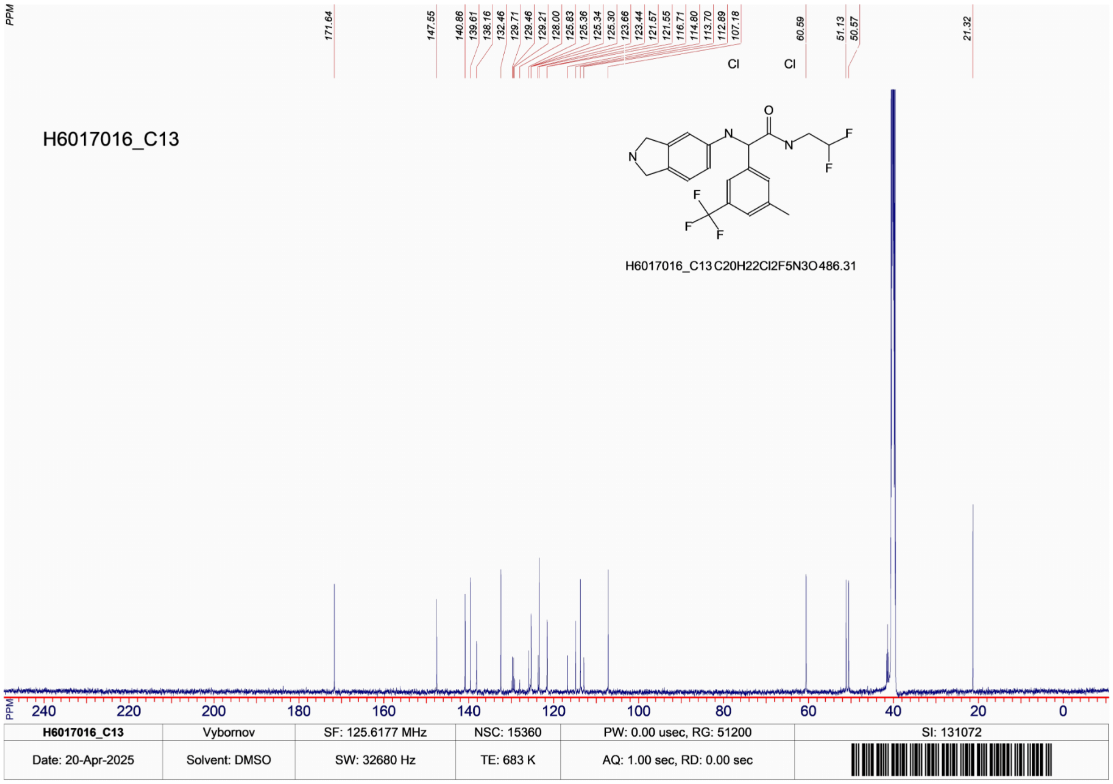
PPM
171.64
147.55
140.86
139.61
138.16
132.46
129.71
129.46
129.21
128.00
125.83
125.36
125.34
125.30
123.66
123.44
121.57
121.55
116.71
114.80
113.70
112.89
107.18
60.59
51.13
50.57
21.32
Cl
Cl
H6017016_C13
O
N
N
N
F
F
F
F
F
H6017016_C13 C20H22Cl2F5N3O 486.31
PPM 240 220 200 180 160 140 120 100 80 60 40 20 0
H6017016_C13
Vybornov
SF: 125.6177 MHz
NSC: 15360
PW: 0.00 usec, RG: 51200
SI: 131072
Date: 20-Apr-2025
Solvent: DMSO
SW: 32680 Hz
TE: 683 K
AQ: 1.00 sec, RD: 0.00 sec

#### 4. Supplementary Table 1. Data collection and refinement statistics

| Compound ID | Z68299550 | ASAP-0015373 | ASAP-0016806 |
|---|---|---|---|
| PDB accession code | 7H1J | 9RM4 | 7I9J |
| Data Collection | | | |
| Resolution[a] (Å) | 54.02 - 1.55 (1.58 -1.55) | 42.46 - 1.95 (2.00 -1.95) | 23.19 – 1.74 (1.77 -1.74) |
| Spacegroup | $P\,4_322$ | $P\,4_322$ | $P\,4_322$ |
| Cell dimensions | *a* = 42.7, b = 42.7, *c* = 216.1 Å<br>*α*= *β* = *γ* = 90.0° | *a* = 42.4, b = 42.4, *c* = 216.0 Å<br>*α*= *β* = *γ* = 90.0° | *a* = 42.7, b = 42.7, *c* = 217.5 Å<br>*α*= *β* = *γ* = 90.0° |
| No. unique reflections[a] | 30428 (1463) | 15473 (1050) | 22005 (1203) |
| Completeness[a] (%) | 100.0 (99.7) | 100.0 (100.0) | 99.8 (97.2) |
| I/σI[a] | 9.4 (0.84) | 13.8 (1.0) | 5.8 (0.2) |
| Rpim[a] | 0.026 (1.410) | 0.003 (0.746) | 0.039 (0.682) |
| CC (1/2) | 0.997 (0.414) | 0.999 (0.667) | 0.998 (0.295) |
| Redundancy[a] | 25.2 (21.5) | 16.7 (17.9) | 25.1 (25.3) |
| Refinement | | | |
| No. atoms in refinement | 1712 | 3005 | 1641 |
| Average B factor (Å$^2$) | 53.4 | 52.0 | 50.0 |
| $R_{fact}$ (%) | 23.4 | 21.5 | 24.6 |
| $R_{free}$ (%) | 27.0 | 27.8 | 28.2 |
| rms deviation bond[b] (Å) | 0.01 | 0.014 | 0.008 |
| rms deviation angle[b] (°) | 1.04 | 1.49 | 0.96 |
| Molprobity Ramachandran | | | |
| Favored (%) | 97 | 95.21 | 96.86 |
| Outlier (%) | 0 | 0 | 0 |

[a] Values in brackets show the statistics for the highest resolution shells.

[b] rms indicates root-mean-square.

**Supplementary Table 1. *(continued)* Data collection and refinement statistics**

| Compound ID | ASAP-0029000 | ASAP-0027808 | ASAP-0036543 |
|---|---|---|---|
| PDB accession code | 7I9R | 9RM6 | 9RM7 |
| Data Collection | | | |
| Resolution[a] (Å) | 54.09 – 2.60 (2.74 -2.60) | 42.25 - 1.50 (1.53 -1.50) | 54.51 – 1.45 (1.47 -1.45) |
| Spacegroup | $P\,4_322$ | $P\,4_322$ | $P\,4_322$ |
| Cell dimensions | *a* = 42.6, b = 42.6, *c* = 216.3 Å<br>*α*= *β* = *γ* = 90.0° | *a* = 42.2, b = 42.2, *c* = 218.0 Å<br>*α*= *β* = *γ* = 90.0° | *a* = 42.5, b = 42.5, *c* = 218.0 Å<br>*α*= *β* = *γ* = 90.0° |

| | | | |
|---|---|---|---|
| No. unique reflections[a] | 6844 (966) | 33083 (1576) | 36991 (1789) |
| Completeness[a] (%) | 100.0 (100.0) | 100.0 (100.0) | 100.0 (100.0) |
| I/σI[a] | 6.7 (0.9) | 24.6 (1.9) | 24.9 (1.2) |
| Rpim[a] | 0.127 (1.179) | 0.011 (0.361) | 0.012 (0.581) |
| CC (1/2) | 0.991 (0.483) | 1.0 (0.836) | 1.0 (0.714) |
| Redundancy[a] | 21.9 (23.1) | 1.8 (1.9) | 1.8 (1.9) |
| Refinement | | | |
| No. atoms in refinement | 1574 | 3014 | 3073 |
| Average B factor ($Å^2$) | 53.0 | 32.0 | 32.0 |
| $R_{fact}$ (%) | 21.7 | 19.5 | 19.5 |
| $R_{free}$ (%) | 29.1 | 22.7 | 22.2 |
| rms deviation bond[b] (Å) | 0.01 | 0.011 | 0.019 |
| rms deviation angle[b] (°) | 1.41 | 1.77 | 1.93 |
| Molprobity Ramachandran | | | |
| Favored (%) | 96.34 | 96.3 | 96.3 |
| Outlier (%) | 0 | 0 | 0 |

[a] Values in brackets show the statistics for the highest resolution shells.

[b] rms indicates root-mean-square.

5. **FRET assay for ZIKV/WNV/DENV-2 NS2B-NS3 inhibition**

This protocol is taken from https://www.science.org/doi/10.1126/science.abo7201#sec-6 with minor modifications. The following enzymes were used:

| Virus | Enzyme | MW | Extinction Coefficient | Sequence |
|---|---|---|---|---|
| ZIKV | ZV-NS2BgsgNS3 | 25556.55 | 43555 | SMSGKSVDMYIERAGDITWEKDAEVTGNSPRLDVALDESGDFSLVEDDGPPMREGGGGSGGGGGSGALWDVPAPKEVKKGETTDGVYRVMTRRLLGSTQVGVGVMQEGVFHTMWHVTKGSALRSGEGRLDPYWGDVKQDLVSYCGPWKLDAAWDGHSEVQLLAVPPGERARNIQTLPGIFKTKDGDIGAVALDYPAGTSGSPILDKCGRVIGLYGNGVVIKNGSYVSAITQGRREEETPVE |
| WNV | WNV-NS2BgsgNS3 | 26401.64 | 54430 | SMSTDMWIERTADISWESDAEITGSSERVDVRLDDDGNFQLMNDPGAPWKGGGGSGGGGGVLWDTPSPKEYKKGDTTTGVYRIMTRGLLGSYQAGAGVMVEGVFHTLWHTTKGAALMSGEGRLDPYWGSVKEDRLCYGGPWKLQHKWNGQDEVQMIVVEPGKNVKNVQTKPGVFKTPEGEIGAVTLDFPTGTSGSPIVDKNGDVIGLYGNGVIMPNGSYISAIVQGERMDEPIPAGFEPEMLRKK |
| DENV-2 | DV2-NS2BgsgNS3 | 25849.03 | 41940 | SMADLELERAADVKWEDQAEISGSSPILSITISEDGSMSIKNEEEEQTLGGGGSGGGGAGVLWDVPSPPPMGKAELEDGAYRIKQKGILGYSQIGAGVYKEGTFHTMWHVTRGAVLMHKGKRIEPSWADVKKDLISYGGGWKLEGEWKEGEEVQVLALEPGKNPRAVQTKPGLFKTNAGTIGAVSLDFSPGTSGSPIIDKKGKVVGLYGNGVVTRSGAYVSAIAQTEKSIEDNPEIEDDIFRK |

Dose response assays were performed in 12-point dilutions of twofold, starting at 25 uM for ZIKA and 100 uM for WNV and DENV-2.

Compounds were seeded into assay-ready plates (Greiner 384 low volume, cat. no. 784076) using an Echo 555 acoustic dispenser. DMSO was back-filled for a uniform concentration in assay plates (DMSO concentration 0.13% for ZIKA and 0.5% for WNV). Reagents for both assays were dispensed into the assay plate in 10 µl volumes for a final volume of 20 µl using a CERTUS FLEX dispenser.

Final enzyme concentrations were 15 nM for ZIKV NS2B-NS3 and 100 nM for WNV in buffer of 20 mM Tris pH 8.5, 0.01% Triton and 10% glycerol. Concentration of the fluorogenic peptide substrate (Boc-Gly-Arg-Arg-AMC, CAS # [113866-14-1, Biosynth FB110553) was used for ZIKV + WNV enzymes was 5 µM. For DENV-2, the same procedure was used, except with the substrate Bz-Nle-Lys-Arg-Arg-AMC (CAS # 863975-32-0; Cayman #27710).

NS2B-NS3 proteases were pre-incubated for 2 hr followed by the addition of substrate and a further 30-min incubation with the substrate (all incubations performed at room temperature).

Protease reaction was measured in a BMG Pherastar FS with a 360/470 excitation/emission filter set. Raw data were mapped and normalized to high (Protease with DMSO, no compounds) and low (No Protease, no compounds) controls using Genedata Screener software. Normalized data were then uploaded to CDD Vault (Collaborative Drug Discovery). Dose response curves were generated for $IC_{50}$ using nonlinear regression with the Levenberg–Marquardt algorithm with minimum inhibition = 0% and maximum inhibition = 100%.

To each run known inhibitors were added ASAP-0015081 (CN-716, see https://www.science.org/doi/10.1126/science.aag2419) and ASAP-0000570 (see https://pubs.acs.org/doi/10.1021/acs.jmedchem.5b01441, compound **83**), to serve as positive controls.

Km values for the substrates are provided below; these values were used to calculate Ki values from IC50 values using the Cheng-Prusoff equation from 5 uM substrate.

| | Km value (uM) | 95% CI (profile likelihood) |
|---|---|---|
| ZIKA | 43.81 | 35.03 - 55.44 |
| DENV-2 | 35.15 | 27.08 - 46.19 |
| WNV | 81.82 | 73.92 - 90.82 |

#### 6. Antiviral assays

**Zika virus antiviral assay in SH-SY5Y cells**

Cell Culture

SH-SY5Y cells (CRL-2266, ATCC) are propagated in GROWTH MEDIUM which is prepared by supplementing DMEM (Gibco cat no 41965-039) with heat-inactivated 10% v/v FCS.

Cells are cultured in T150 flasks and split 1:10 once a week.

Splitting cells:

1. Prepare cell suspension:
   a. Start with confluent T150 flasks
   b. Wash monolayer with DPBS, and aspirate
   c. Add 2 mL trypsin 0.25%trypsine/EDTA, and aspirate
   d. Incubate 5 minutes at 37°C
   e. Resuspend thoroughly to separate the cells in 10 mL GROWTH MEDIUM

2. Transfer 1 mL to x new T150 flasks prefilled with 25 mL growth medium

   3. This is your mother culture, which you will use one flask to split the cells in the following weeks. You can use the other flasks to seed your assay plates.

4. Incubate at 37°C and 5% $CO_2$

Antiviral Assay

ASSAY MEDIUM is prepared by supplementing MEM (Gibco cat no 21090-022) with heat-inactivated 2% v/v FCS, 1% sodium bicarbonate (Gibco cat no 25080-060), 1% glutamine (Gibco cat no 25030-024) and 1% NEAA (Gibco cat no 11140-035) .

1. Prepare 96w plates with cells:
   a. Start with confluent T150 flasks
   b. Wash monolayer with DPBS
   c. Add 2 mL trypsin 0.25%trypsine/EDTA, and aspirate
   d. Incubate 5 minutes at 37°C
   e. Resuspend in 10 mL ASSAY MEDIUM
   f. Count cells using coulter (2 samples are counted)
   g. Resuspend at 10000 cells/100 µL in ASSAY MEDIUM
      Add CP100356 at a concentration of 1 µM (stock = 2 mM, dilute 1/2000 in cell suspension)
   h. Add 100 µL cell suspension to each well of a 96w plate (Falcon)
   i. Incubate overnight at 37°C and 5% $CO_2$

2. Adding Compound:

a. Add 50 µL medium to column 2

b. Add compound to column 2, rows B-G:

i. In case of a 100 µM final start concentration, add 3 µL of a 10 mM stock to column 2

ii. In case of a 10 µM final start concentration, first prepare a 1:10 compound dilution in medium (for example mix 10 µL of a 10 mM compound stock with 90 µL medium in column 1) and subsequently add 3 µL of the 1:10 compound dilution to column 2

c. Make a 1:3 serial dilution of the compounds by transferring 50 µL from one column to the next column (columns 2-9) with tip change in columns 4 and 7.

3. Adding Virus:

a. Prepare virus (ZIKV MR766) to the appropriate dilution in ASSAY MEDIUM

i. Titer of the ZIKV MR766 stock is 2.6E+07 PFU/ml: prepare 1:400 (final dilution 1:1600)

b. Add 50 µL of the virus solution to columns 2-10

c. Add 50 µL of assay medium to columns 2-11.

(thus, 100 µL of cells containing 1 µM CP100356 + 50 µL cpd + 50 µL virus @ final concentration CP100356 = 0.5 µM)

i. In case of insufficient cells: medium instead of cells can be added to the border wells

4. Incubate plates (37°C / 5% $CO_2$)

5. On day 7 post-infection, determine cell viability using MTS (see Jochmans D et al. J Virol Methods. 2012 Aug;183(2):176-9. doi: 10.1016/j.jviromet.2012.04.011)

**Zika virus antiviral assay in Vero cells**

Vero E6 cells (Vero C1008;ATCC CRL-1586) are propagated in GROWTH MEDIUM which is prepared by supplementing DMEM (Gibco cat no 41965-039) with 10% v/v heat-inactivated FCS and 1% Sodium Bicarbonate (Gibco cat no 25080-060).

Cells are cultured in T150 bottle and split 1/10 once a week.

Splitting cells:

1. Prepare cell suspension:
   a. Start with confluent T150 bottle
   b. Wash monolayer with DPBS, and aspirate
   c. Add 3 mL trypsin 0.25%trypsine/EDTA, and aspirate
   d. Incubate 15 minutes at 37°C
   e. Resuspend thoroughly to separate the cells in 10 mL GROWTH MEDIUM

2. Transfer 1 mL to x new T150 bottles prefilled with 35 ml growth medium

3. For one of the bottles you add the antibiotics. This is your mother culture, and this one you will use to split the cells for the week after. The other 3 bottles you can use to seed your plates.

4. Incubate at 37°C, 5%$CO_2$

Antiviral Assay

ASSAY MEDIUM is prepared by supplementing MEM (Gibco cat no) with 2% v/v heat-inactivated FCS, 1% Sodium Bicarbonate (Gibco cat no 25080-060) and 1% Glutamine (Gibco cat no 25030-024).

1. Prepare cell suspension:
   a. Start with confluent T150 bottle
   b. Wash monolayer with DPBS
   c. Add 3 mL trypsin 0.25%trypsine/EDTA, and aspirate
   d. Incubate 15 minutes at 37°C
   e. Resuspend in 10 mL ASSAY MEDIUM
   f. Count cells using coulter (3 samples are counted)
   g. Resuspend at 10000 cells/50 µL in ASSAY MEDIUM
   Add CP100356 at a concentration of 2 µM (stock = 2 mM, dilute 1/1000 in cell suspension)

2. Adding Compound:
   a. Add 100 µL ASSAY medium to each well of a 96w plate (Falcon)
   b. Add 50 µL medium to C2
   c. Add compound to C2 row B-G:
      i. For stock 10 mM to 100 µM final start conc. you need to add 3 µL

ii. For stock 10 mM and 10 µM final start conc. you prepare a 1/10 predilution in medium (for example mix 10 µL 10 mM with 90 µL in C1) and add 3 µL

d. Dilute 1/3 over the plate (transfer 50 µL) from C2-9 – change tips in C4 and C7

3. Adding Virus:
   a. Prepare virus (ZIKV MR766) to appropriate dilution in ASSAY MEDIUM
      i. Titer of the ZIKV MR766 stock is 1.98E+05 PFU/ml: prepare 1/100 (final dilution 1/400; MOI 0.01)
   b. Add 50 µL of virus preparation to C1-10
   c. Add 50 µl of cell suspension to C1-12
   (thus, add 50 µl of cells containing 2 µM CP100356 to 150 µl cmpd + virus already in plate ® final concentration CP100356 = 0.5 µM)
      i. in case of insufficient cells: medium instead of cells can be added to border wells

4. Incubate plates (37°C / 5% $CO_2$)

5. On day 7 post-infection, determine cell viability using MTS (see Jochmans D et al. J Virol Methods. 2012 Aug;183(2):176-9. doi: 10.1016/j.jviromet.2012.04.011)

**Zika virus antiviral assay in JEG-3 cells**

Cell Culture

JEG-3 cells (HTB-36, ATCC) are propagated in GROWTH MEDIUM which is prepared by supplementing MEM (Gibco cat no 25080-060) with heat-inactivated 10% v/v FCS, 1% glutamine (Gibco cat no 25030-024), 1% NEAA (Gibco cat no 11140-035), 1% HEPES (Gibco cat no 15630-056) and 1% sodium pyruvate (Gibco cat no 11360-070) .

Cells are cultured in T150 flasks and split 1:10 twice a week.

Splitting cells:

1. Prepare cell suspension:
   a. Start with confluent T150 flasks
   b. Wash monolayer with DPBS, and aspirate
   c. Add 2 mL trypsin 0.25%trypsine/EDTA, and aspirate
   d. Incubate 5 minutes at RT
   e. Resuspend thoroughly to separate the cells in 10 mL GROWTH MEDIUM

2. Transfer 1 mL to x new T150 flasks prefilled with 25 mL growth medium

3. This is your mother culture, which you will use one flask to split the cells in the following weeks. You can use the other flasks to seed your assay plates.

4. Incubate at 37°C and 5% $CO_2$

Antiviral Assay

GROWTH Medium is also used as ASSAY MEDIUM.

1. Prepare 96w plates with cells:
   a. Start with confluent T150 flasks
   b. Wash monolayer with DPBS
   c. Add 2 mL trypsin 0.25%trypsine/EDTA, and aspirate
   d. Incubate 5 minutes at RT
   e. Resuspend in 10 mL ASSAY MEDIUM
   f. Count cells using coulter (2 samples are counted)
   g. Resuspend at 3000 cells/100 µL in ASSAY MEDIUM
   Add CP100356 at a concentration of 1 µM (stock = 2 mM, dilute 1/2000 in cell suspension)
   h. Add 100 µL cell suspension to each well of a 96w plate (Falcon)
   i. Incubate overnight at 37°C and 5% $CO_2$

2. Adding Compound:
   a. Add 50 µL medium to column 2

b. Add compound to column 2, rows B-G:
   i. In case of a 100 µM final start concentration, add 3 µL of a 10 mM stock to column 2
   ii. In case of a 10 µM final start concentration, first prepare a 1:10 compound dilution in medium (for example mix 10 µL of a 10 mM compound stock with 90 µL medium in column 1) and subsequently add 3 µL of the 1:10 compound dilution to column 2
   c. Make a 1:3 serial dilution of the compounds by transferring 50 µL from one column to the next column (columns 2-9) with tip change in columns 4 and 7.

3. Adding Virus:
   a. Prepare virus (ZIKV MR766) to the appropriate dilution in ASSAY MEDIUM
      i. Titer of the ZIKV MR766 stock is 2.6E+07 PFU/ml: prepare 1:400 (final dilution 1:1600)
   b. Add 50 µL of the virus solution to columns 2-10
   c. Add 50 µL of assay medium to columns 2-11.
   (thus, 100 µL of cells containing 1 µM CP100356 + 50 µL cpd + 50 µL virus ® final concentration CP100356 = 0.5 µM)
      i. In case of insufficient cells: medium instead of cells can be added to the border wells

4. Incubate plates (37°C / 5% $CO_2$)

5. On day 4 post-infection, determine cell viability using MTS (see Jochmans D et al. J Virol Methods. 2012 Aug;183(2):176-9. doi: 10.1016/j.jviromet.2012.04.011)

**West Nile virus antiviral assay in Vero cells**

Cell Culture

Vero E6 cells (Vero C1008;ATCC CRL-1586) are propagated in GROWTH MEDIUM which is prepared by supplementing DMEM (Gibco cat no 41965-039) with heat-inactivated 10% v/v FCS and 1% sodium bicarbonate (Gibco cat no 25080-060).

Cells are cultured in T150 flasks and split 1:10 once a week.

Splitting cells:

1. Prepare cell suspension:
   a. Start with confluent T150 flasks
   b. Wash monolayer with DPBS, and aspirate
   c. Add 2 mL trypsin 0.25%trypsine/EDTA, and aspirate
   d. Incubate 5 minutes at 37°C
   e. Resuspend thoroughly to separate the cells in 10 mL GROWTH MEDIUM

2. Transfer 1 mL to x new T150 flasks prefilled with 25 mL growth medium

3. This is your mother culture, which you will use one flask to split the cells in the following weeks. You can use the other flasks to seed your assay plates.

4. Incubate at 37°C and 5% $CO_2$

Antiviral Assay

ASSAY MEDIUM is prepared by supplementing MEM (Gibco cat no 21090-022) with heat-inactivated 2% v/v FCS, 1% sodium bicarbonate (Gibco cat no 25080-060), 1% glutamine (Gibco cat no 25030-024) and 1% NEAA (Gibco cat no 11140-035) .

1. Prepare 96w plates with cells:
   a. Start with confluent T150 flasks
   b. Wash monolayer with DPBS
   c. Add 2 mL trypsin 0.25%trypsine/EDTA, and aspirate
   d. Incubate 5 minutes at 37°C
   e. Resuspend in 10 mL ASSAY MEDIUM
   f. Count cells using coulter (2 samples are counted)
   g. Resuspend at 10000 cells/100 µL in ASSAY MEDIUM
   Add CP100356 at a concentration of 1 µM (stock = 2 mM, dilute 1/2000 in cell suspension)
   h. Add 100 µL cell suspension to each well of a 96w plate (Falcon)
   i. Incubate overnight at 37°C and 5% $CO_2$

2. Adding Compound:
   a. Add 50 µL medium to column 2
   b. Add compound to column 2, rows B-G:
      i. In case of a 100 µM final start concentration, add 3 µL of a 10 mM stock to column 2
      ii. In case of a 10 µM final start concentration, first prepare a 1:10 compound dilution in medium (for example mix 10 µL of a 10 mM compound stock with 90 µL medium in column 1) and subsequently add 3 µL of the 1:10 compound dilution to column 2

   c. Make a 1:3 serial dilution of the compounds by transferring 50 µL from one column to the next column (columns 2-9) with tip change in columns 4 and 7.

3. Adding Virus:
   a. Prepare virus (WNV NY99) to the appropriate dilution in ASSAY MEDIUM
      i. Titer of the WNV stock is 8.89E+07 TCID50/ml: prepare 1:125 (final dilution 1:500)
   b. Add 50 µL of the virus solution to columns 2-10
   c. Add 50 µL of assay medium to columns 2-11.
   (thus, 100 µL of cells containing 1 µM CP100356 + 50 µL cpd + 50 µL virus ® final concentration CP100356 = 0.5 µM)
      i. In case of insufficient cells: medium instead of cells can be added to the border wells

4. Incubate plates (37°C / 5% $CO_2$)

5. On day 7 post-infection, determine cell viability using MTS (see Jochmans D et al. J Virol Methods. 2012 Aug;183(2):176-9. doi: 10.1016/j.jviromet.2012.04.011)

7. **In vitro ADMET**

In vitro ADMET experiments for this work were performed at Concept Life Sciences (United Kingdom) and Bienta (Ukraine). Links to the protocols that were employed are below:

MDR1-MDCKII cell permeability: dx.doi.org/10.17504/protocols.io.n2bvjne6ngk5/v1

Microsomal stability (human + mouse): dx.doi.org/10.17504/protocols.io.5qpvokdb9l4o/v1

Kinetic solubility: dx.doi.org/10.17504/protocols.io.j8nlk8y41l5r/v1

LogD: dx.doi.org/10.17504/protocols.io.e6nvw14kdlmk/v1

Plasma protein binding (mouse): dx.doi.org/10.17504/protocols.io.dm6gp9yb5vzp/v1

Fetal calf serum protein binding: dx.doi.org/10.17504/protocols.io.e6nvwbzk2vmk/v1.

To estimate unbound fraction in 2% and 10% FCS for cell assays in VeroE6, SH-SY5Y, and JEG-3 cells, formula 1 was used:

$$f_{u,\text{medium}} = \frac{1}{(\text{protein\%})(1/f_u - 1) + 1} \quad (1)$$

In 100% FCS, ASAP-0036543 $F_u$ = 0.114 (measured at Concept Life Sciences)

In 2% FCS (VeroE6 + SH-SY5Y assays), $F_{u,calc}$ = 0.865

In 10% FCS (JEG-3 assay), $F_{u,calc}$ = 0.563

See refs for more information: [2,3]

#### 8. ASAP-0027808 human protease selectivity panel

The inhibitory activity of ASAP-0027808 was evaluated against 30 human proteases (Eurofins Panlabs, 6 Research Park Dr., St. Charles, MO) in single-point format.

| Cat # | Assay Name | Batch* | Spec. | Rep. | Conc. | % Inh. |
|---|---|---|---|---|---|---|
| 107371 | Peptidase Bleomycin Hydrolase (BLMH) | 522649 | hum | 2 | 10 µM | -2 |
| 108020 | Peptidase CAN1 (CANPL1 Calpain-1) | 522930 | hum | 2 | 10 µM | 15 |
| 113810 | Peptidase, Kallikrein, Plasma | 522657 | hum | 2 | 10 µM | 6 |
| 165060 | Peptidase Tissue Plasminogen Activator (tPA) | 522928 | hum | 2 | 10 µM | -3 |
| 199011 | Peptidase, CASP1 (Caspase 1) | 522877 | hum | 2 | 10 µM | 0 |
| 163100 | Peptidase, CASP2 (Caspase 2) | 522874 | hum | 2 | 10 µM | 3 |
| 163240 | Peptidase, CASP4 (Caspase 4) | 522875 | hum | 2 | 10 µM | -1 |
| 163280 | Peptidase, CASP5 (Caspase 5) | 522876 | hum | 2 | 10 µM | -3 |
| 113310 | Peptidase, Chymase | 522869 | hum | 2 | 10 µM | -2 |
| 113400 | Peptidase, Chymotrypsin | 522870 | hum | 2 | 10 µM | -7 |
| 112250 | Peptidase, CTSB (Cathepsin B) | 522421 | hum | 2 | 10 µM | 9 |
| 112310 | Peptidase, CTSC (Cathepsin C) | 522422 | hum | 2 | 10 µM | 6 |
| 112510 | Peptidase, CTSG (Cathepsin G) | 522423 | hum | 2 | 10 µM | -1 |
| 112550 | Peptidase, CTSH (Cathepsin H) | 522644 | hum | 2 | 10 µM | -12 |
| 112600 | Peptidase, CTSK (Cathepsin K) | 522424 | hum | 2 | 10 µM | -2 |
| 112650 | Peptidase, CTSL (Cathepsin L) | 522645 | hum | 2 | 10 µM | 3 |
| 112800 | Peptidase, CTSL2 (Cathepsin L2) | 523114 | hum | 2 | 10 µM | 17 |
| 112750 | Peptidase, CTSS (Cathepsin S) | 522646 | hum | 2 | 10 µM | -4 |
| 112900 | Peptidase, CTSZ (Cathepsin Z) | 522648 | hum | 2 | 10 µM | 3 |
| 199007 | Peptidase, Dipeptidyl Peptidase 4 (DPP4, DPP IV) | 522652 | hum | 2 | 10 µM | -1 |
| 166050 | Peptidase, ELA1 (Pancreatic Elastase 1) | 522993 | pig | 2 | 10 µM | -2 |
| 166020 | Peptidase, ELA2 (Neutrophil Elastase 2) | 523050 | hum | 2 | 10 µM | 7 |
| 113510 | Peptidase, Factor VIIa | 522655 | hum | 2 | 10 µM | -7 |
| 113610 | Peptidase, Factor Xa | 522656 | hum | 2 | 10 µM | -5 |
| 164510 | Peptidase Plasmin | 522925 | hum | 2 | 10 µM | -1 |
| 115900 | Peptidase, PLAU (Urokinase) | 522884 | hum | 2 | 10 µM | 21 |
| 164410 | Peptidase, Prolyl Oligopeptidase (POP) | 523048 | hum | 2 | 10 µM | 0 |
| 165000 | Peptidase, Thrombin | 522885 | hum | 2 | 10 µM | -2 |
| 165100 | Peptidase, Trypsin | 522886 | hum | 2 | 10 µM | 1 |
| 165200 | Peptidase, Tryptase | 523049 | hum | 2 | 10 µM | -5 |

#### 9. Murine pharmacokinetics

In vivo experiments and formulation of ASAP-036543 were conducted at WuXi Apptec (Shanghai, CN). Studies were conducted in accordance with the WuXi AppTec IACUC standard animal procedures along with the IACUC guidelines that are in compliance with the Animal Welfare Act, the Guide for the Care and Use of Laboratory Animals.

In brief, healthy animals were acclimated at least 3 days upon arrival before beginning the studies. Environment controls will be set to maintain a temperature range of 20-26°C, a relative humidity range of 30 to 70% with no less than 15 air changes/hour, and a 12-hour light/12-hour dark cycle. Animals were group-housed in a polysulfone cage with certified aspen shaving bedding or corncob bedding. Animals were offered water ad libitum and certified rodent breeding and growth diet ad libitum beginning four hours after dosing.

For PO studies, 6-8 week old female C57BL/6J mice were fasted overnight and dosed orally by gavage tube. Food was returned 4 hours after dosing. For the IV study, a solution of the test compound was delivered intravenously by bolus injection via the tail vein.

##### Analytical and collection methods

Each blood collection (about 0.02 mL per time point) was performed via saphenous vein or other suitable blood collection site of each animal into commercial microcentrifuge tubes containing K2-EDTA as anti-coagulant and placed on wet ice until centrifugation. Samples were centrifuged (3200×g for 10 minutes at 2 to 8°C) within one hour of collection. The plasma samples were transferred into labeled polypropylene micro-centrifuge tubes and stored frozen in a freezer set to maintain -60°C or lower until bio-analysis.

A calibration curve with at least 6 non-zero calibration standards was applied for each batch including LLOQ. Plasma concentration versus time was plotted and analyzed by non-compartmental approaches using Phoenix WinNonlin 6.3. Related PK parameters were be calculated according to dosing route, e.g. Cmax, Tmax for extravascular administration, and T½, $AUC_{(0\text{-}t)}$, $AUC_{(0\text{-}inf)}$, $MRT_{(0\text{-}t)}$, $MRT_{(0\text{-}inf)}$ for all routes.

##### Formulation procedure, batch 3 (single dose PK, Fig 9A):

ASAP-0036543 (1 HCl salt) was dosed as a clear solution in 9:1 water / solutol at concentrations of 1-60 mg/mL. In brief, solid compound was added to a glass vial and the appropriate quantity of solutol was added. The mixture was stirred for 10 min at 1700 rpm, 37℃ to get a clear solution. The appropriate quantity of water was then added into the vial and vortexed for 2 min to get a clear solution. The solution was then filtered through a 0.22 μm PVDF membrane prior to dosing.

##### Formulation procedure, batch 8 (used for in vivo efficacy study, Fig 9B-C):

ASAP-0036543 (freebase) was formulated at 30 mg/mL and dosed at 5 mL/kg, for a final dose of 150 mg/kg, as follows: solid compound was added to a glass vial and the appropriate quantity of solutol was added. The mixture was stirred for 30 min at 1700 rpm, 37℃ to get a nearly clear solution with fine particles. The appropriate quantity of citrate buffer was then added into the vial and vortexed for 10 min to get a clear solution. The final pH was 6.7 when formulated at 60 mg/mL. The solution was then filtered through a 0.22 μm PVDF membrane and administered as an oral gavage at 10 mL/kg.

The concentration of the test compound in dose formulation samples was determined by the LC/UV or LC/MS/MS.

**Pharmacokinetic Data**

Data from the single 150 mg/kg PO experiment is shown below (values in micromolar).

| Time (h) | M01 | M02 | M03 | Mean | Mean Unbound |
|---|---|---|---|---|---|
| **0.083** | 3.92 | 1.32 | 1.07 | 2.10 | 0.20 |
| **0.25** | 5.70 | 2.89 | 3.42 | 4.00 | 0.38 |
| **0.5** | 7.74 | 9.89 | 4.73 | 7.45 | 0.72 |
| **1** | 9.59 | 11.9 | 6.16 | 9.22 | 0.88 |
| **2** | 12.9 | 12.1 | 9.92 | 11.6 | 1.12 |
| **5** | 10.2 | 5.68 | 5.13 | 7.00 | 0.67 |
| **8** | 3.11 | 1.55 | 2.82 | 2.49 | 0.24 |
| **24** | 0.120 | 0.0215 | 0.0448 | 0.0621 | 0.01 |

Data from single 2mg/kg IV experiment are shown below (unless otherwise specified, values in micromolar)

| Time (h) | M01 | M02 | M03 | Mean |
|---|---|---|---|---|
| **0.083** | 1.03 | 1.11 | 0.925 | 1.02 |
| **0.25** | 0.632 | 0.690 | 0.638 | 0.653 |
| **0.5** | 0.375 | 0.507 | 0.429 | 0.437 |
| **1** | 0.215 | 0.246 | 0.261 | 0.241 |
| **2** | 0.0980 | 0.120 | 0.131 | 0.116 |
| **5** | 0.0141 | 0.0263 | 0.0316 | 0.0240 |
| **8** | 0.00379 | 0.00794 | 0.0145 | 0.00874 |
| **24** | BQL | BQL | BQL | ND |
| **PK Parameters** | **M01** | **M02** | **M03** | **Mean** |
| **Rsq_adj** | 0.982 | 0.991 | 0.956 | -- |
| **No. points used for $T_{1/2}$** | 4.00 | 3.00 | 4.00 | ND |
| **$C_0$ (umol/L)** | 1.31 | 1.41 | 1.11 | 1.28 |
| **$T_{1/2}$ (h)** | 1.20 | 1.53 | 1.69 | 1.47 |
| **$Vd_{ss}$ (L/kg)** | 7.35 | 7.24 | 9.02 | 7.87 |
| **Cl(mL/min/kg)** | 99.8 | 80.3 | 79.6 | 86.6 |
| **$T_{last}$ (h)** | 8.00 | 8.00 | 8.00 | 8.00 |
| **$AUC_{0-last}$ (h*umol/L)** | 0.802 | 0.987 | 0.978 | 0.922 |
| **$AUC_{0-inf}$ (h*umol/L)** | 0.809 | 1.00 | 1.01 | 0.940 |
| **$MRT_{0-last}$ (h)** | 1.16 | 1.35 | 1.58 | 1.36 |
| **$MRT_{0-inf}$ (h)** | 1.23 | 1.50 | 1.89 | 1.54 |
| **$AUC_{Extra}$ (%)** | 0.813 | 1.75 | 3.49 | 2.02 |
| **$AUMC_{Extra}$ (%)** | 6.44 | 11.9 | 19.3 | 12.5 |

#### 10. In vivo efficacy

The in vivo efficacy study described in Fig 9C was performed at KU LEUVEN in AG129 mice infected with ZIKV strain MR766 using modifications of a previously published protocol (Zmurko *et al*., PLoS Negl Trop Dis. 2016; 10(5):e0004695. doi:10.1371/journal.pntd.0004695; PMID: 27163257).

**Cells and virus**

ZIKV (strain MR766) was generated synthetically from overlapping DNA fragments (IDT Integrated DNA Technologies), based on the sequence available in GenBank (accession number DQ859059) and assembled by overlap extension PCR. Virus stocks were generated on Vero E6 cell cultures (African Green monkey kidney cells, Vero C1008; ATCC CRL-1586) grown in Dulbecco's Modified Eagle Medium (DMEM) supplemented with 10% fetal calf serum (FCS) and 1% sodium bicarbonate at 37 °C in a $CO_2$ incubator. Virus stocks were grown in Minimum Essential Medium (MEM) supplemented with 2% FCS, 1% non-essential amino acids (NEAA), 2 mM L-glutamine and 1% sodium bicarbonate at 37 °C in a $CO_2$ incubator. At the time ZIKV caused a complete cytopathic effect (CPE) [d5-d7 post infection; pi], the supernatant was harvested and viral titers were determined by plaque assay using BHK-21 cells (baby hamster kidney cells; ATCC CCL-10), essentially as described previously (Kum *et al*., npj Vaccines 2023; doi:10.1038/s41541-023-00699-7; PMID: 37433816). The different cell types as well as the ZIKV stock tested negative for mycoplasma.

**ZIKV infection/viremia model in AG129 mice (Fig 9C)**

7-15 weeks old male and female AG129 mice of 20-25 grams (129/Sv mice with knockouts in both IFNα/β and IFNγ receptor genes) were bred in-house. Breeding couples of AG129 mice were purchased from Marshall BioResources. The specific pathogen-free status of the mice was regularly checked at the KU Leuven animal facility of the Rega Institute. Mice were housed in individually ventilated isolator cages (IsoCage N Biocontainment System, Techniplast) at a temperature of 21°C, humidity of 55%, and subjected to 12:12 dark/light cycles. They had access to food and water *ad libitum*, and their cages were enriched with cotton and cardboard play tunnels. Housing conditions and experimental procedures were approved by the Ethical Committee Dierproeven (animal experimentation) of KU Leuven (License DMIT-110/2025), following institutional guidelines approved by the Federation of European Laboratory Animal Science Associations. The animal work was conducted under Biosafety Level 2 conditions.

Study groups consisted of 5 males and 5 females aged 7-15 weeks and weighing 20-25 grams. At the start of the study, the dosing group was given the first dose of ASAP-0035643. Two hours after the start of the study, all mice were inoculated intraperitoneally (ip; 200 µL) with $1\times10^4$ PFU ZIKV (strain MR766). Mice were observed daily for body weight change and the development of virus-induced disease. In case of a body weight loss of ≥20% and/or severe illness OR at study endpoint, mice were euthanized by intraperitoneal injection of 50 µL dolethal.

Formulation in 10% Solutol in 50mM citrate buffer was used, pH 3, prepared for oral administration at 150 mg/kg/dose. Compound was administered twice a day (BID); Fresh formulations were prepared prior to each administration.

The schedule of the study is described below.

For dosing group:

| **Day** | 0 | 0 | 0 | 1 | 1 | 2 | 2 | 3 |
|---|---|---|---|---|---|---|---|---|
| **Time** | 8 am (T0hr) | 10 am | 6 pm | 8 am | 6 pm | 8 am | 6 pm | 11 am (T+75h) |
| **Action** | Study start: dose compound | Infect with ZIKV MR766 | Dose compound | Dose compound | Dose compound | Dose compound | Dose compound | Euthanize |

For infected control group:

| **Day** | 0 | 0 | | | | | | 3 |
|---|---|---|---|---|---|---|---|---|
| **Time** | 8 am (T0hr) | 10 am | | | | | | 11 am (T+75h) |
| **Action** | Study start | Infect with ZIKV MR766 | | | | | | Euthanize |

**Sample collection and processing**

RNA isolation and quantitative RT-PCR: RNA was isolated from 75 µl plasma using the NucleoSpin RNA virus kit (Filter Service, Germany), according to the manufacturer's protocol. RT-qPCR was performed on a LightCycler96 platform (Roche). During RT-qPCR the ZIKV NS1 region (nucleotides 2472–2565) was amplified using primers 5'-TGA CTC CCC TCG TAG ACT G-3' and 3'-CTC TCC TTC CAC TGA TTT CCA C-5' and a Double-Quenched Probe 5'-6-FAM/AGA TCC CAC /ZEN/AAA TCC CCT CTT CCC/3'IABkFQ/ (Integrated DNA Technologies, IDT). Viral RNA was quantified using serial dilutions of a standard curve consisting of a synthesized gene block containing 145 bp of ZIKV NS1 (nucleotides 2456–2603): 5'-GGT ACA AGT ACC ATC CTG ACT CCC CTC GTA GAC TGG CAG CAG CCG TTA AGC AAG CTT GGG AAG AGG GGA TTT GTG GGA TCT CCT CTG TTT CTA GAA TGG AAA ACA TAA TGT GGA AAT CAG TGG AAG GAG AGC TCA ATG CAA TCC TAG-3' (Integrated DNA Technologies).

**Blood draws and PK quantitation (Fig 9B)**

During the course of the in vivo efficacy study described in Fig. 9C, blood was collected from tail veins every 24 h into $K_2$EDTA-coated tubes: ~20 µL plasma for PK, inactivated 30 min at 56 °C before freezing at -80 °C (12 h after 2nd dose, immediately prior to 3rd dose), 48 h (12 h after 4th dose, immediately prior to 5th dose), and 72 h (12 h after 6th dose, 3 h before study conclusion). 30% isopropanol in water was used as the homogenization buffer.

Samples were stored at -80 °C until shipment to Frontage Labs, Exton, PA, US. ASAP-0036543 in mouse plasma was quantitated at Frontage.

10 µL of control matrix was added to separate wells of a 96 well plate for STDs, QCs, and Blanks. To the control matrix, 10 µL of the appropriate neat solution was added to STDs and QCs. (10 µL of diluent was added to blanks).

For study samples, 10 µL of each sample was added to separate wells of the 96-well plate and a 10 µL aliquot of diluent was added to each study sample. To all samples, 200 µL of internal standard (200 ng/mL Warfarin/Verapamil) solution in ACN was added. The plate was shaken vigorously for 20 minutes in a Qiagen TissueLyser II at 15.0 Hz and then centrifuged for 10 minutes at 4000 rpm (Sorvall Legend X1R centrifuge) at 15 C. After centrifugation, 100 µL aliquots of supernatant were transferred to a new 96-well plate containing 100 µL of water in each well. The plate was vortexed for approximately 10 minutes and then aliquots were injected on LC/MS/MS.

| | | |
|---|---|---|
| Instrumentation: | | |
| LC System: System #902 | | |
| HPLC Pump: Shimadzu LC-30AD | | |
| Autosampler: Shimadzu SIL-30AC MP | | |
| Column Oven: Shimadzu CTO-20AC | | |
| HPLC Conditions: | | |
| Mobile Phase A = 0.1% Formic Acid in Water | | Injection Volume = 0.6 µL |
| Mobile Phase B = 0.1% Formic Acid in ACN | | Column = ACE 5 C8, 2.1 x 50 mm, 5 µm (ACE) |
| Column Oven Temperature = Off | | Flow Rate = 0.65 mL/min |
| Gradient: | | |

| Time (min.) | %B |
|---|---|
| 0.00 | 5 |
| 0.80 | 5 |
| 2.20 | 95 |
| 3.50 | 95 |
| 3.60 | 5 |
| 4.50 | 5 |

| | |
|---|---|
| MS/MS System: System #899 | |
| Sciex Triple Quad 6500+ (S/N DZ221761812) | |
| MS Conditions: | |
| Scan Type = MRM | EP = 10 |
| Polarity = Positive | IS = 3500 |
| Ion Source = ESI | NC = NA |
| CAD = 10 | GS1/GS2 = 50/50 |
| CUR = 20 | TEM = 550 |

**Weight change in AG129 mice during efficacy study**

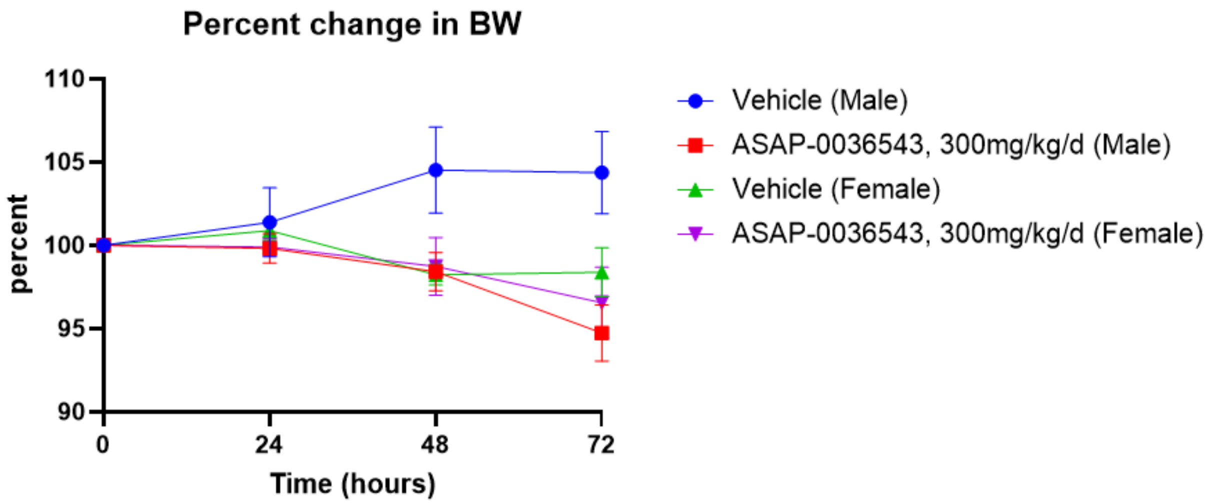

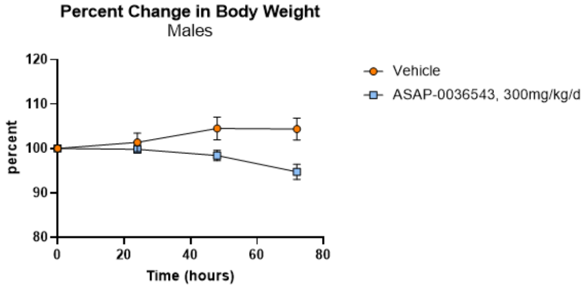

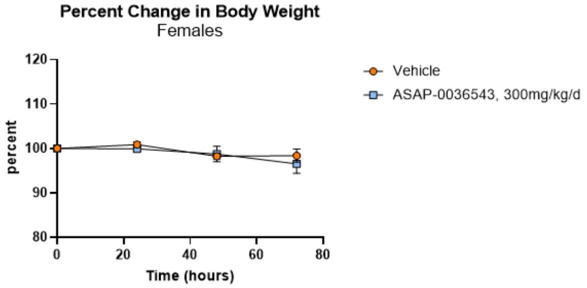

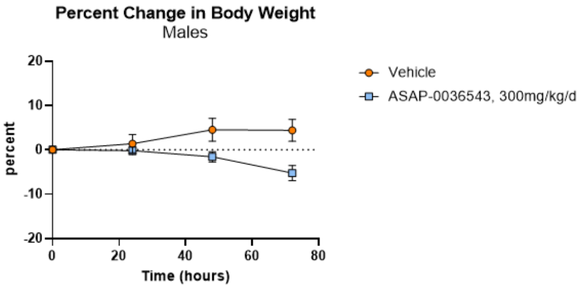

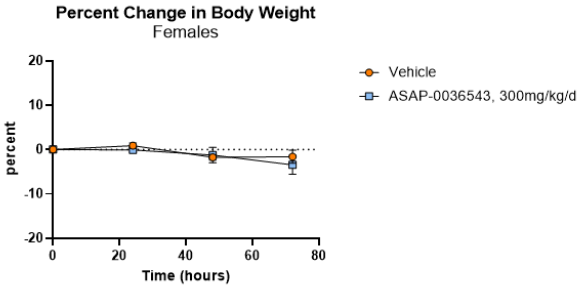

| | | d = -3 | d = 0 | d = 1 | d = 2 | d = 3 | |
|---|---|---|---|---|---|---|---|
| | Weight (g) | | | | | | |
| | ORAL GAVAGE, all groups | | *Treat* | *Treat* | *Treat* | *Treat* | Day of euthanasia |
| | | | | | | | %Weight loss |
| MALES | GROEP 1a - VEHICLE | | | no PK | no PK | no PK | |
| | | 32.7 | 33.1 | 32.5 | Died | | 1.8 |
| | | 30.8 | 31.5 | 31.1 | 32.5 | 32.6 | -3.5 |
| | | 32.2 | 30.0 | 32.0 | 33.1 | 32.7 | -9.0 |
| | | 31.3 | 32.0 | 31.2 | 31.4 | 31.3 | 2.2 |
| | | 26.0 | 26.3 | 27.0 | 28.0 | 28.2 | -7.2 |
| FEMALES | GROEP 1b - VEHICLE | | | no PK | no PK | no PK | |
| | | 22.5 | 22.8 | 23.3 | 22.9 | 22.2 | 2.6 |
| | | 24.5 | 25.1 | 25.7 | 24.8 | 25.1 | 0.0 |
| | | 23.7 | 23.8 | 23.7 | 23.1 | 23.8 | 0.0 |
| | | 25.8 | 27.0 | 27.2 | 26.4 | 27.4 | -1.5 |
| | | 24.9 | 24.7 | 24.6 | 24.0 | 23.0 | 6.9 |
| MALES | ASAP-0036543 - 150 mg/kg/d | | | *Blood-PK* | *Blood-PK* | *Blood-PK* | |
| | | 27.7 | 28.6 | 28.2 | 27.6 | 27.0 | 5.6 |
| | | 30.2 | 31.6 | 31.2 | 30.7 | 29.1 | 7.9 |
| | | 27.6 | 27.8 | 27.2 | 26.8 | 25.3 | 9.0 |
| | | 27.9 | 28.9 | 29.5 | 29.6 | 29.1 | -0.7 |
| | | 26.7 | 27.0 | 27.5 | 26.9 | 25.8 | 4.4 |
| FEMALES | ASAP-0036543 - 150 mg/kg/d | | | no PK | no PK | no PK | |
| | | 24.2 | 23.3 | 23.4 | 23.0 | 22.4 | 3.9 |
| | | 22.2 | 23.5 | 23.3 | 23.2 | 22.8 | 3.0 |
| | | 22.1 | 22.5 | 22.1 | 20.8 | 20.0 | 11.1 |
| | | 21.4 | 22.6 | 22.8 | 23.1 | 22.4 | 0.9 |
| | | 24.7 | 24.3 | 24.5 | 24.7 | 24.7 | -1.6 |

Day of euthanasia

#### 11. Safety profiling of ASAP-0036543

Safety profiling of ASAP-0036543 was conducted at Eurofins

### 6. COMPOUNDS

#### 6.1. Test Compounds

| Client Compound ID | Compound ID | Reference Number | Batch Number | FW | MW | Purity | Received Form | Stock solution | Flag |
|---|---|---|---|---|---|---|---|---|---|
| ASAP-0036543 | 100078696-1 | Z9389150425 | 7.00 | 486.31 | 413.38 | 100.0 | Powder | 1.E-02 M DMSO | - |

*FW: Formula Weight - MW: Molecular Weight*

#### 6.2. Reference Compounds

In each experiment and if applicable, the respective reference compound was tested concurrently with ASAP-0036543, and the data were compared with historical values determined at Eurofins. The experiment was accepted in accordance with Eurofins validation Standard Operating Procedure.

| Assay | 1.0E-05 M |
|---|---|
| 5-$HT_{2A}$(*h*) (agonist radioligand) | 80.6% |
| $beta_1$(*h*) (agonist radioligand) | 79.3% |
| Cav1.2 (L-type) Human Calcium Ion Channel Binding (Dihydropyridine Site) LeadHunter Assay - FR | 67.7% |
| Potassium Channel hERG (human)- [3H] Dofetilide | 83.2% |
| Sodium Channel Site2 (Non-selective) Rat Ion Channel Batrachotoxin Mass Spectrometry Binding | 86.5% |
| COX1(h) | 54.6% |
| COX2(h) | 57.2% |

#### 7.1. *In Vitro* Pharmacology: Binding Assays

##### 7.1.1. Test Compound Results

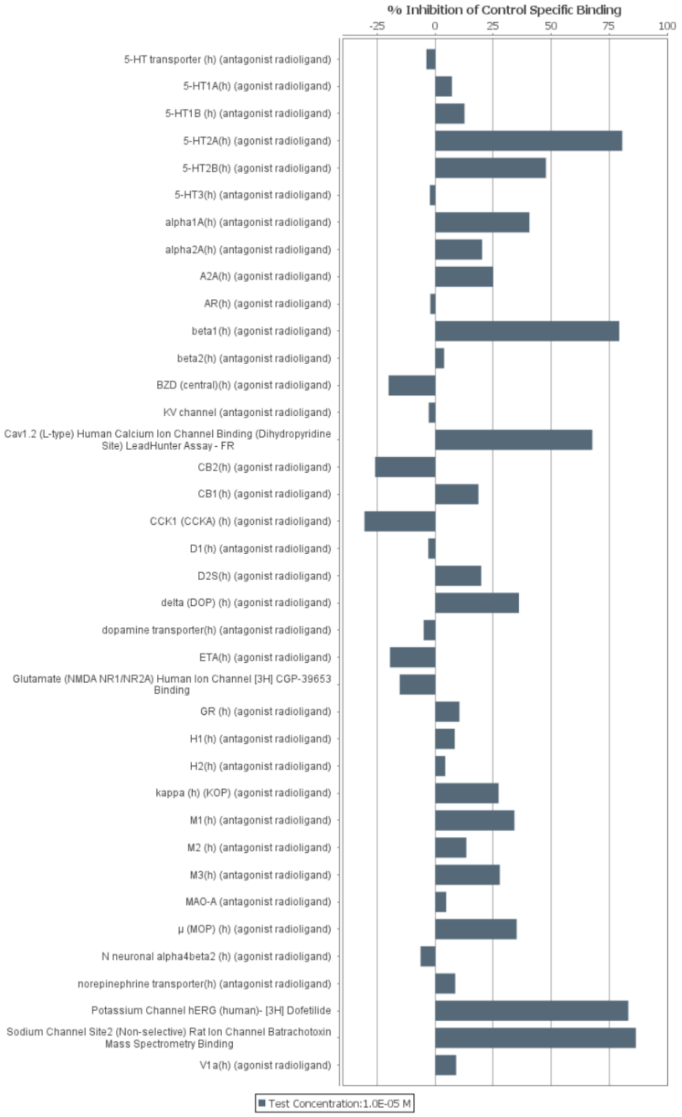

**Figure 1. Histogram for ASAP-0036543**

| Compound I.D. | Client Compound I.D. | Test Concentration | % Inhibition of Control Specific Binding | | |
|---|---|---|---|---|---|
| | | | 1st | 2nd | Mean |
| **5-HT transporter *(h)* (antagonist radioligand)** | | | | | |
| 100078696-1 | ASAP-0036543 | 1.0E-05 M | -5.2 | -2.5 | -3.8 |
| **5-HT$_{1A}$*(h)* (agonist radioligand)** | | | | | |
| 100078696-1 | ASAP-0036543 | 1.0E-05 M | 11.7 | 2.7 | 7.2 |
| **5-HT$_{1B}$ *(h)* (antagonist radioligand)** | | | | | |
| 100078696-1 | ASAP-0036543 | 1.0E-05 M | 8.8 | 16.4 | 12.6 |
| **5-HT$_{2A}$*(h)* (agonist radioligand)** | | | | | |
| 100078696-1 | ASAP-0036543 | 1.0E-05 M | 79.6 | 81.7 | **80.6** |
| **5-HT$_{2B}$*(h)* (agonist radioligand)** | | | | | |
| 100078696-1 | ASAP-0036543 | 1.0E-05 M | 48.9 | 46.6 | 47.7 |
| **5-HT$_{3}$*(h)* (antagonist radioligand)** | | | | | |
| 100078696-1 | ASAP-0036543 | 1.0E-05 M | 1.8 | -6.4 | -2.3 |
| **alpha$_{1A}$*(h)* (antagonist radioligand)** | | | | | |
| 100078696-1 | ASAP-0036543 | 1.0E-05 M | 40.4 | 40.8 | 40.6 |
| **alpha$_{2A}$*(h)* (antagonist radioligand)** | | | | | |
| 100078696-1 | ASAP-0036543 | 1.0E-05 M | 19.9 | 20.5 | 20.2 |
| **A$_{2A}$*(h)* (agonist radioligand)** | | | | | |
| 100078696-1 | ASAP-0036543 | 1.0E-05 M | 32.1 | 18.0 | 25.0 |
| **AR(h) (agonist radioligand)** | | | | | |
| 100078696-1 | ASAP-0036543 | 1.0E-05 M | 3.4 | -7.6 | -2.1 |
| **beta$_{1}$*(h)* (agonist radioligand)** | | | | | |
| 100078696-1 | ASAP-0036543 | 1.0E-05 M | 79.9 | 78.6 | **79.3** |
| **beta$_{2}$*(h)* (antagonist radioligand)** | | | | | |
| 100078696-1 | ASAP-0036543 | 1.0E-05 M | 4.1 | 3.6 | 3.8 |
| **BZD (central)(h) (agonist radioligand)** | | | | | |
| 100078696-1 | ASAP-0036543 | 1.0E-05 M | -15.9 | -24.4 | -20.1 |
| **K$_{V}$ channel (antagonist radioligand)** | | | | | |
| 100078696-1 | ASAP-0036543 | 1.0E-05 M | -2.1 | -3.4 | -2.8 |
| **Cav1.2 (L-type) Human Calcium Ion Channel Binding (Dihydropyridine Site) LeadHunter Assay - FR** | | | | | |
| 100078696-1 | ASAP-0036543 | 1.0E-05 M | 68.0 | 67.4 | **67.7** |
| **CB$_{2}$*(h)* (agonist radioligand)** | | | | | |
| 100078696-1 | ASAP-0036543 | 1.0E-05 M | -16.5 | -35.4 | -25.9 |
| **CB$_{1}$*(h)* (agonist radioligand)** | | | | | |
| 100078696-1 | ASAP-0036543 | 1.0E-05 M | 17.3 | 19.8 | 18.6 |
| **CCK$_{1}$ (CCK$_{A}$) *(h)* (agonist radioligand)** | | | | | |
| 100078696-1 | ASAP-0036543 | 1.0E-05 M | -30.5 | -30.5 | -30.5 |
| **D$_{1}$*(h)* (antagonist radioligand)** | | | | | |
| 100078696-1 | ASAP-0036543 | 1.0E-05 M | 1.1 | -7.1 | -3.0 |
| **D$_{2S}$*(h)* (agonist radioligand)** | | | | | |
| 100078696-1 | ASAP-0036543 | 1.0E-05 M | 17.5 | 22.1 | 19.8 |
| **delta (DOP) *(h)* (agonist radioligand)** | | | | | |
| 100078696-1 | ASAP-0036543 | 1.0E-05 M | 39.3 | 32.9 | 36.1 |
| **dopamine transporter*(h)* (antagonist radioligand)** | | | | | |
| 100078696-1 | ASAP-0036543 | 1.0E-05 M | -4.7 | -5.4 | -5.0 |
| **ET$_{A}$*(h)* (agonist radioligand)** | | | | | |
| 100078696-1 | ASAP-0036543 | 1.0E-05 M | -17.5 | -21.6 | -19.5 |
| **Glutamate (NMDA NR1/NR2A) Human Ion Channel [3H] CGP-39653 Binding** | | | | | |
| 100078696-1 | ASAP-0036543 | 1.0E-05 M | -17.4 | -13.2 | -15.3 |
| **GR *(h)* (agonist radioligand)** | | | | | |
| 100078696-1 | ASAP-0036543 | 1.0E-05 M | 15.2 | 5.7 | 10.4 |
| **H$_{1}$*(h)* (antagonist radioligand)** | | | | | |
| 100078696-1 | ASAP-0036543 | 1.0E-05 M | 13.0 | 3.7 | 8.4 |

| Compound I.D. | Client Compound I.D. | Test Concentration | % Inhibition of Control Specific Binding | | |
|---|---|---|---|---|---|
| | | | 1st | 2nd | Mean |
| **$H_2$*(h)* (antagonist radioligand)** | | | | | |
| 100078696-1 | ASAP-0036543 | 1.0E-05 M | 1.6 | 7.0 | 4.3 |
| **kappa *(h)* (KOP) (agonist radioligand)** | | | | | |
| 100078696-1 | ASAP-0036543 | 1.0E-05 M | 31.7 | 22.9 | 27.3 |
| **$M_1$*(h)* (antagonist radioligand)** | | | | | |
| 100078696-1 | ASAP-0036543 | 1.0E-05 M | 34.8 | 33.4 | 34.1 |
| **$M_2$ *(h)* (antagonist radioligand)** | | | | | |
| 100078696-1 | ASAP-0036543 | 1.0E-05 M | 13.8 | 13.0 | 13.4 |
| **$M_3$*(h)* (antagonist radioligand)** | | | | | |
| 100078696-1 | ASAP-0036543 | 1.0E-05 M | 29.7 | 25.9 | 27.8 |
| **MAO-A (antagonist radioligand)** | | | | | |
| 100078696-1 | ASAP-0036543 | 1.0E-05 M | 5.7 | 3.6 | 4.7 |
| **μ (MOP) *(h)* (agonist radioligand)** | | | | | |
| 100078696-1 | ASAP-0036543 | 1.0E-05 M | 38.2 | 31.9 | 35.1 |
| **N neuronal alpha4beta2 *(h)* (agonist radioligand)** | | | | | |
| 100078696-1 | ASAP-0036543 | 1.0E-05 M | -7.6 | -5.1 | -6.3 |
| **norepinephrine transporter*(h)* (antagonist radioligand)** | | | | | |
| 100078696-1 | ASAP-0036543 | 1.0E-05 M | 6.1 | 11.1 | 8.6 |
| **Potassium Channel hERG (human)- [3H] Dofetilide** | | | | | |
| 100078696-1 | ASAP-0036543 | 1.0E-05 M | 84.5 | 81.9 | **83.2** |
| **Sodium Channel Site2 (Non-selective) Rat Ion Channel Batrachotoxin Mass Spectrometry Binding** | | | | | |
| 100078696-1 | ASAP-0036543 | 1.0E-05 M | 89.0 | 84.0 | **86.5** |
| **$V_{1a}$*(h)* (agonist radioligand)** | | | | | |
| 100078696-1 | ASAP-0036543 | 1.0E-05 M | 9.9 | 8.1 | 9.0 |

##### 7.1.2. Reference Compound Results

| Compound I.D. | $IC_{50}$(M) | $K_i$(M) | nH |
|---|---|---|---|
| **5-HT transporter *(h)* (antagonist radioligand)** | | | |
| imipramine | 3.9E-09 M | 1.8E-09 M | 0.9 |
| **$5\text{-}HT_{1A}$*(h)* (agonist radioligand)** | | | |
| 8-OH-DPAT | 9.3E-10 M | 4.6E-10 M | 1.0 |
| **$5\text{-}HT_{1B}$ *(h)* (antagonist radioligand)** | | | |
| Serotonine | 1.3E-07 M | 6.0E-08 M | 0.7 |
| **$5\text{-}HT_{2A}$*(h)* (agonist radioligand)** | | | |
| (±)DOI | 2.8E-10 M | 2.1E-10 M | 1.0 |
| **$5\text{-}HT_{2B}$*(h)* (agonist radioligand)** | | | |
| (±)DOI | 4.5E-09 M | 2.2E-09 M | 0.7 |
| **$5\text{-}HT_{3}$*(h)* (antagonist radioligand)** | | | |
| MDL 72222 | 5.4E-09 M | 3.7E-09 M | 1.0 |
| **$alpha_{1A}$*(h)* (antagonist radioligand)** | | | |
| WB 4101 | 4.6E-10 M | 2.3E-10 M | 1.5 |
| **$alpha_{2A}$*(h)* (antagonist radioligand)** | | | |
| yohimbine | 7.1E-09 M | 3.2E-09 M | 1.1 |
| **$A_{2A}$*(h)* (agonist radioligand)** | | | |
| NECA | 5.9E-08 M | 4.8E-08 M | 1.7 |
| **AR(h) (agonist radioligand)** | | | |
| testosterone | 1.0E-08 M | 3.8E-09 M | 1.0 |
| **$beta_1$*(h)* (agonist radioligand)** | | | |
| atenolol | 2.5E-07 M | 1.4E-07 M | 1.0 |
| **$beta_2$*(h)* (antagonist radioligand)** | | | |
| ICI 118551 | 9.4E-10 M | 3.1E-10 M | 1.1 |
| **BZD (central)(h) (agonist radioligand)** | | | |
| Diazepam | 5.2E-08 M | 4.6E-08 M | 1.5 |
| **$K_V$ channel (antagonist radioligand)** | | | |
| α-dendrotoxin | 1.2E-10 M | 9.8E-11 M | 1.1 |
| **Cav1.2 (L-type) Human Calcium Ion Channel Binding (Dihydropyridine Site) LeadHunter Assay - FR** | | | |
| Nitrendipine | 7.3E-10 M | 2.3E-10 M | 0.9 |
| **$CB_2$*(h)* (agonist radioligand)** | | | |
| WIN 55212-2 | 8.5E-10 M | 5.5E-10 M | 1.2 |
| **$CB_1$*(h)* (agonist radioligand)** | | | |
| CP 55940 | 3.3E-09 M | 1.0E-09 M | 1.0 |
| **$CCK_1$ ($CCK_A$) *(h)* (agonist radioligand)** | | | |
| CCK-8s | 1.0E-10 M | 7.7E-11 M | 1.3 |
| **$D_1$*(h)* (antagonist radioligand)** | | | |
| SCH 23390 | 4.4E-10 M | 1.8E-10 M | 1.2 |
| **$D_{2S}$*(h)* (agonist radioligand)** | | | |
| 7-OH-DPAT | 5.1E-09 M | 2.1E-09 M | 0.7 |
| **delta (DOP) *(h)* (agonist radioligand)** | | | |
| DPDPE | 2.8E-09 M | 1.5E-09 M | 0.9 |
| **dopamine transporter*(h)* (antagonist radioligand)** | | | |
| BTCP | 1.4E-08 M | 7.6E-09 M | 1.0 |
| **$ET_A$*(h)* (agonist radioligand)** | | | |
| endothelin-1 | 4.9E-11 M | 2.4E-11 M | 1.8 |
| **Glutamate (NMDA NR1/NR2A) Human Ion Channel [3H] CGP-39653 Binding** | | | |
| CGS 19755 | 1.6E-06 M | 7.7E-07 M | 1.0 |
| **GR *(h)* (agonist radioligand)** | | | |
| dexamethasone | 3.8E-09 M | 1.9E-09 M | 0.9 |

| Compound I.D. | $IC_{50}$(M) | $K_i$(M) | nH |
|---|---|---|---|
| **$H_1$*(h)* (antagonist radioligand)** | | | |
| pyrilamine | 1.9E-09 M | 1.2E-09 M | 1.1 |
| **$H_2$*(h)* (antagonist radioligand)** | | | |
| cimetidine | 5.5E-07 M | 5.4E-07 M | 1.0 |
| **kappa *(h)* (KOP) (agonist radioligand)** | | | |
| U50488 | 6.9E-10 M | 3.8E-10 M | 2.1 |
| **$M_1$*(h)* (antagonist radioligand)** | | | |
| pirenzepine | 4.2E-08 M | 3.7E-08 M | 1.0 |
| **$M_2$ *(h)* (antagonist radioligand)** | | | |
| methoctramine | 4.6E-08 M | 3.2E-08 M | 0.9 |
| **$M_3$*(h)* (antagonist radioligand)** | | | |
| 4-DAMP | 1.0E-09 M | 7.3E-10 M | 1.2 |
| **MAO-A (antagonist radioligand)** | | | |
| clorgyline | 2.1E-09 M | 1.2E-09 M | 1.4 |
| **µ (MOP) *(h)* (agonist radioligand)** | | | |
| DAMGO | 9.0E-10 M | 3.7E-10 M | 1.1 |
| **N neuronal alpha4beta2 *(h)* (agonist radioligand)** | | | |
| nicotine | 9.6E-09 M | 3.2E-09 M | 0.9 |
| **norepinephrine transporter*(h)* (antagonist radioligand)** | | | |
| protriptyline | 5.0E-09 M | 3.7E-09 M | 1.3 |
| **Potassium Channel hERG (human)- [3H] Dofetilide** | | | |
| Terfenadine | 6.6E-08 M | 4.6E-08 M | 0.7 |
| **Sodium Channel Site2 (Non-selective) Rat Ion Channel Batrachotoxin Mass Spectrometry Binding** | | | |
| Veratridine | 3.0E-06 M | 1.1E-06 M | 0.7 |
| **$V_{1a}$*(h)* (agonist radioligand)** | | | |
| [d$(CH_2)_5^1$,Tyr$(Me)_2$]-AVP | 5.5E-10 M | 3.4E-10 M | 0.7 |

#### 7.2. *In Vitro* Pharmacology: Enzyme and Uptake Assays

##### 7.2.1. Test Compound Results

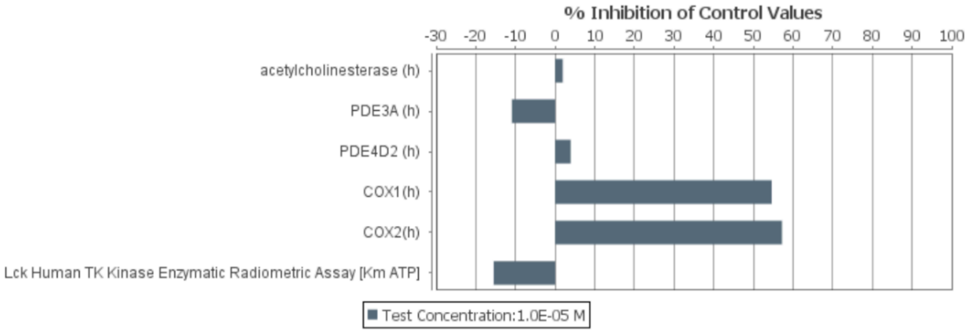

**Figure 2. Histogram for ASAP-0036543**

| Compound I.D. | Client Compound I.D. | Test Concentration | % Inhibition of Control Values 1st | 2nd | Mean |
|---|---|---|---|---|---|
| **acetylcholinesterase *(h)*** | | | | | |
| 100078696-1 | ASAP-0036543 | 1.0E-05 M | 1.5 | 2.2 | 1.9 |
| **PDE3A *(h)*** | | | | | |
| 100078696-1 | ASAP-0036543 | 1.0E-05 M | -8.7 | -13.2 | -10.9 |
| **PDE4D2 *(h)*** | | | | | |
| 100078696-1 | ASAP-0036543 | 1.0E-05 M | 8.9 | -1.2 | 3.9 |
| **COX1(h)** | | | | | |
| 100078696-1 | ASAP-0036543 | 1.0E-05 M | 52.5 | 56.7 | **54.6** |
| **COX2(h)** | | | | | |
| 100078696-1 | ASAP-0036543 | 1.0E-05 M | 54.3 | 60.2 | **57.2** |
| **Lck Human TK Kinase Enzymatic Radiometric Assay [Km ATP]** | | | | | |
| 100078696-1 | ASAP-0036543 | 1.0E-05 M | -18.7 | -12.3 | -15.5 |

##### 7.2.2. Reference Compound Results

| Compound I.D. | $IC_{50}$(M) | nH |
|---|---|---|
| **acetylcholinesterase *(h)*** | | |
| galanthamine | 9.1E-07 M | 1.0 |
| **PDE3A *(h)*** | | |
| milrinone | 4.6E-07 M | 0.9 |
| **PDE4D2 *(h)*** | | |
| Ro 20-1724 | 3.7E-07 M | 0.9 |
| **COX1(h)** | | |
| Diclofenac | 8.4E-09 M | 1.5 |
| **COX2(h)** | | |
| NS398 | 8.5E-08 M | 1.4 |
| **Lck Human TK Kinase Enzymatic Radiometric Assay [Km ATP]** | | |
| Staurosporine | 2.3E-09 M | 1.2 |

### 8. RESULTS INTERPRETATION GUIDE

##### *In Vitro* Pharmacology

Results showing an inhibition (or stimulation for assays run in basal conditions) higher than 50% are considered to represent significant effects of the test compounds. 50% is the most common cut-off value for further investigation (determination of $IC_{50}$ or $EC_{50}$ values from concentration-response curves) that we would recommend.
Results showing an inhibition (or stimulation) between 25% and 50% are indicative of weak to moderate effects (in most assays, they should be confirmed by further testing as they are within a range where more inter-experimental variability can occur).
Results showing an inhibition (or stimulation) lower than 25% are not considered significant and mostly attributable to variability of the signal around the control level.
Low to moderate negative values have no real meaning and are attributable to variability of the signal around the control level. High negative values (≥ 50%) that are sometimes obtained with high concentrations of test compounds are generally attributable to non-specific effects of the test compounds in the assays. On rare occasion they could suggest an allosteric effect of the test compound.

### 9. MATERIALS AND METHODS

#### 9.1. Experimental Conditions

Minor variations to the experimental protocol described below may have occurred during the testing, they have no impact on the quality of the results obtained.

##### 9.1.1. *In Vitro* Pharmacology: Binding Assays

| Assay | Source | Ligand | Conc. | Kd | Non Specific | Incubation | Detection Method | Bibl. |
|---|---|---|---|---|---|---|---|---|
| **Receptors** | | | | | | | | |
| **$5\text{-HT}_{1A}$ *(h)* (agonist radioligand)** | human recombinant (HEK-293 cells) | $[^3H]$8-OH-DPAT | 0.5 nM | 0.5 nM | 8-OH-DPAT (10 µM) | 60 min RT | Scintillation counting | 164 |
| **$5\text{-HT}_{1B}$ *(h)* (antagonist radioligand)** | human recombinant (Chem-1 (RBL) cells) | $[^3H]$GR125743 | 1 nM | 0.8 nM | Serotonine (30 µM) | 60 min 37°C | Scintillation counting | 1451 |
| **$5\text{-HT}_{2A}$ *(h)* (agonist radioligand)** | human recombinant (HEK-293 cells) | $[^{125}I]$(±)DOI | 0.1 nM | 0.3 nM | (±)DOI (1 µM) | 60 min RT | Scintillation counting | 288 |
| **$5\text{-HT}_{2B}$ *(h)* (agonist radioligand)** | human recombinant (CHO cells) | $[^{125}I]$(±)DOI | 0.2 nM | 0.2 nM | (±)DOI (1 µM) | 60 min RT | Scintillation counting | 571 |
| **$alpha_{1A}$ *(h)* (antagonist radioligand)** | human recombinant (CHO cells) | $[^3H]$prazosin | 0.1 nM | 0.1 nM | epinephrine (0.1 mM) | 60 min RT | Scintillation counting | 897 |
| **$alpha_{2A}$ *(h)* (antagonist radioligand)** | human recombinant (CHO cells) | $[^3H]$RX 821002 | 1 nM | 0.8 nM | (-)epinephrine (100 µM) | 60 min RT | Scintillation counting | 542 |
| **$A_{2A}$ *(h)* (agonist radioligand)** | human recombinant (HEK-293 cells) | $[^3H]$CGS 21680 | 6 nM | 27 nM | NECA (10 µM) | 120 min RT | Scintillation counting | 141 |
| **AR(h) (agonist radioligand)** | human endogenous (LNCaP cells) | $[^3H]$methyltrienolone | 1 nM | 0.6 nM | testostérone (1 µM) | 4 hr 22°C | Scintillation counting | 498 |
| **$beta_1$ *(h)* (agonist radioligand)** | human recombinant (HEK-293 cells) | $[^3H]$(-)CGP 12177 | 0.3 nM | 0.39 nM | alprenolol (50 µM) | 60 min RT | Scintillation counting | 548 |
| **$beta_2$ *(h)* (antagonist radioligand)** | human recombinant (CHO cells) | $[^3H]$(-)CGP 12177 | 0.3 nM | 0.15 nM | alprenolol (50 µM) | 120 min RT | Scintillation counting | 794 |
| **BZD (central) (h) (agonist radioligand)** | human recombinant (CHO cells) | [3H] Flunitrazepam | 1 nM | 7.0 nM | Diazepam (10 µM) | 120 min 4°C | Scintillation counting | 1589 |
| **$CB_1$ *(h)* (agonist radioligand)** | human recombinant (Chem-RBL cells) | $[^3H]$CP 55940 | 2 nM | 0.9 nM | AM281 (10 µM) | 30 min 22°C | Scintillation counting | 657 |
| **$CB_2$ *(h)* (agonist radioligand)** | human recombinant (CHO cells) | $[^3H]$WIN 55212-2 | 0.8 nM | 1.5 nM | WIN 55212-2 (5 µM) | 120 min 37°C | Scintillation counting | 165 |
| **$CCK_1$ ($CCK_A$) *(h)* (agonist radioligand)** | human recombinant (CHO cells) | $[^{125}I]$CCK-8s | 0.08 nM | 0.24 nM | CCK-8s (1 µM) | 60 min RT | Scintillation counting | 562 |

| Assay | Source | Ligand | Conc. | Kd | Non Specific | Incubation | Detection Method | Bibl. |
|---|---|---|---|---|---|---|---|---|
| **$D_1$ *(h)* (antagonist radioligand)** | human recombinant (CHO cells) | [$^3$H]SCH 23390 | 0.3 nM | 0.2 nM | SCH 23390 (1 µM) | 60 min RT | Scintillation counting | 281 |
| **$D_{2S}$ *(h)* (agonist radioligand)** | human recombinant (HEK-293 cells) | [$^3$H]7-OH-DPAT | 1 nM | 0.68 nM | butaclamol (10 µM) | 60 min RT | Scintillation counting | 87 |
| **delta (DOP) *(h)* (agonist radioligand)** | human recombinant (Chem-1 (RBL) cells) | [$^3$H]DADLE | 0.5 nM | 0.6 nM | naltrexone (10 µM) | 60 min RT | Scintillation counting | 501 |
| **$ET_A$ *(h)* (agonist radioligand)** | human recombinant (CHO cells) | [$^{125}$I]endothelin-1 | 0.03 nM | 0.03 nM | endothelin-1 (100 nM) | 120 min 37°C | Scintillation counting | 30 |
| **Glutamate (NMDA NR1/ NR2A) Human Ion Channel [3H] CGP-39653 Binding** | human recombinant (HEK-293 cells) from B'SYS | [3H] CGP-39653 | 30 nM | 28 | CGS 19755 (100 µM) | 120 minutes à 4 °C | Scintillation counting | 221 |
| **GR *(h)* (agonist radioligand)** | human endogenous (IM-9 cells) | [$^3$H]dexamethasone | 1.5 nM | 1.5 nM | triamcinolone (10 µM) | 24 hr 4°C | Scintillation counting | 283 |
| **$H_1$ *(h)* (antagonist radioligand)** | human recombinant (HEK-293 cells) | [$^3$H]pyrilamine | 1 nM | 1.7 nM | pyrilamine (1 µM) | 60 min RT | Scintillation counting | 492 |
| **$H_2$ *(h)* (antagonist radioligand)** | human recombinant (CHO cells) | [$^{125}$I]APT | 0.075 nM | 2.9 nM | tiotidine (100 µM) | 120 min RT | Scintillation counting | 540 |
| **kappa *(h)* (KOP) (agonist radioligand)** | human recombinant (RBL cells) | [$^3$H]U69593 | 0.5 nM | 0.6 nM | naloxone (10µM) | 60 min RT | Scintillation counting | 222 |
| **$M_1$ *(h)* (antagonist radioligand)** | human recombinant (CHO cells) | [$^3$H]pirenzepine | 2 nM | 13 nM | atropine (1 µM) | 60 min RT | Scintillation counting | 59 |
| **$M_2$ *(h)* (antagonist radioligand)** | human recombinant (CHO cells) | [$^3$H]AF-DX 384 | 2 nM | 4.6 nM | atropine (1 µM) | 60 min RT | Scintillation counting | 59 |
| **$M_3$ *(h)* (antagonist radioligand)** | human recombinant (CHO cells) | [$^3$H]4-DAMP | 0.2 nM | 0.5 nM | atropine (1 µM) | 60 min RT | Scintillation counting | 546 |
| **µ (MOP) *(h)* (agonist radioligand)** | human recombinant (HEK-293 cells) | [$^3$H]DAMGO | 0.5 nM | 0.35 nM | naloxone (10 µM) | 120 min RT | Scintillation counting | 260 |
| **N neuronal alpha4beta2 *(h)* (agonist radioligand)** | human recombinant (SH-SY5Y cells) | [$^3$H]cytisine | 0.6 nM | 0.3 nM | nicotine (10 µM) | 120 min 4°C | Scintillation counting | 1084 |
| **$V_{1a}$ *(h)* (agonist radioligand)** | human recombinant (CHO cells) | [$^3$H]AVP | 0.3 nM | 0.5 nM | AVP (1 µM) | 60 min RT | Scintillation counting | 343 |
| **Ion channels** | | | | | | | | |
| **5-$HT_3$ *(h)* (antagonist radioligand)** | human recombinant (CHO cells) | [$^3$H]BRL 43694 | 0.5 nM | 1.15 nM | MDL 72222 (10 µM) | 120 min RT | Scintillation counting | 109 |
| **$K_V$ channel (antagonist radioligand)** | rat cerebral cortex | [$^{125}$I]α-dendrotoxin | 0.01 nM | 0.04 nM | α-dendrotoxin (50 nM) | 60 min RT | Scintillation counting | 225 |

| Assay | Source | Ligand | Conc. | Kd | Non Specific | Incubation | Detection Method | Bibl. |
|---|---|---|---|---|---|---|---|---|
| **Cav1.2 (L-type) Human Calcium Ion Channel Binding (Dihydropyridine Site) LeadHunter Assay - FR** | human recombinant (CHO cells) | [$^3$H]nitrendipine | 1.5 nM | 0.7 nM | Nitrendipine (1µM) | 120 minutes RT | Scintillation counting | 1659 |
| **Potassium Channel hERG (human)- [3H] Dofetilide** | human recombinant (HEK-293 cells) | [$^3$H]Dofetilide | 3 nM | 6.6 nM | Terfenadine (25 µM) | 60 min RT | Scintillation counting | 1398 |
| **Sodium Channel Site2 (Non-selective) Rat Ion Channel Batrachotoxin Mass Spectrometry Binding** | rat brain | Batrachotoxin | 15 nM | 8.9 nM | Veratridine (1 mM) | 60 minutes at 37°C | MS | 28 |
| **Transporters** | | | | | | | | |
| **5-HT transporter *(h)* (antagonist radioligand)** | human recombinant (CHO cells) | [$^3$H]imipramine | 2 nM | 1.7 nM | imipramine (10 µM) | 60 min RT | Scintillation counting | 566 |
| **dopamine transporter *(h)* (antagonist radioligand)** | human recombinant (CHO cells) | [$^3$H]BTCP | 4 nM | 4.5 nM | BTCP (10 µM) | 120 min 4°C | Scintillation counting | 190 |
| **norepinephrine transporter *(h)* (antagonist radioligand)** | human recombinant (CHO cells) | [$^3$H]nisoxetine | 1 nM | 2.9 nM | desipramine (1 µM) | 120 min 4°C | Scintillation counting | 180 |
| **Other enzymes** | | | | | | | | |
| **MAO-A (antagonist radioligand)** | rat cerebral cortex | [$^3$H]Ro 41-1049 | 10 nM | 14 nM | clorgyline (1 µM) | 60 min 37°C | Scintillation counting | 36 |

#### 9.1.2. *In Vitro* Pharmacology: Enzyme and Uptake Assays

| Assay | Source | Substrate/ Stimulus/Tracer | Incubation | Measured Component | Detection Method | Bibl. |
|---|---|---|---|---|---|---|
| **Kinases** | | | | | | |
| **Lck Human TK Kinase Enzymatic Radiometric Assay [Km ATP]** | human recombinant (insect cells) | 33P | 40 min RT | ATP (90 µM) + KVEKIGEGTYGVVYK Cdc2 peptide (250 µM) | Scintillation counting | 1645, 1646 |
| **Other enzymes** | | | | | | |
| **acetylcholinesterase *(h)*** | human recombinant (HEK-293 cells) | Acetylthiocholine (400 µM) | 30 min RT | 5 thio 2 nitrobenzoic acid | Photometry | 63 |
| **PDE3A *(h)*** | human recombinant (Sf21 cells) | [$^3$H]cAMP + cAMP (0.5µM) | 15 min RT | [3H]5'AMP | Scintillation counting | 1399 |
| **PDE4D2 *(h)*** | human recombinant (Sf9 cells) | [$^3$H]cAMP + cAMP (0.5µM) | 20 min RT | [3H]5'AMP | Scintillation counting | 1399 |
| **COX1(h)** | human recombinant | Arachidonic acid (3µM) + ADHP ( 25 µM) | 3 min RT | Resorufin (oxydized ADHP) | Fluorimetry | 1480 |
| **COX2(h)** | human recombinant (Sf9 cells) | arachidonic acid (1.2 µM)+ ADHP (25 µM) | 5 min RT | Resorufin (oxydized ADHP) | Fluorimetry | 1480 |

#### 9.2. Analysis and expression of results

##### 9.2.1. *In Vitro* Pharmacology: Binding Assays

The results are expressed as a percent of control specific binding

$$\frac{\text{measured specific binding}}{\text{control specific binding}} * 100$$

and as a percent inhibition of control specific binding

$$100 - \left(\frac{\text{measured specific binding}}{\text{control specific binding}} * 100\right)$$

obtained in the presence of ASAP-0036543.

The $IC_{50}$ values (concentration causing a half-maximal inhibition of control specific binding) and Hill coefficients (nH) were determined by non-linear regression analysis of the competition curves generated with mean replicate values using Hill equation curve fitting

$$Y = D + \left[\frac{A - D}{1 + (C/C_{50})^{nH}}\right]$$

where Y = specific binding, A = left asymptote of the curve, D = right asymptote of the curve, C = compound concentration, $C_{50}$ = $IC_{50}$, and nH = slope factor. This analysis was performed using software developed at Cerep (Hill software) and validated by comparison with data generated by the commercial software SigmaPlot® 4.0 for Windows® (© 1997 by SPSS Inc.).
The inhibition constants ($K_i$ ) were calculated using the Cheng Prusoff equation

$$K_i = \frac{IC_{50}}{(1 + L/K_D)}$$

where L = concentration of ligand in the assay, and $K_D$ = affinity of the ligand for the receptor.

##### 9.2.2. *In Vitro* Pharmacology: Enzyme and Uptake Assays

The results are expressed as a percent of control specific activity

$$\frac{\text{measured specific activity}}{\text{control specific activity}} * 100$$

and as a percent inhibition of control specific activity

$$100 - \left(\frac{\text{measured specific activity}}{\text{control specific activity}} * 100\right)$$

obtained in the presence of ASAP-0036543.

The $IC_{50}$ values (concentration causing a half-maximal inhibition of control specific activity), $EC_{50}$ values (concentration producing a half-maximal increase in control basal activity), and Hill coefficients (nH) were determined by non-linear regression analysis of the inhibition/concentration-response curves generated with mean replicate values using Hill equation curve fitting

$$Y=D+\left[\frac{A-D}{1+(C/C_{50})^{nH}}\right]$$

where Y = specific activity, A = left asymptote of the curve, D = right asymptote of the curve, C = compound concentration, $C_{50}$ = $IC_{50}$ or $EC_{50}$ , and nH = slope factor.
This analysis was performed using software developed at Cerep (Hill software) and validated by comparison with data generated by the commercial software SigmaPlot® 4.0 for Windows® (© 1997 by SPSS Inc.).
